# A statistical censoring approach accounts for hook competition in abundance indices from longline surveys

**DOI:** 10.1101/2022.07.14.500091

**Authors:** Joe Watson, Andrew M. Edwards, Marie Auger-Méthé

## Abstract

Fishery-independent longline surveys provide valuable data to monitor fish populations. However, competition for bait on the finite number of hooks leads to biased estimates of relative abundance when using simple catch-per-unit-effort methods. Numerous bias-correcting instantaneous-catch-rate methods have been proposed, modelling the bait removal times as independent random variables. However, experiments have cast doubts on the many assumptions required for these to accurately infer relative abundance. We develop a new approach by treating some observations as right-censored, acknowledging that observed catch counts are lower bounds of what they would have been in the absence of hook competition. Through simulation experiments we confirm that our approach consistently outperforms previous methods. We demonstrate performance of all methods on longline survey data of eleven species. Accounting for hook competition leads to large differences in relative indices (often −50% to +100%), with effects of hook competition varying between species (unlike other methods). Our method can be applied using existing statistical packages and can include environmental influences, making it a general and reliable method for analyzing longline survey data.

## 1 Introduction

Monitoring changes in population abundance is essential for the successful management of fisheries (Hilborn and Walters, 2013; Kuriyama *et al.*, 2019). Estimates of abundance trends are frequently made using data collected from fishery-independent surveys and these estimates are subsequently used as inputs for formal stock assessments to advise fisheries management on harvest controls (Gunderson, 1993; Hilborn and Walters, 2013). Longline surveys play a crucial role in the management of many demersal fish species, such as rockfish species that frequently inhabit rocky and untrawlable habitats (Rodgveller *et al.*, 2011; Obradovich, 2018). For longline surveys, the commonly-used catch-per-unit-effort (CPUE) method typically considers catch counts scaled by the number of baited hooks deployed (e.g., Anderson *et al.* 2019). Additional factors such as hook spacing, hook type, and soak time may also need to be considered (Yamanaka *et al.*, 2008). However, using CPUE to estimate the local abundances of fish species is associated with numerous challenges (Obradovich, 2018).

The challenges with longline fishing data originate from processes that sequentially occur between the deployment of longline gear and the subsequent capture of each fish. In particular, captured individuals must be (1) close to the baited hooks, (2) sufficiently attracted by the bait odour to move towards the hooks (Løkkeborg *et al.*, 1995; Sigler, 2000), (3) able to reach a baited hook before it is taken by another animal and before the end of the soak time (Rothschild, 1967; Somerton and Kikkawa, 1995), (4) unable to escape capture by the hook, and (5) able to avoid predation by scavengers for the remainder of the soak time (Ward *et al.*, 2004). Each of these processes can vary between deployments. For example, bait attractiveness is linked to bait type and size (Løkkeborg and Bjordal, 1995), hunger levels of the fish (Løkkeborg *et al.*, 1995), temperature (Stoner *et al.*, 2006), availability of natural prey (Stoner, 2004), light intensity (Stoner, 2003), time of day (Fernö *et al.*, 1986; Ward *et al.*, 2004), and currents (Fernö *et al.*, 1986). Therefore, to link changes in observed catch counts from longline surveys to true changes in the abundance of a species, one must account for the variability due to heterogeneous bait attractiveness, heterogeneous soak times, hook escape, the removal of captured fish by scavengers, interspecific and intraspecific competition for finite hooks, and gear saturation (Sigler, 2000). Although carefully designed longline surveys with standardized sampling protocols (e.g. fixed gear and bait types used from year to year) can help control for some of these factors (Gunderson, 1993; Hilborn and Walters, 2013), survey design alone cannot remove the biasing effects that hook competition and gear saturation have on current estimators of abundance (Rothschild, 1967; Ricker, 1975; Somerton and Kikkawa, 1995; Kuriyama *et al.*, 2019).

A desirable method for estimating relative abundance should simultaneously have two fundamental properties. First, when the true abundance is stationary, the long-run frequency in which the method falsely infers a change in abundance should be controllable by the fisheries scientist (a controllable type I error rate, hereafter referred to as the type I property). Second, it should identify, with high probability, a change in abundance when it occurs, and accurately estimate the magnitude of the change (related to the type II error rate and thus called the type II property hereafter).

Theoretical examples demonstrate why unadjusted CPUE indices of relative abundance exhibit neither the type I nor the type II properties. For the type I property, consider a standardized annual longline survey that catches species A and species B. For all fishing events, suppose that the same number of baited hooks are deployed and that all baits are taken by the end of each soak time. Both species are caught throughout the soak times of the fishing events. If the absolute abundance of species A is constant from year to year but species B increases, the expected CPUE of species A will decrease with time (violating the type I property), due to the increased competition from species B for the finite hooks during the fishing events in the later years. Indeed, the longline CPUE of a species can be highly sensitive to the density of competing species (Rothschild, 1967). The magnitude of the expected decline in CPUE of a species will depend upon its ability to locate baited hooks ahead of the competing species. Numerous biological characteristics, such as its aggressiveness and olfactory sensitivity, will determine this ability (Sigler, 2000).

For the type II property, consider a standardized longline survey over two years for which only one species is caught. Ignoring variability for simplicity, suppose each fishing event deploys 1000 baited hooks and catches 800 fish in year 1. Suppose the abundance of the species doubles in year 2. Despite the true abundance having increased by 100%, the limited number of baited hooks available to the fish prevents the estimated abundance from an unadjusted CPUE index from increasing by more than 20%. This is called gear saturation and as a consequence, relative abundance estimates in year 2 will be negatively biased (less than the true change in abundance).

The processes leading to the negative biases in the above two examples are called hook competition and gear saturation. To simplify exposition, we hereafter refer to these processes as hook competition. Not all baits need to be removed during a fishing event for biases due to hook competition to creep into CPUE estimates of relative abundance. The ability of an attracted fish to locate a baited hook may decrease as baits are removed from the fishing gear, possibly once a threshold proportion of baits has been removed (Sigler, 2000). The threshold is expected to be species-specific and depend on numerous biological factors of the fish species (Sigler, 2000). Thus, CPUE estimates of relative abundance may poorly correlate with the true species’ abundance in regions where total fish density is high (relative to the number of baited hooks deployed) and for species that travel in large groups. This result has been determined through mathematical derivation (Rothschild, 1967; Somerton and Kikkawa, 1995), observed in simulation studies (Kuriyama *et al.*, 2019), observed in experimental settings with observations taken from submersible remote operated vehicles (ROVs; Obradovich 2018), and confirmed through the careful analysis of longline survey data (Rodgveller *et al.*, 2008).

Numerous estimators of relative abundance have been put forward with the aim of better controlling for bias due to hook competition (Rothschild, 1967; Ricker, 1975; Somerton and Kikkawa, 1995). All of these methods focus on modelling the fate of each baited hook present in a longline fishing event. In particular, by deriving a system of fixed-rate ordinary differential equations (ODEs) that closely follows the intuition behind the Baranov catch equation, the bait removal times from each species are modelled (Gulland, 1955). The target of inference for each species becomes their fixed instantaneous catch rate (ICR) parameter, defined as their expected rate of bait removal from a hook per unit time.

Various statistical distributions have been proposed to account for the random variability observed in the data that is not explained by the system of ODEs. These include models for the bait removal times when hook timer data are available, and models for binary catch/no-catch events, useful in standard longline fishing scenarios (see Obradovich, 2018, for a detailed discussion). Maximum likelihood and Bayesian approaches have been derived and methods have been developed to control for correlations between hooks on the same longline skate and to adjust for hook-level covariates (Somerton and Kikkawa, 1995; Obradovich, 2018). Finally, a link between a species’ ICR and its underlying abundance is made by assuming a functional relationship between the ICR and its abundance, often assumed to be linear (Obradovich, 2018).

Three assumptions are crucial for ICR-based abundance indices to successfully describe the relative abundance of a species through time (Obradovich, 2018), but are generally not met in practice. First, the ICR is assumed to be constant throughout the soak time of a fishing event. However, studies where the exact bait removal times were recorded (e.g. using hook-timers or ROVs) demonstrated time-varying rates in numerous species, including Sablefish (*Anoplopoma fimbria*; Løkkeborg *et al.*, 1995; Sigler, 2000), Pelagic Armorhead (*Pentaceros richardsoni*; Somerton and Kikkawa, 1995), Yelloweye Rockfish (*Sebastes ruberrimus*) and Quillback Rockfish (*S. maliger*; Obradovich, 2018). Second, the expected catch count is assumed to be linearly proportional to the number of hooks (required due to the assumed independence between the bait-removal events). However, experiments have demonstrated that increasing the number of hooks on a longline significantly decreases the average catch per hook for both Pacific Halibut (Skud and Hamley, 1978) and Sablefish (Sigler, 2000). Third, the ICR is assumed to be linearly proportional to the true species’ abundance locally around the longline gear. However, results from experiments that compared direct estimates of abundance from ROV-mounted cameras with ICR-based abundance indices have been inconsistent with the assumption of linearity (Rodgveller *et al.*, 2011; Obradovich, 2018).

Under the above three assumptions, and assuming that bait removal times are exponentially-distributed, the maximum likelihood estimator of the relative abundance equals the CPUE value multiplied by a nonlinear scale factor that increases with the total proportion of baits removed by all sources. The scale factor attempts to account for all the bait removals from fish of the target species that were ‘lost to observation’ due to hook competition by enumerating the expected number of ‘missed’ catches. Crucially, the scale factor inflates the ICR estimator of relative abundance away from the CPUE value by an amount that increases with the total catch of non-target species. This ICR estimator has been proposed as a corrective offset term within a modelling framework (Webster *et al.*, 2011). We refer to this method as the ICR method.

When species’ true ICRs are time-varying within the soak time, the ICR method is biased and fails to attain the type I property. To demonstrate why, suppose standardized longline fishing gear with 1000 hooks is deployed in each of two years. Let there be two species present, with 990 fish of species A caught each year and 0 and 9 fish of species B caught in years 1 and 2 respectively. Suppose no hook escape or mechanical bait loss occurs and consider time-varying ICR’s whereby species A always reaches a bait ahead of species B during a soak time. Given such perfect knowledge, we would not infer a change in species A’s abundance from year 1 to year 2. However, from the catch data an ICR method’s estimate of relative abundance assuming time-constant ICRs would estimate species A’s abundance to increase by 50% (see Supplementary Material).

These demonstrations of the CPUE and ICR methods failing to attain the type I and type II properties motivate us to derive a new censored method whose performance is robust. The censoring effectively treats a subset of the observed catch counts (deemed to have been affected by hook competition) as lower bounds of what the catch counts would have been in the absence of hook competition. The prevalence and effect of hook competition are dependent on the species of interest. We conduct extensive simulation experiments covering multiple realistic data-generating scenarios and find that: i) the CPUE and ICR methods fail to satisfy the type I property in many realistic settings; ii) our proposed method satisfies the type I and type II properties in most settings. We apply our method to real data collected for multiple species on an annual longline survey. We estimate relative abundance indices (such as are used as inputs to formal stock assessment models), finding large differences between indices from the CPUE and ICR methods and from our new method (often by −50% to 100%, sometimes up to 150%). Thus, we recommend use of our new method in future applications.

For reproducibility we supply all simulation and case study R (R Core Team, 2022) code and data files at https://github.com/joenomiddlename/Censored_Longline_RCode. We provide short template code for implementing our method using each of three commonly-used R packages: R-INLA (Rue *et al.*, 2009; Lindgren *et al.*, 2011; Lindgren and Rue, 2015), TMB (Kristensen *et al.*, 2016) via sdmTMB (Anderson *et al.*, 2022), and Stan (Carpenter *et al.*, 2017) via brms (Bürkner, 2017). The R-INLA and sdmTMB R packages were updated specifically to implement our censored method.

## 2 Materials and methods

To develop our new modelling approach, we first build a flexible process model to describe the spatial density of fish. Next, we develop an observer model for linking observed catch counts collected using longline gear to the process model in an idealized setting without hook competition. We then modify the resulting likelihood function by treating hook competition as a data-quality issue. The catch counts from fishing events experiencing sufficiently high bait-removal levels are treated as right-censored (i.e. treated as lower bounds to the truth). This allows us to account for hook competition without needing to specify its precise nature or magnitude. We finish the section with descriptions of the simulation experiments and the case study using real data.

### 2.1 Mechanism describing the densities of fish

Let **Ω** denote our spatial region, **s** a point in the region, *ω* a subregion within **Ω**, and *t* = 1,2,…,*T* denote years (notation is summarized in Table 1). The density *λ_it_* (**s**) defines the expected number of individuals of species *i* found per unit area of the seabed around location **s** at an arbitrary moment in year *t*. As is frequently done in the fisheries literature (Thorson *et al.*, 2015, 2016; Grüss and Thorson, 2019), we use the following general model:

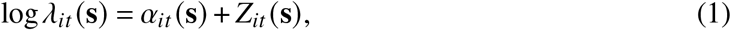

where *α_it_* (**s**) captures the overall effect of measured covariates (such as temperature and other environmental influences) affecting the population density across space and time, and (**s**) is a stochastic term that captures the overall effect of unmeasured (perhaps unknown) sources of vari-ability. Specifically, *Z_it_* (**s**) captures residual spatial, temporal, and spatio-temporal correlations due to, for example, unmeasured changes in predator abundance.

**Table 1:**
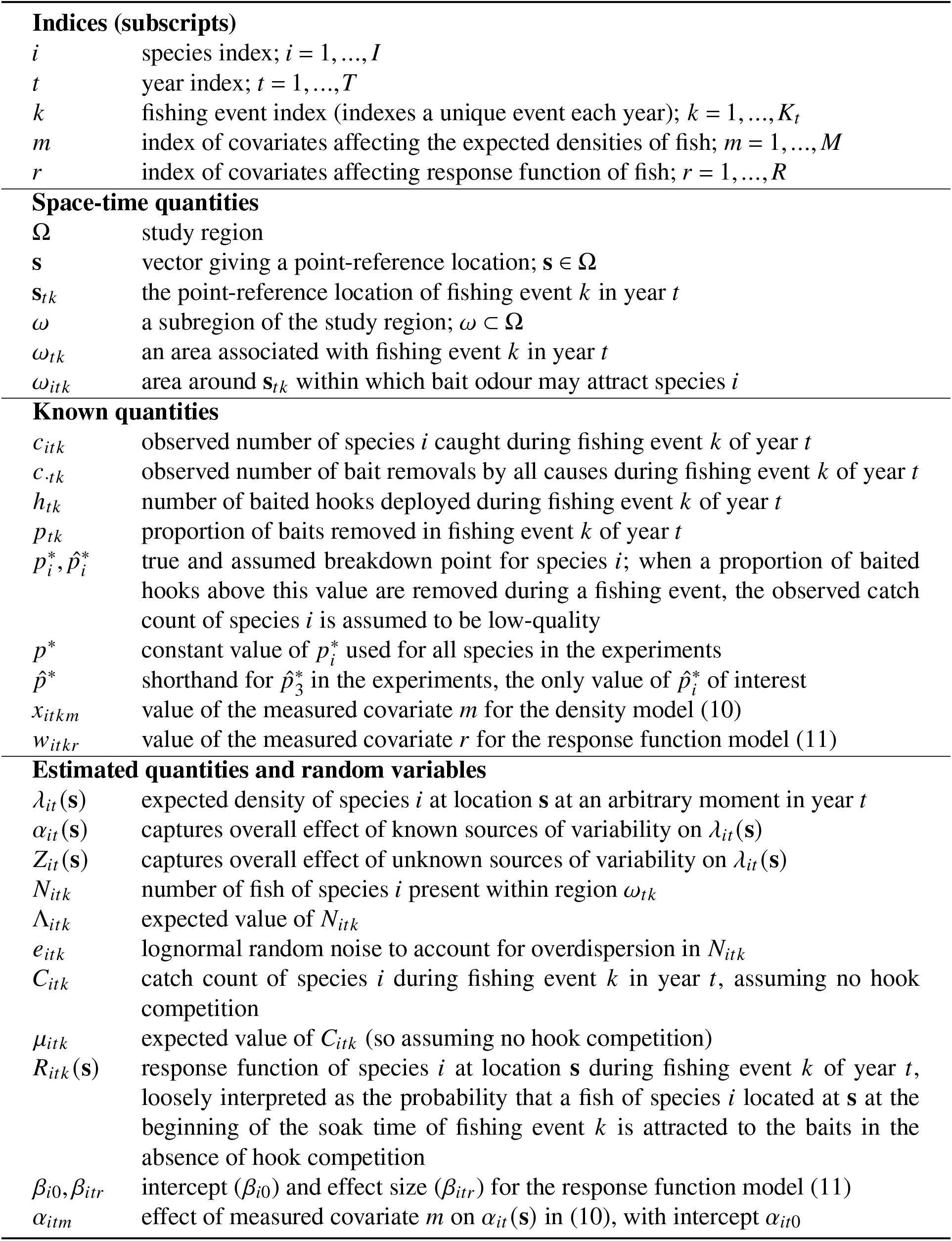
List of main notation used.

Next, consider *K_t_* fishing events in year *t*, indexed by *k* = 1,2,…,*K_t_*, with associated spatial regions *ω_tk_*. Define random variable *N_itk_* as the number of fish of species *i* present within region *ω_tk_* during the fishing event *k* in year *t,* with mean, *Λ_itk_*, therefore given by

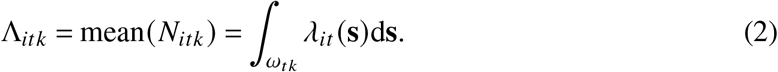

A statistical model must now be specified on *N_itk_* and *Z_it_* (**s**), with *N_itk_* conditional upon *Λ_itk_* and *Z_it_* (**s**). We assume the *Nnk* are conditionally independent across the fishing events *k* (i.e. local depletion due to the fishing event does not occur).

The simplest choice of conditional count distribution is the Poisson distribution. For species that exhibit schooling behaviour, the variance of their catch counts *N_itk_* may be far higher than expected from a Poisson distribution (and the signal-to-noise ratio far lower) due to increased overdispersion and/or right-skewness. To account for both, a popular choice in ecology is to scale *Λ_itk_* by an independent identically distributed (IID) lognormal white noise term, which we denote *e_itk_*, that is unique for every fishing event and species (Bulmer, 1974; Engen *et al.*, 2002). Incorporating this term is equivalent to adding IID normal random effects to (1). Then *N_itk_* is conditionally distributed as:

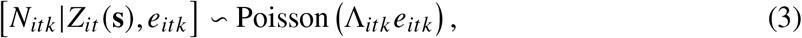

where the square brackets denote ‘the probability distribution of’ and | is ‘conditional upon’.

Finally, we are often interested in an index of abundance of species *i* over the whole region **Ω** for years *t* = 1,2,…,*T*. This index can be calculated as

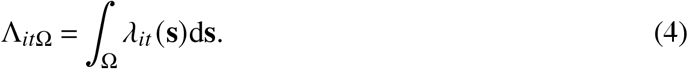

Since longline fishing events catch an unknown fraction of the fish present around the longline gear, absolute values of (4) will be unidentifiable. Thus, our inferential target will be relative values of (4).

### 2.2 Observation model that accounts for fishing but not hook competition

To account for the longline fishing process, we link the underlying fish density model to a novel observation process. Fishing event *k* of year *t* deploys *h_tk_* baited hooks at location *s_tk_*. Around **s**_*tk*_ is believed to be a species-specific region, *ω_itk_*, where the bait odour is sufficiently high that a fish species *i* located within *ω_uk_* may be stimulated to seek a baited hook during the soak time.

Let *R_itk_* (**s**) ∈ [0,1] denote a response function, loosely representing the probability that an individual of species *i* present at location **s** at the beginning of the soak time of longline event *k* in year *t* will seek and remove a bait (and be caught by the gear) in the absence of any other fish or other sources of bait removal; *R_itk_* (**s**) = 0 for **s** outside *ω_itk_*.

Assume the area of *ω_itk_*, denoted |*ω_itk_*|, is sufficiently small such that throughout *ω_itk_* the species density *λ_it_* (**s**) can be assumed constant, such that *λ_it_* (**s**) = *λ_it_* (**s**_*tk*_). Then, from (2) we have *λ_it_*(**s**_*tk*_) = Λ_*itk*_/|*ω_itk_*|. Next, we can model the observed longline catch count of species *i* from longline fishing event *k* in year *t* in the *absence of hook competition,* denoted *C_itk_*, as

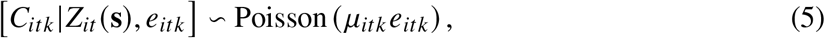

where the expected catch count is

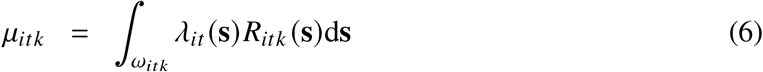

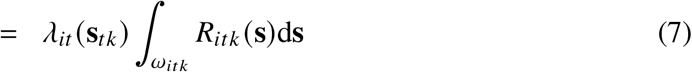

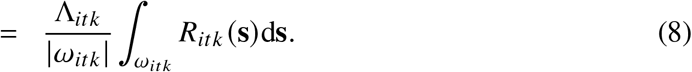

If every fish of species *i* within *ω_itk_* is caught, then we have *R_itk_*(**s**) = 1 within *ω_itk_*, such that *μ_itk_* = Λ_*itk*_ and (5) reduces to model (3) with *N_itk_* = *C_itk_*; i.e. if every fish is caught then the number of fish within the region equals the number caught within the region. The function *R_itk_* (**s**) is known as a thinning function, with the resulting counts arising from a thinned point process (Watson *et al.*, 2021). The function *R_itk_*(**s**) is therefore similar to catchability.

We assume that the true response function of species *i* does not change during the soak time of the fishing event as baits are removed from hooks and for now assume that there are enough baited hooks available for each attracted fish of species *i* (i.e. no hook competition). The function *R_itk_* (**s**) can depend on covariates, including various biotic and abiotic factors affecting the fish behaviours or the characteristics of the odour plumes (Sigler, 2000), such as average hunger levels (Rodgveller *et al.*, 2008), numbers of baited hooks deployed and soak times (Sigler, 2000), and currents (Fernö *et al.*, 1986).

### 2.3 Extending the observation model to account for hook competition

To account for hook competition, we assume that fishing events with high proportions of baits removed are likely to suffer from hook competition, and label them as ‘low-quality’. We replace the catch counts from all low-quality fishing events with inequalities that imply the ‘true’ observed catch counts could have been greater than the actual observed ones if hook competition was absent (some fish that would have been caught were not caught due to hook competition). In statistical terms, we are treating all low-quality fishing events as right-censored.

Specifically, we define the quantity 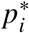 as the minimum proportion of the *h_t_k* deployed baits that need to be taken during a fishing event for us to deem it low-quality for species *i.* For simplicity, 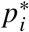 is allowed to be unique for each species but is fixed across *t* and *k*. The term 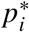 denotes a breakdown point: a point after which the assumed model (5) and/or its assumptions no longer adequately describe the observed catch counts. Hook competition strictly *reduces* the chances of attracted fish of species *i* from being caught once 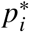 is exceeded. So the observed catch counts (which are affected by hook competition) will be underestimates of the *C_itk_* from (5) which does not consider hook competition. We do not need to know the causes of the bait removals that led to the breakdown (e.g. they can be due to combinations of fish, invertebrates, and mechanical failure), the reasons behind the decrease in species *i*’s ability to locate a bait (e.g. they can be due to changes in the bait odour plume or avoidance of captured fish), nor the magnitude of divergence between model (5) and the expected catch count. Values of 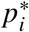 are not estimable within the model and are assigned values 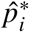; a method for assigning 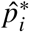 is later developed, tested, and used in the case study.

For all high-quality fishing events, we assume that we perfectly observe the *C_itk_* described by our assumed conditional Poisson model (5). Conversely, for low-quality fishing events, we assume that the observed catch counts of species *i* are right-censored and hence we only observe lower bounds on *C_itk_*. Denoting the observed catch counts as *C_itk_*, the high-quality observations can be written as events *C_itk_ = C_itk_* and the low-quality observations as *C_itk_* ≥ *C_itk_*. Right-censoring reflects the failure of the measuring device (longline gear) to measure the local abundance of species *i* when hook competition is present.

To model the right-censoring, we keep track of the number of baits removed in each fishing event. Let *c_.tk_* = ∑_*i*_ *c_itk_* denote the known number of deployed baits removed in fishing event *k* of year *t* by all species, with *p_tk_* = *c_.itk_/h_tk_* denoting the corresponding proportion of baits removed (and so 1 – *p_tk_* is the proportion of hooks returned with bait). To account for the empty hooks (i.e. hooks where the baits were removed during the soak time, but retrieved without fish attached) we consider them as a unique species. Then, the conditional likelihood function for the observed catch count of species *i* in fishing event *k* of year *t* is

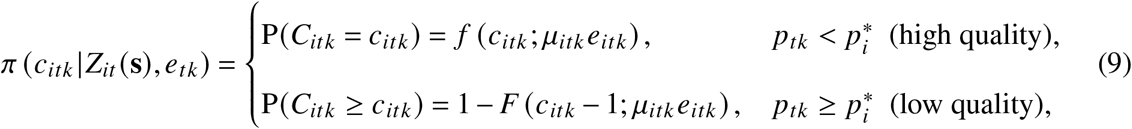

with *f*(*x*;*μ*) denoting the probability mass function, P(*X* = *x*), and *F*(*x*; *μ*) the cumulative distribution function, P(*X* ≤ *x*), of a Poisson random variable *X* with mean parameter *μ.* Note that the likelihood function for censored zeros is exactly one (because *c_itk_* = 0 and *P*(*C_itk_* ≥ 0) = 1 by definition) and so contributes no information to the overall likelihood. Thus, low-quality zero catch counts are effectively discarded.

In settings with very high levels of censorship and limited data, the above model may not converge numerically. To improve convergence in such situations, we can define a species-specific upper bound *u_itk_* and consider an interval-censored event by modifying the second line of (9) to P(*c_itk_* ≤ *C_itk_* ≤ *u_itk_*) = *F*(*u_itk_*;*μ_itk_*) - *F*(*c_itk_* – 1;*μ_itk_e_itk_*). In addition, we can also set the largest observed catch counts each year as ‘high-quality’ (9), regardless of the values of *p_tk_*. We consider both approaches in the Supplementary Material, however, convergence issues appear to be rare if the model-fitting algorithm is started with initial values selected from a converged CPUE-based model with equivalent specification (considering all data as high-quality and using (9)). Note that with or without these modifications, the response variables for all censored (low quality) observations are changed from (scalar) catch counts *c_itk_* to intervals.

### 2.4 Fitting the models

The following choices of model components enable the above framework to be fit using existing software. First, define a linear model for *α_it_*(**s**_*k*_):

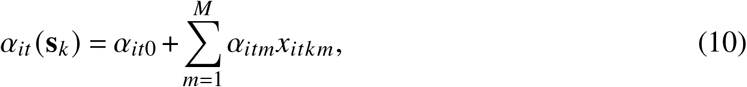

where *α*_*it*0_ is an intercept term and *α_itm_* denotes the effect of measured covariate *m* (where *m* = 1,2,…,*M*) that takes value *x_itkm_*, on the log mean density of species *i* around the fishing gear of event *k* in year *t*. The inclusion of measured covariates is optional, and if none are available then (10) reduces to *α_it_*(**s**_*k*_) = *α*_*it*0_.

Next, define the following log-linear model for the response function:

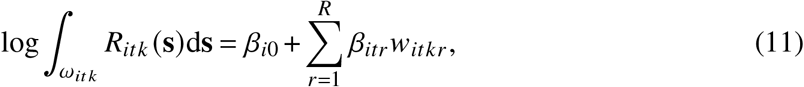

where *β*_*i*0_ is an intercept term and *β_itr_* denotes the effect of covariate *r* (where *r* = 1,2,…,*R*) that takes value *w_itkr_*, on the log of the integrated response function of species *i* for fishing event *k* in year *t*.

Thus, taking the log of the expected catch count *μ_itk_* (7) and using (1), (10), and (11) gives:

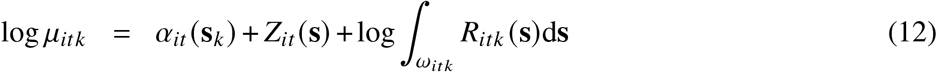

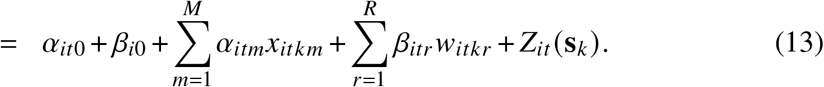

Lastly, specify *Z_it_* (**s**) as a Gaussian process and *e_itk_* as IID lognormal random effects. This leads to the catch counts arising from a log-Gaussian Cox process model (Watson *et al.*, 2021). Then, given observed catch counts *c_itk_* and known proportions of baits removed *p_tk_*, the censored Poisson model defined by equations (9) and (13), can be fit using any preferred software package for fitting generalized linear mixed effects models. Estimates of the number of species *i* over region **Ω** in year *t*, Λ_*it*Ω_, can then be obtained by predicting *λ_it_* (**s**) from (1) across a dense grid of points covering **Ω**, using either maximum likelihood estimates or posterior means in a Bayesian context, and then approximating the spatial integral (8) with finite sums.

Other choices of distributions for *Z_it_* (**s**) and *e_itk_* can be made, but custom code would need to be written. Finally, equation (13) implies that the two intercept terms will not be identifiable from the model. Only their sum will be identifiable. Thus, so long as the response function intercept *β*_*i*0_ does not change over time, and so long as the covariates *x_itkm_* and *w_itkr_* are unique, relative values of **Λ**_*it*Ω_ (up to a constant multiplicative factor) will remain identifiable.

For brevity, we refer to our new method, that involves fitting the censored Poisson-lognormal regression model defined by (9) and (13), as simply the ‘censored method’.

### 2.5 Simulation study

To assess the relative performances of the CPUE and ICR methods against our censored method at inferring relative abundance, we simulate fish populations and data collected from a fishery-independent longline survey using standardized sampling protocols. Longline fishing events are performed at 30 or 100 stations across 6 years, each deploying 800 hooks with soak times of 5 hours. No systematic inter-station variability is present, but there are important temporal trends. We create different experiments that explore a range of potential fish behaviour (e.g., schooling, aggressiveness, correlation in abundance between species), as well as whether only the target species is considered or whether two additional species groups remove the baits: a main competitor whose abundance increases in time, and an ‘other’ species group that includes multiple species (and can include bait loss, etc.). In each experiment, we explore the type I and type II properties by either keeping the abundance of the target species constant or increasing it through time.

Bait-removal events are simulated by: i) sampling numbers of each of the three species groups from their abundance distributions; ii) sampling the bait removal time of each fish from group-specific bite-time distributions; iii) matching fish to the next available baited hook in ascending order of their sampled bite times. Each matched fish-hook pair results in a successful capture and bait-removal event with a capture probability that depends on the group-specific sensitivity to hook competition.

In particular, once the proportion of baits removed exceeds 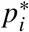, we decrease the probability of capture for species *i.* We then estimate relative abundance indices for only the target species, for which we specify 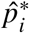 in model fitting. For simplicity, we set the true value 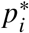 to be the same for all species, however, a second parameter for each species group can change its sensitivity to hook competition. Note that typically 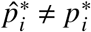.

Assumptions and parameter values are varied to test the performance of the methods under multiple settings. Each assumption-parameter combination is repeated 100 times. The resulting six simulation experiments (which each contain various combinations of settings) are summarized in Table 2 and briefly described when we discuss their results; full details are given in the Supplementary Material, including model fitting details.

**Table 2:** Brief descriptions of the type I and II properties, methods, and simulation experiments.

| **Desirable Properties** |  |
| --- | --- |
| Type I | Correctly infers a stationary abundance when the true abundance is stationary. |
| Type II | Correctly infers a changing abundance when the true abundance is changing and estimates the magnitude of change with high accuracy |
| **Methods** |  |
| CPUE | Catch per unit effort, typically the catch count divided by the number of hooks. For the simulation study and case study, estimators and intervals quantifying uncertainty are derived from Poisson-lognormal regression models. |
| ICR | Instantaneous catch-rate based. Raw CPUE values multiplied by the hook competition scale factor, assuming baits are removed by each species at independent exponentially-distributed times. For the simulation study and case study, estimators and intervals quantifying uncertainty are derived from Poisson-lognormal regression models fit to adjusted CPUE values. |
| Censored | Estimates and intervals of uncertainty derived from Poisson-lognormal regression models. For high-quality fishing events exact CPUE values are used, and for low-quality events inequalities are used (lower bound set to CPUE value). |
| **Experiments** |  |
| 1 | One schooling species is caught, sometimes present in very large groups in numbers far exceeding the number of available hooks (giving a right-skewed catch count distribution). The relative abundance is fixed through time to assess the type I property of the methods, and increased to assess the type II property. Having all baits removed ($p_{tk} = 1$) is commonplace. |
| 2 | Many species are caught (simplified as a target, a main competitor, and other species). The target species is non-schooling and never present in large groups. A main schooling competitor, whose abundance increases through time, is able to reach the baits ahead of the target species on average (under the ‘differing aggressiveness’ setting). The other species group’s abundance remains constant. The relative abundance of the target species is fixed through time to assess the type I property of the methods, and increased to assess the type II property. |
| 3 | The same setup as 2, but the target species’ abundance distribution is overdispersed to represent occasional schooling behaviour, and allowed to be positively and negatively correlated with a competitor’s. Settings are more realistic. |
| 4 | Similar to 2, but the breakdown point $\hat{p}^*$ is very poorly specified to investigate robustness of the censored method to the mis-specification of $\hat{p}^*$. |
| 5 | The same setup as 3, but the target species now reaches the baits (on average) ahead of the competing species. The goal is to investigate the performance of the censored method in settings where it is not expected to perform well because the target species is largely immune to inter-species hook competition. |
| 6 | Regression models are fit to the species’ catch counts to investigate the trend in the mean catch count as the proportion of baits removed during a fishing event increases from $p_{tk} \approx \hat{p}^*$ to $p_{tk} \approx 1$. Multiple data-generating settings are considered. The goal is to identify an empirical approach for determining a suitable $\hat{p}^*$ value. |

The CPUE, ICR, and censored method’s estimators are the posterior medians from fitted Poisson lognormal models. We use identical model specifications to make the comparisons between the estimators fair (e.g. identical linear predictors, priors, software). The only exceptions are the conditional likelihood function and/or ICR scale factor. We assess the performance of each of the methods by comparing their estimation bias, estimation median-squared error (MSE), and observed coverage of their posterior credible intervals. Bias close to zero is desired to demonstrate accuracy. But being an average from simulations, bias is not a guarantee of precision, for which MSE is shown, with low values preferred. Observed coverage is the percentage of the 100 95% credible intervals that cover the known value, with values close to 95% indicating accurate model-based uncertainty quantification. Robust 95% confidence intervals for bias and MSE (where the uncertainty arises from the 100 repeated simulations) are computed using the median absolute deviance from the median values to reduce the influence of outliers (Maronna *et al.*, 2019), while 95% intervals of coverage are calculated using a normal approximation to the binomial probability (binomial because each interval either contains the true value or not).

### 2.6 Case study: International Pacific Halibut Commission fishery-independent setline survey

The International Pacific Halibut Commission (IPHC) fishery-independent setline survey is an annual longline survey spanning waters from California to Alaska, USA (IPHC, 2022). While the survey’s main goal is to provide data on Pacific Halibut (*Hippoglosus stenolepis*) for stock assessment purposes, non-halibut species have been enumerated within Canadian waters off British Columbia since 1995 (see Supplementary Material for map). The survey provides valuable information on many non-halibut species in these waters, and for 32 species it yields the longest ongoing time series from all types of surveys (Anderson *et al.*, 2019). Hook competition is believed to be high, with over 11% of the IPHC fishing events returning zero baited hooks (Anderson *et al.*, 2019), but is not currently accounted for in Canadian analyses which generate inputs to stock assessments used to provide advice to fisheries managers.

We restrict our analysis to data collected on 11 species from fishing events between 1998-2020 at 171 stations within British Columbia waters, 109 of which were fished at every year in the time period. The IPHC data are amalgamated using the R package gfiphc (Edwards *et al.*, 2022).

The stations are located at the intersections of a 10 × 10 nautical mile square grid. At each station, fishing gear consisting of a set of up to eight skates is deployed, with each skate containing around 100 hooks. For each set, the IPHC calculates an effective skate number to account for variations in numbers of hooks, hook spacing, or hook type (Yamanaka *et al.*, 2008). An effective skate of 1 attempts to represent a skate of 100 circle hooks with 18-foot spacing (Yamanaka *et al.*, 2008). Consistent bait are used in every year with the one exception of 2012. We ignore the potential effect of this bait change and consider all catch count data at the set level.

The 11 species are chosen due to their high average catch counts. There are three rockfish species: Redbanded Rockfish (*Sebastes babcocki*), Shortspine Thornyhead (*Sebastolobus alascanus*), and Yelloweye Rockfish; three roundfish species: Lingcod (*Ophiodon elongatus*), Pacific Cod (*Gadus macrocephalus*), and Sablefish; three elasmobranchs: Big Skate (*Beringraja binoculata*), Longnose Skate (*Raja rhina*), and North Pacific Spiny Dogfish (*Squalus suckleyi);* and two flatfish species: Arrowtooth Flounder (*Atheresthes stomias*) and Pacific Halibut.

Current analyses for British Columbia groundfish compute a CPUE value for each fishing event at each set by calculating the number of individuals of that species caught divided by the effective skate of the set (Edwards *et al.*, 2017; Yamanaka *et al.*, 2018; Anderson *et al.*, 2019). Relative abundance indices with 95% confidence intervals are computed by bootstrapping the observed CPUE values within each year (Efron, 1987). Preliminary investigation of hook competition, based on the methods of Clark (2008) and Webster *et al.* (2011), showed it to be important in the data for British Columbia groundfish (Anderson *et al.*, 2019), providing further motivation for the present study.

## 3 Results

### 3.1 Simulation study results

Experiment 1 conducts proof-of-concept trials, simulating longline catch data of a single, highly-abundant species where all baits are commonly removed during fishing events. Results confirm that the censored method is able to infer stationary and nonstationary abundance trends with high accuracy, high precision, and with coverage levels close to the target, even with frequently saturated fishing gear (see Supplementary Material). Results also confirm our earlier claim that the CPUE method fails the type II property when the demand for baited hooks from only a single species frequently exceeds the number supplied by the longline gear. The ICR method performed worse.

Experiment 2 simulates multiple species (Figures 1 and 2). The target species is non-schooling, its distribution is independent of the other species, and its local abundance is less than the number of hooks (800). The main competitor increases in abundance through time with individuals, on average, reaching the baits at the same time as individuals from the target species when all species’ bite times are simulated from uniform distributions, and ahead of the target species when the competitor’s bite times are simulated from an exponential distribution. Thus, hook competition occurs and is sometimes severe. Values of *p*\* equal to 0.85 and 1 are used to generate the data, while 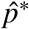 values of 0.85 and 0.95 are assumed throughout.

**Figure 1:**
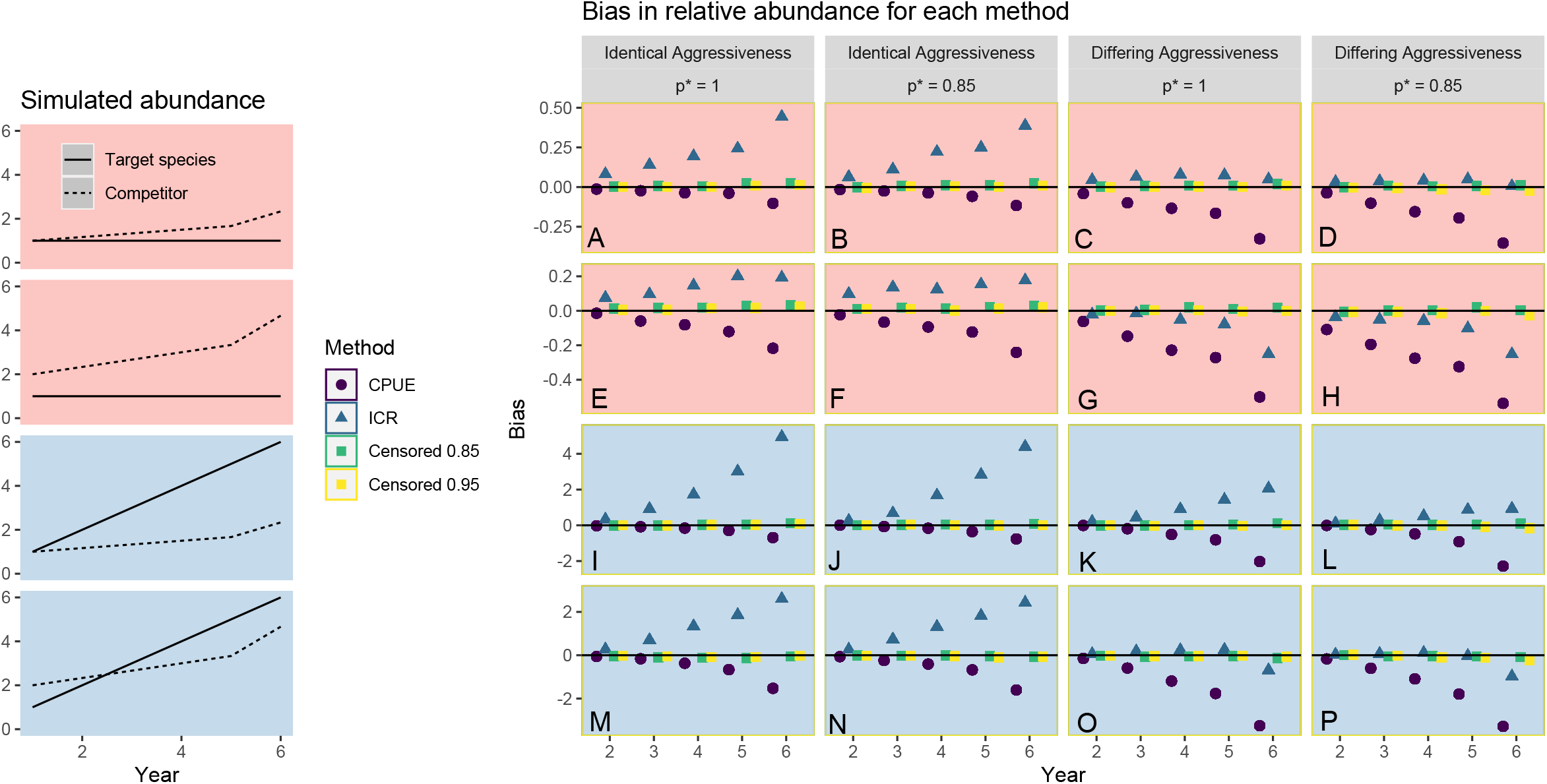
Bias results for experiment 2. Leftmost column shows the relative abundances for the target species and the main competitor in the simulations. The relative abundance of the main competitor determines the average proportion of baits removed for each fishing event. Whether the target species’ abundance remains constant or increases establish whether the simulations are investigating the methods’ type I (red panels A-H) or type II property (blue panels I-P). The ‘other’ species group’s abundance remains constant (not shown). The four right columns indicate whether (i) the true breakdown point, which represents the proportion of baited-hooks removed needed to trigger a decline in each species’ ability to locate the remaining baits, is *p*\* = 1 or *p*\* = 0.85, and (ii) fish of all species arrive to the hooks at the same rate on average across the soak time (“Identical Aggressiveness”), or the target species is typically slowest to reach the hooks (“Differing Aggressiveness”). Results are shown for the CPUE and ICR methods, and for the censored method with assumed values for *p*\* of 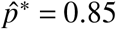 (Censored 0.85) and 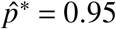 (Censored 0.95). Plotted are the median values of bias, with values close to 0 (solid black line) desired; 95% confidence intervals are omitted since they are almost always smaller than the plotting symbols. Average levels of bias in estimates of relative abundance from the CPUE and ICR methods are high under most scenarios. On average, the level of bias in estimates from the censored method is close to 0 in all scenarios, even when *p*\* is mis-specified 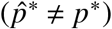.

**Figure 2:**
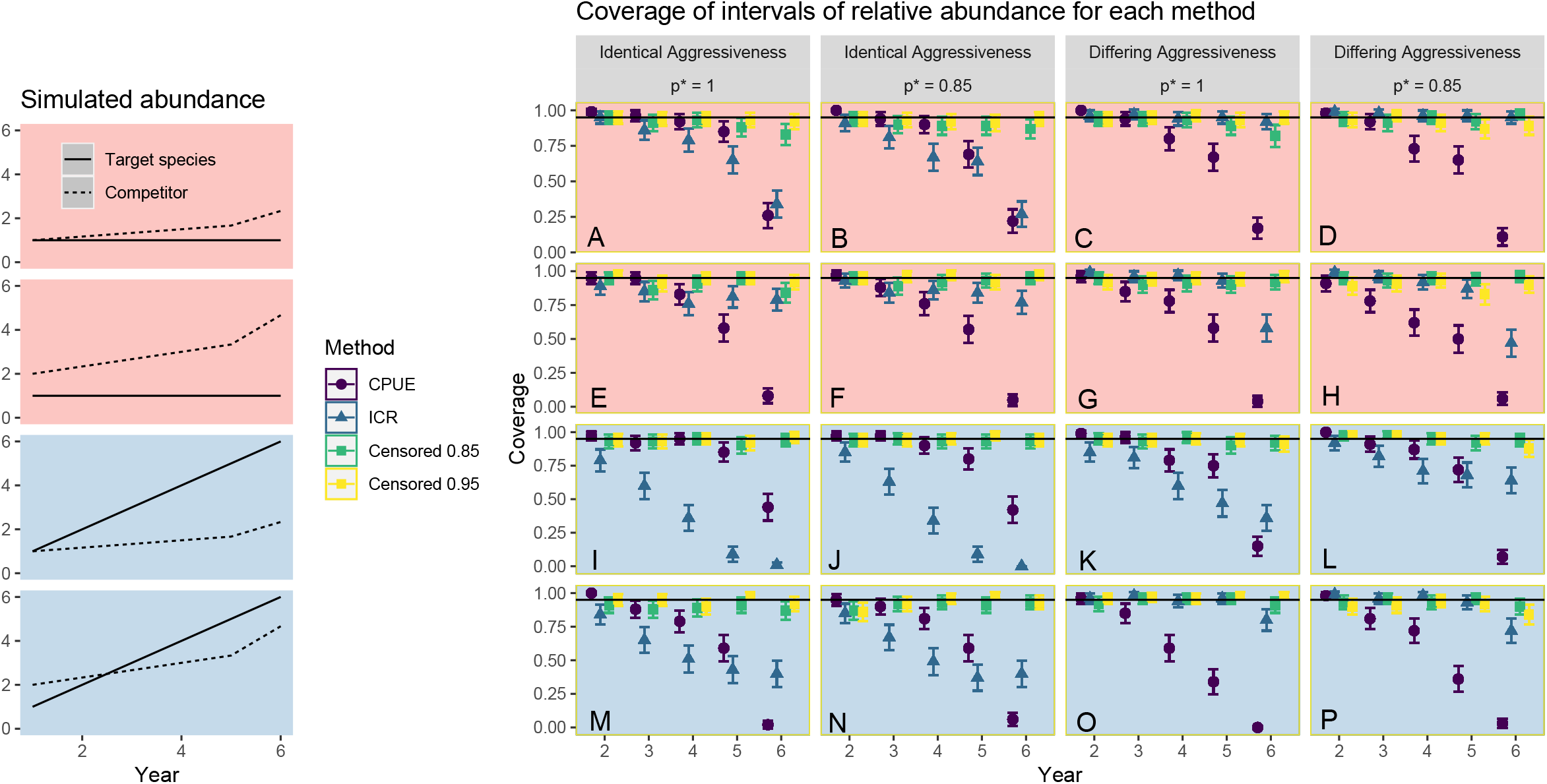
As for Figure 1 but evaluating the nominal coverage of the 95% credible intervals for experiment 2. The mean coverage and its approximate 95% confidence interval is plotted for each scenario. Values close to 95% (the solid black line) are desired. Only the censored method attains close to 95% coverage, with the CPUE and ICR methods failing to satisfy both the type I property (red panels A-H) and the type II property (blue panels I-P) with respect to coverage.

Figure 1 A-H shows that on average, the CPUE method underestimates the relative abundance of the target species and the ICR method overestimates it. Biases from the censored method are negligible. Furthermore, Figure 2 A-H shows that the credible intervals of the censored method attain nominal coverage levels close to the target of 95%, while the coverage for the other two methods can fall below 50%. Thus, the censored method correctly identifies that the relative abundance of the target species is stationary (the type I property), whereas the CPUE and ICR methods frequently infer incorrect trends in relative abundance with high, but misplaced, confidence. The confidence is high because many credible intervals of relative change do not cover zero (the known true value because the target species’ abundance is held constant), and so such intervals are confidently inferring (incorrectly) that the species’ abundance is changing. Incorrectly specifying 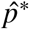 (as 0.85 when *p*\* = 1, or as 0.95 when *p*\* = 0.85 or 1) appears to have a negligible impact on the type I property of the censored method.

Similarly, Figure 1 I-P (investigating Type II properties) shows that the CPUE method typically underestimates the true change in relative abundance by up to 80%, and the ICR method overestimates it by up to 50%. Again, the bias from the censored method is negligible (satisfying the type II property). Figure 2 I-P demonstrates that the censored method attains close to 95% coverage, unlike the other two methods where coverage can fall to 0%. Thus, the correct trends in abundance are rarely captured within the credible intervals from the CPUE and ICR methods. Again, mis-specifying 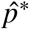 for the target species does not negatively impact the performance of the censored method. For all scenarios considered, up to 100-fold reductions in MSE are observed for the censored method compared to the others (see Supplementary Material), further corroborating the censored method’s improved performance.

Experiment 3 better exhibits the complexities of real-world data (Figures 3 and 4). The target species’ abundance distribution is overdispersed such that moderate schooling can occur (yielding occasional very large catches) and is allowed to be correlated with the abundance distributions of other species within fishing events. While *p*\* = 0.85 is used to generate the data, 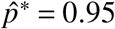 is incorrectly assumed for fitting the censored method. When the experiment was performed, the censored method failed to converge in over 20% of the runs. When this happened, we implemented additional modifications that assisted with the convergence and only slightly impacted performance. However, these convergence issues are much less problematic if the model-fitting routine is started with initial values chosen from the CPUE-based model, a fact only discovered after the simulation study was conducted. We present results only for the converged runs using the unmodified censored method (see Supplementary Material for modified censored results).

**Figure 3:**
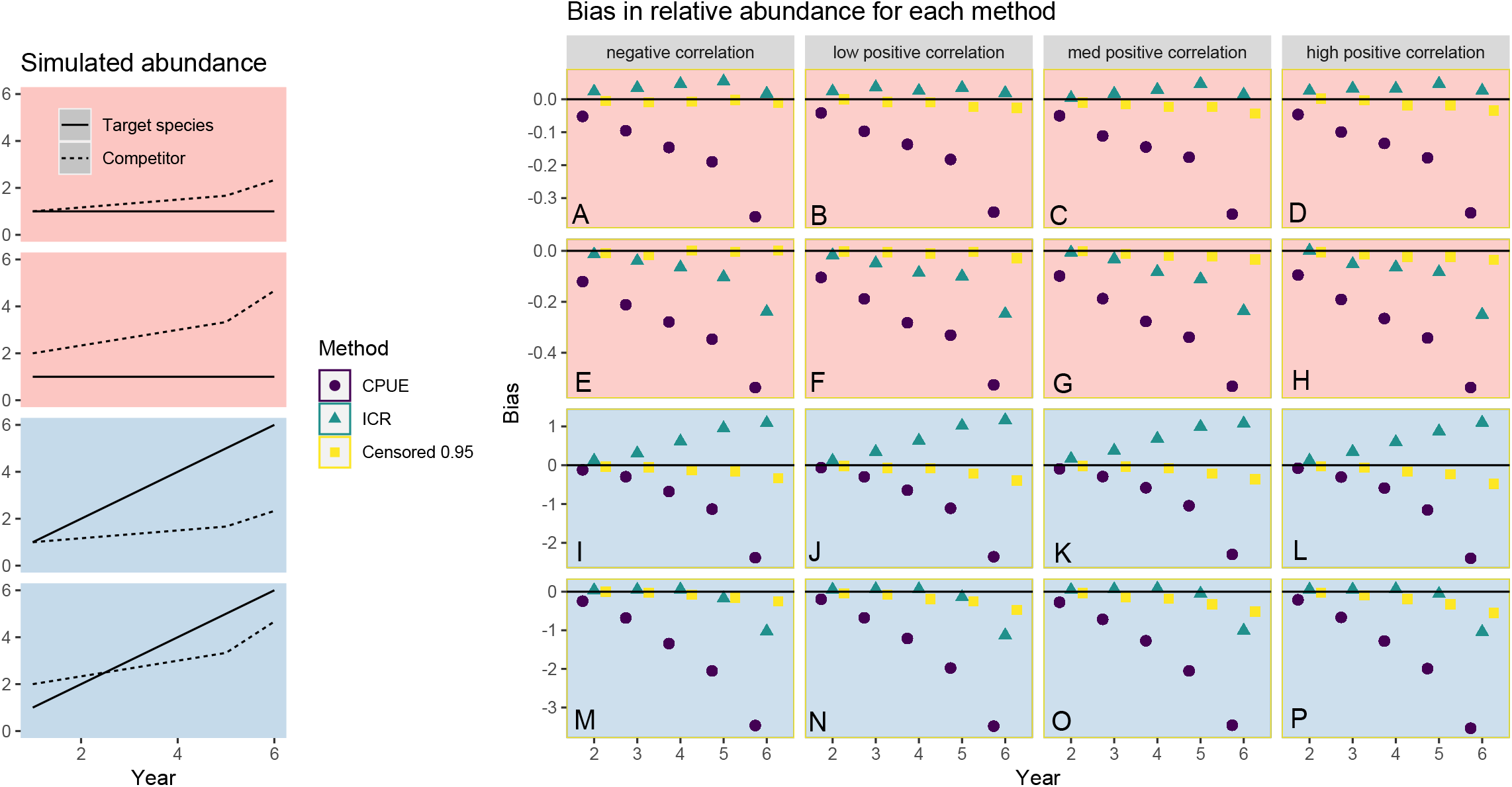
Bias results for experiment 3. Leftmost column and the rows are interpreted as in Figure 1. Columns indicate the levels of correlation between the catch counts of the target species and the ‘other’ species group. Plotted are the median values of bias (confidence intervals are again omitted for clarity) from the 100 repetitions. The ‘Censored 0.95’ method assumes the incorrect value 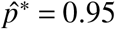 (while *p*\* = 0.85). The censored method infers changes in relative abundance with less bias than the other methods in almost all settings.

**Figure 4:**
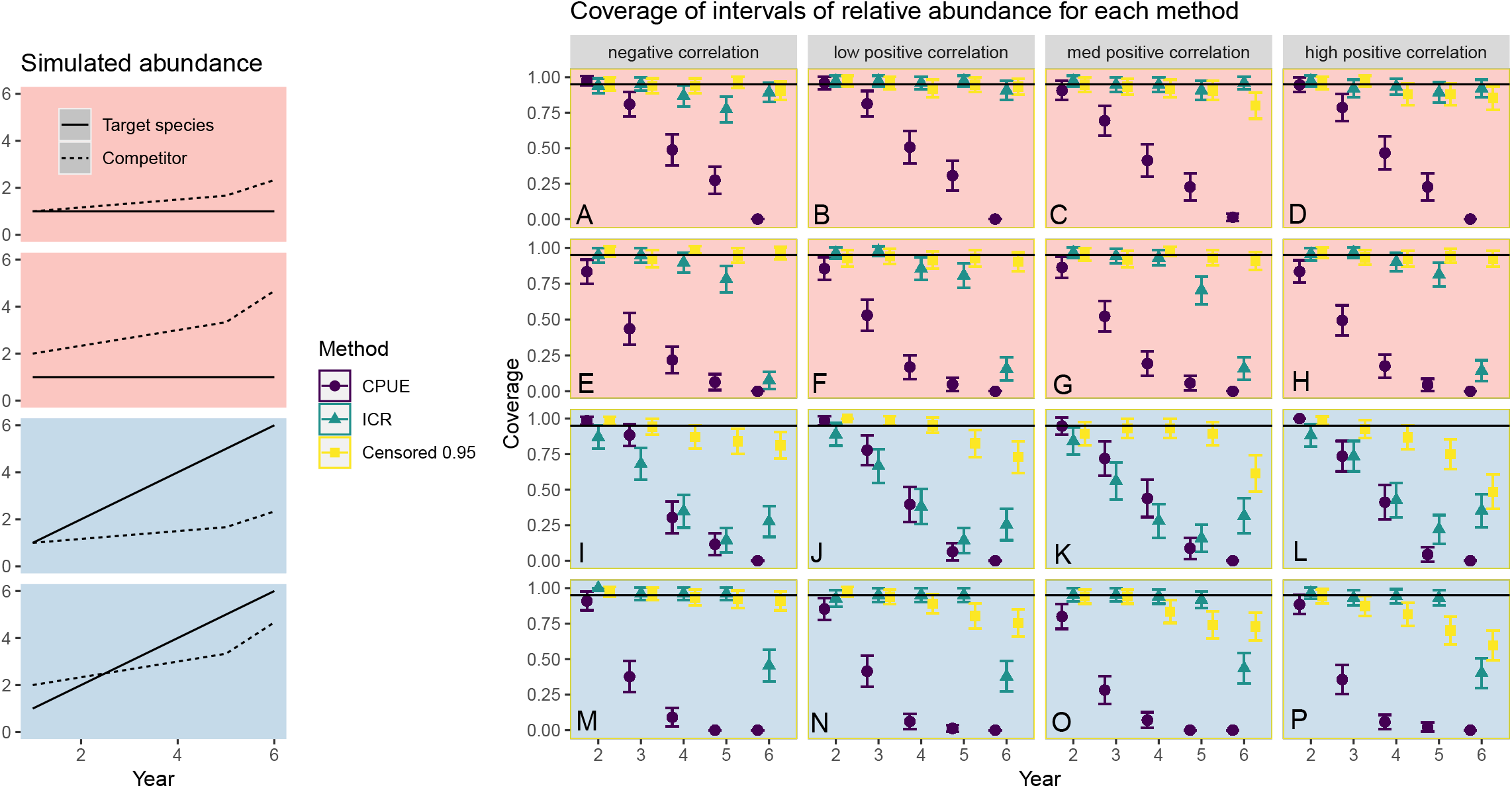
As for Figure 3 but evaluating the nominal coverage of the 95% credible intervals for experiment 3. The mean coverage and its approximate 95% confidence interval is plotted for each scenario. Only the censored method satisfies the type I property (red panels A-H). The censored method outperforms the other two with respect to the type II property (blue panels I-P), but falls short of fully satisfying it.

The censored method again outperforms the CPUE and ICR methods in terms of bias and coverage (Figures 3 and 4). Crucially, the censored method is found to have superior type I (panesl A-H) and type II (panels I-P) properties in these settings. Again, the magnitude of improvement in MSE offered by the censored method can be dramatic, especially when fishing gear are frequently saturated (see Supplementary Material).

A deterioration in the performance of the censored method occurs for experiment 3 when the target species’ abundance distribution is positively correlated with the abundance distribution of the ‘other’ species group (Figure 4 D,H,L,P). This is to be expected - in such scenarios, the fishing events that would have recorded the largest catch counts of the target species are more frequently right-censored due to the higher probability of larger numbers of competitors being present around the longline gear (i.e. increased hook competition). This stochastic dependence between the probability of a fishing event being right-censored and the value of the target species’ abundance distribution is similar to the problem of informative censoring studied in survival analysis (Ranganathan and Pramesh, 2012).

For experiment 4, we repeat experiment 2 but use *p*\* = 0.5 to generate the data. The assumed values of 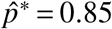 and 0.95 are thus greatly mis-specified. The target species is acutely impacted by hook competition (unbeknownst to the scientist). The performance of the censored method is robust to this mis-specification. Coverage close to 95% is attained for the censored method when the target species abundance does not change over time (satisfying the type I property). When the target species’ abundance changes over time, the coverage of the censored method declines, but never below 50%, whereas for the other two methods the coverage can fall below 10%; thus, the type II property remains superior for the censored method (though is not completely satisfied). The bias and MSE of the censored method are never worse than the CPUE and ICR methods and far better in most settings (see Supplementary Material).

Experiment 5 is purposefully designed to provide a scenario where the target species should be relatively unaffected by hook competition, and so the CPUE method should perform well (and the censored method’s assumption of hook competition is violated). The target species i) occurs in numbers far below the number of hooks (i.e. is non-schooling); and ii) is, on average, able to reach the baited hooks ahead of the competing species groups. Thus, there are often enough hooks available for the target fish. We indeed find that the CPUE method performs best when fish numbers are low (see Supplementary Material). However, this performance improvement is only slight when 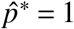. Furthermore, as fish numbers increase (the proportion of baits removed increases), the censored method outperforms across all metrics once again. However, despite the censored method attaining good bias and MSE values in all settings, the coverage can fall below 30% in the most challenging settings (and lower still for the CPUE method). In summary, the censored method performs at least as well as the CPUE method for species that are largely ‘immune’ to hook competition. However, reported estimates of uncertainty from all methods should be treated with caution as they are likely to be underestimates.

So, across experiments 1-5, the CPUE method is unsuitable for use in almost all simulation settings tested due to its high sensitivity to changes in the relative abundance of competing species. The one exception occurs when the target species is both non-schooling and able to reach baits ahead of competing species (experiment 5). The overall performance of the ICR method is worse in many settings. Most concerning is the consistent failure of it to satisfy the type I property; in some settings a change in abundance was incorrectly, but confidently, inferred over 85% of the time (namely when coverage falls below 15% in Figure 4 I-P). Again, this is in agreement with the earlier theoretical example which highlighted the sensitivity of the ICR method to changing levels of non-target species’ abundances.

Conversely, the censored method performs well in almost every setting tested, consistently attaining both the type I and type II properties. Bias, MSE, and coverage are frequently improved, even when the data-generating mechanism is inconsistent with the modelling assumptions (e.g. when strong correlations exist between the species’ abundance distributions). Perhaps most promisingly, the relative improvements in performance offered by the censored method appear to be highly robust to 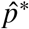 being poorly chosen, and to the implementation of crude methods to improve the model convergence in data-poor settings.

We now investigate how a sensible value of 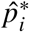 could be chosen. We explore trends between the mean observed catch counts, *c_itk_*, and the proportion of baits removed, *p_tk_*, after controlling for time effects, in data simulated under numerous scenarios. The top panel of Figure 5 shows: i) a strongly decreasing trend between *c_itk_* and *p_tk_* for the target species of experiments 2-4, ii) a negligible trend for the hook-competition-immune target species of experiment 5, iii) a strongly increasing trend for the main competitor species under experiments 2-4. A line is drawn at *p*\* = 0.85 used to generate the data which shows that for i), the decreasing trend begins close to *p*\*.

**Figure 5:**
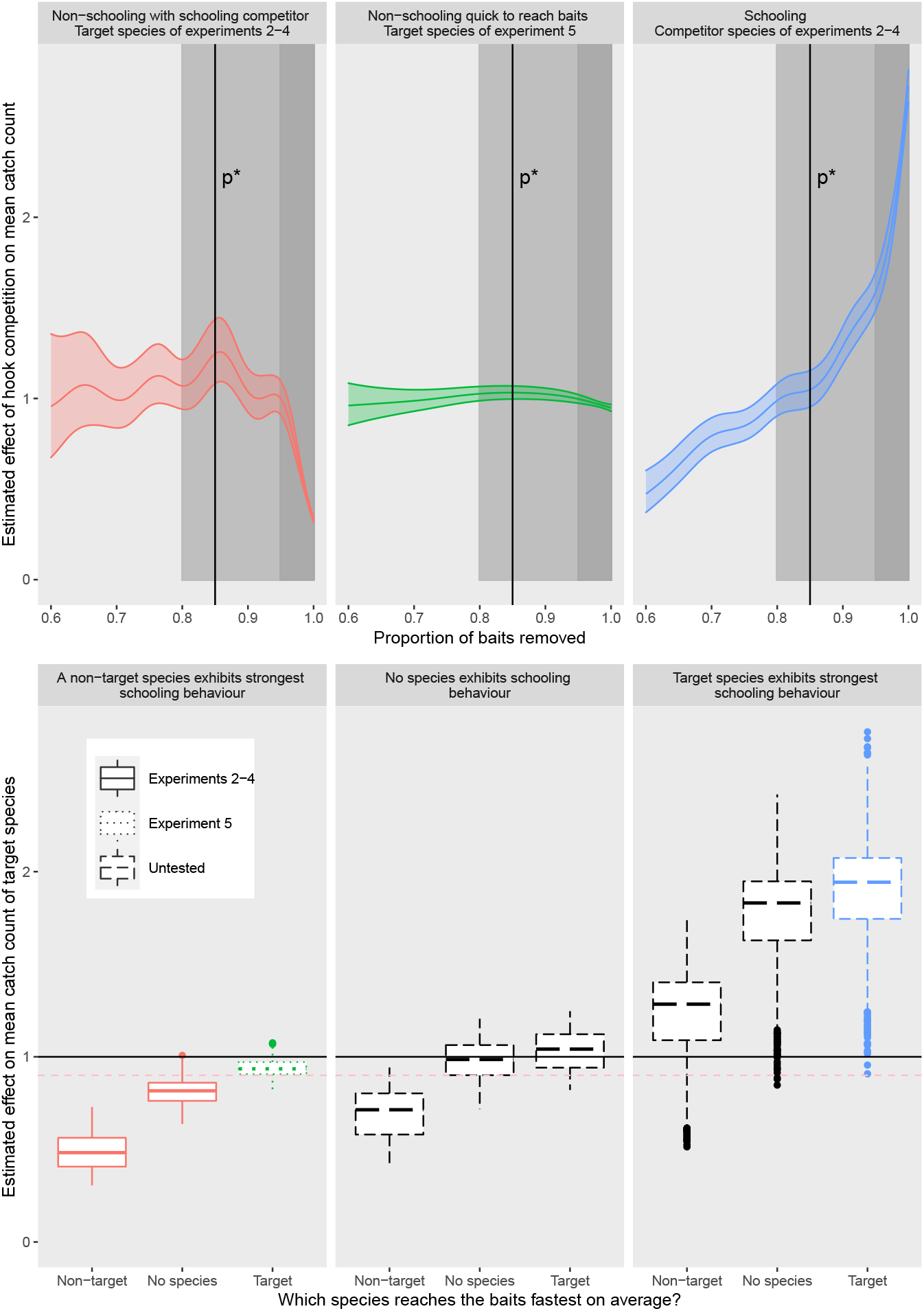
Top: model-estimated trends between the mean observed catch count vs. proportion of bait removed in each of three representative datasets simulated under scenarios described in the panel titles. Vertical lines show true value of *p*\*; a changepoint beginning at *p*\* can be seen in the leftmost trajectory. The two intervals with the darkest shading define the medium and high hook saturation levels used in Experiment 6. Bottom: boxplots of all the results in experiment 6, approximately computed as the relative values of the trend lines within the two shaded intervals. The horizontal dashed line shows where a 10% decline in mean catch count occurs (at the value 0.9 due to the multiplicative scale used). Colours point to the representative trajectories above. The linetype maps the scenario to the corresponding experiment (or lack of). Only the scenarios in the leftmost panel have been simulation tested. Declines of > 10% are seen under scenarios matching experiments 2-4, whereas only small declines are seen under experiment 5’s settings. Both the speed of a species to reach bait and its relative degree of schooling impact the trend.

These observed trends between the mean observed catch counts and the proportion of baits removed motivates experiment 6. To assess whether the conclusions from the exploratory analysis generalize more broadly, we sample longline fishing events under a range of scenarios (see Supplementary Material). For each simulated dataset, we compare the mean observed catch count at high levels of *p_t_k* (*p_tk_* ≈ 1) versus medium levels of *p_tk_* (*p_tk_* ≈ 0.85).

Experiment 6 shows that a > 10% decline in mean catch count from medium to high levels of *p_tk_* almost always occurs for a species that is very slow to reach the baits and/or is outcompeted by large schools of a competing species (Figure 5). The exploratory analysis suggests choosing 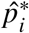 at the value of *p_tk_* on the curve where the decreasing trend begins. Experiments 2-3 suggest that the censored method should perform very well across all metrics for such a species. Experiment 4 implies that the performance is robust to errors in 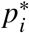 estimates. Conversely, experiment 6 shows that > 20% increases typically only occur for strongly schooling species that frequently saturate the fishing gear (Figure 5). Experiment 1, which considered a species that was solely responsible for saturating the fishing gear, suggests that the censored method will outperform competing methods with 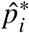 set to 1. However, estimates of uncertainty may be moderately underestimated. Lastly, species with small trends are expected to be largely immune to hook competition, either because it can reach baits in small numbers ahead of its competitors, or because there is an absence of a strongly schooling competitor. Experiment 5 considered such a species, and results suggest that 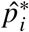 be set to 1. However, levels of uncertainty may be greatly underestimated here. We later use these decision rules for selecting 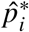 in the case study.

### 3.2 Case study

We compute Canadian coastwide relative abundance indices of 11 species from the IPHC longline survey data using the CPUE, ICR, and censored method, plus bootstrapped raw CPUE values (similar to methods currently used). The focus of the analysis is to produce annual relative abundance indices that could be used as inputs for formal stock assessments, and so we do not place strong modelling assumptions on the year-to-year trends (such as using random walks, smoothing splines, or polynomials). The bootstrapped CPUE values are computed using only the data collected at the 107 stations that recorded data each year, while the other methods use data from all 171 stations and attempt to correct for the differences in stations used each year with random effects. Each resulting time series is divided by its geometric mean to produce comparable relative abundances indices (see Supplementary Material for details).

Nine species show statistically significant declines in *c_itk_* exceeding 10% as *p_tk_* increases, after controlling for the effects of time and for the differences in mean abundance across the fishing stations (Figure 6). Of these, the decline appears to begin at *p_tk_* ≈ 0.85 and *p_tk_* ≈ 0.95 for 2 and 7 species respectively. The conclusions of experiment 6 suggest fixing 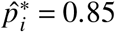 and 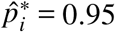. Results from experiments 2-4 suggest the censored method will be able to infer the relative abundance of these species with high accuracy, high precision, and with accurate uncertainty estimates.

**Figure 6:**
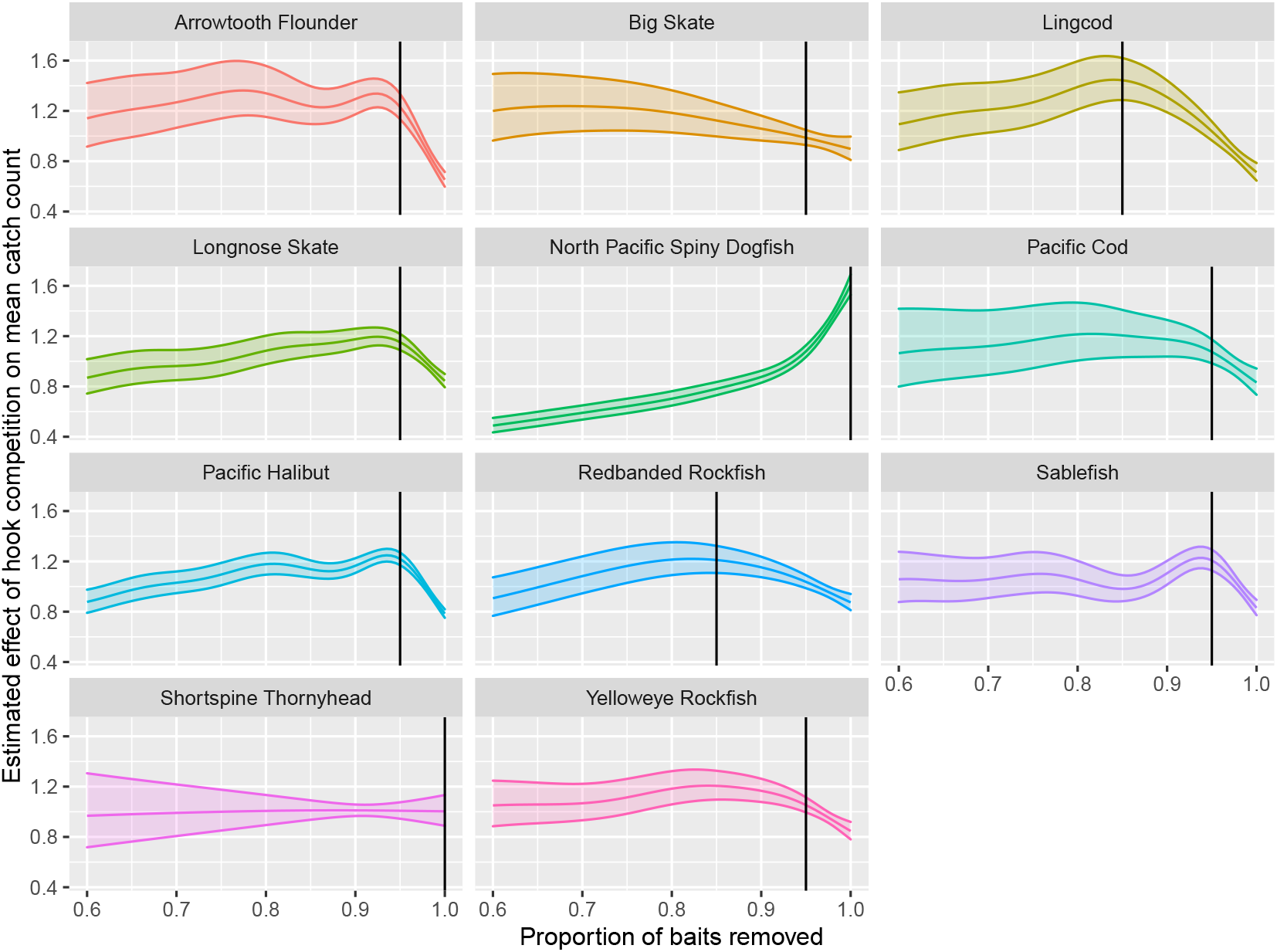
Estimated effects of hook competition in the IPHC data (after controlling for temporal effects), shown as the relative change in the the mean catch count of each species as the proportion of baits removed increases. Effects are shown on the multiplicative scale to match those from Figure 5. Models are fit to the catch counts from all of the fishing events. Vertical lines indicate the species-specific 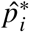 values from this analysis. All species experience a statistically-significant decline in the mean catch count exceeding 10% (approximately a drop larger than 0.1 on the y-axis), except for dogfish, which experiences a statistically significant increase in mean catch count exceeding 10%, and Shortspine Thornyhead, whose estimated 1% decline is non-significant. For these two species results from experiment 6 suggest 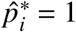. Plots of Lingcod and Redbanded Rockfish suggest 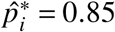 (where the decline commences) and the remaining plots suggest 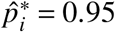.

Shortspine Thornyhead showed no significant trend between *c_itk_* and *p_tk_*, suggesting they may be relatively unaffected by hook competition in this data set. North Pacific Spiny Dogfish (hereafter just ‘dogfish’) show a steady increase in *c_itk_*, even accelerating as *p_tk_* reaches 1, in agreement with our knowledge of them as a highly-mobile schooling species and with the large increasing trends seen in schooling species’ catch counts in experiment 6 (Figure 5). Experiment 6’s results recommend fixing 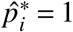 for both species and experiments 1 and 5 advise that reported levels of uncertainty be viewed with caution.

The 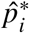 values for each species are then used to calculate the relative indices for the censored method, shown alongside the other methods in Figure 7 for Lingcod, Arrowtooth Flounder, and dogfish (see Supplementary Material for remaining species).

**Figure 7:**
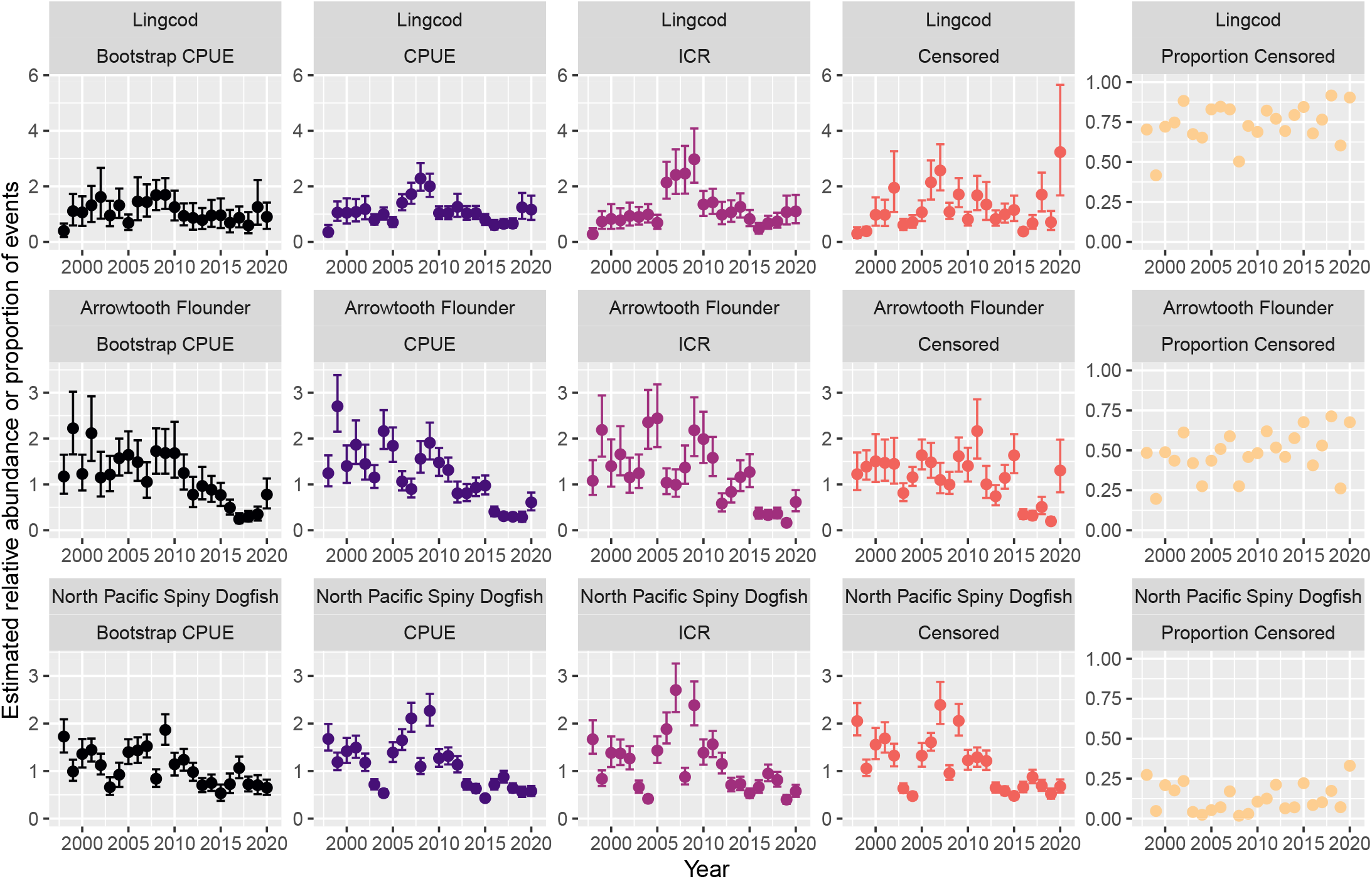
Plots showing normalized (each time series is divided by its geometric mean) estimates of relative abundance from the CPUE, ICR, and censored method, plus normalized bootstrapped percentile intervals of CPUE, for Lingcod, Arrowtooth Flounder, and dogfish. Right column shows the proportions of fishing events each year considered censored (i.e. mean proportion of events with 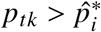). Large differences between the estimates are seen for Lingcod and Arrowtooth Flounder, whereas estimates show good agreement for dogfish. No obvious temporal trend in the proportion of censored events is apparent. Bootstrapped estimates consider only the subset of 109 stations for which data were successfully collected each year, all other methods use data from all 171 stations. The censored method’s trends differ greatly from those of the CPUE and ICR methods, especially in the time periods with the greatest variation in the proportion of censored events: 1998-1999 and 2018-2020.

Interestingly, the CPUE and ICR methods’ estimates are in close agreement for all species, albeit with the ICR method estimating fluctuations with greater magnitude. The censored method’s estimates deviate greatly from the CPUE and ICR methods’ estimates for Lingcod and Arrow-tooth Flounder, most notably during the time periods of 1998-1999 and 2018-2020 when censor-ship levels oscillated. Figure 6 outlines why. Both species show a decline in mean catch counts exceeding 40% between fishing events with a proportion of baits removed around 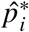 versus those with all baits removed. Furthermore, Figure 7 shows that the average proportion of censored fishing events (i.e. fishing events with 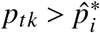) is close to 75% and 50% for Lingcod and Arrowtooth Flounder respectively. Thus, it is estimated that fishing events are frequently impacted by hook competition bias, and that the magnitude of this bias is very high for both species.

Conversely, although Figure 6 suggests that estimates of dogfish abundance are sensitive to hook competition bias caused by complete bait depletion, Figure 7 shows that it is relatively rare for fishing events to return zero baits (averaging around 10% of fishing events). Consequently, estimates of relative dogfish abundance from the censored method are in close agreement with those from the CPUE method. See Supplementary Material for full details.

Figure 8 shows the magnitude of the differences between the estimates from the CPUE and the censored method, and how these change with the annual proportion of fishing events that are censored. Strikingly, for Lingcod the censored method can yield relative abundance estimates that are deflated by 50% compared to those from the CPUE method in years with low censorship, but inflated by 150% in years with high censorship. Note that although the censored method will not deflate catch counts for individual fishing events, deflation of the annual relative abundance estimates occurs because of the scaling of each multi-year time series by its geometric mean. The magnitude of inflation/deflation at a given level of censorship varies greatly from species to species (see Supplementary Material). Thus, the censored method identifies a unique level of hook competition sensitivity for each species. Lingcod is estimated to have been most affected by hook competition. Dogfish and Pacific Halibut are estimated to be the least affected, with maximum inflations in their relative abundance estimates of around 20%, and 50% respectively compared to the CPUE method. The species-specific effects of hook competition is unlike the ICR method which assumes the same impact of hook competition on all species.

**Figure 8:**
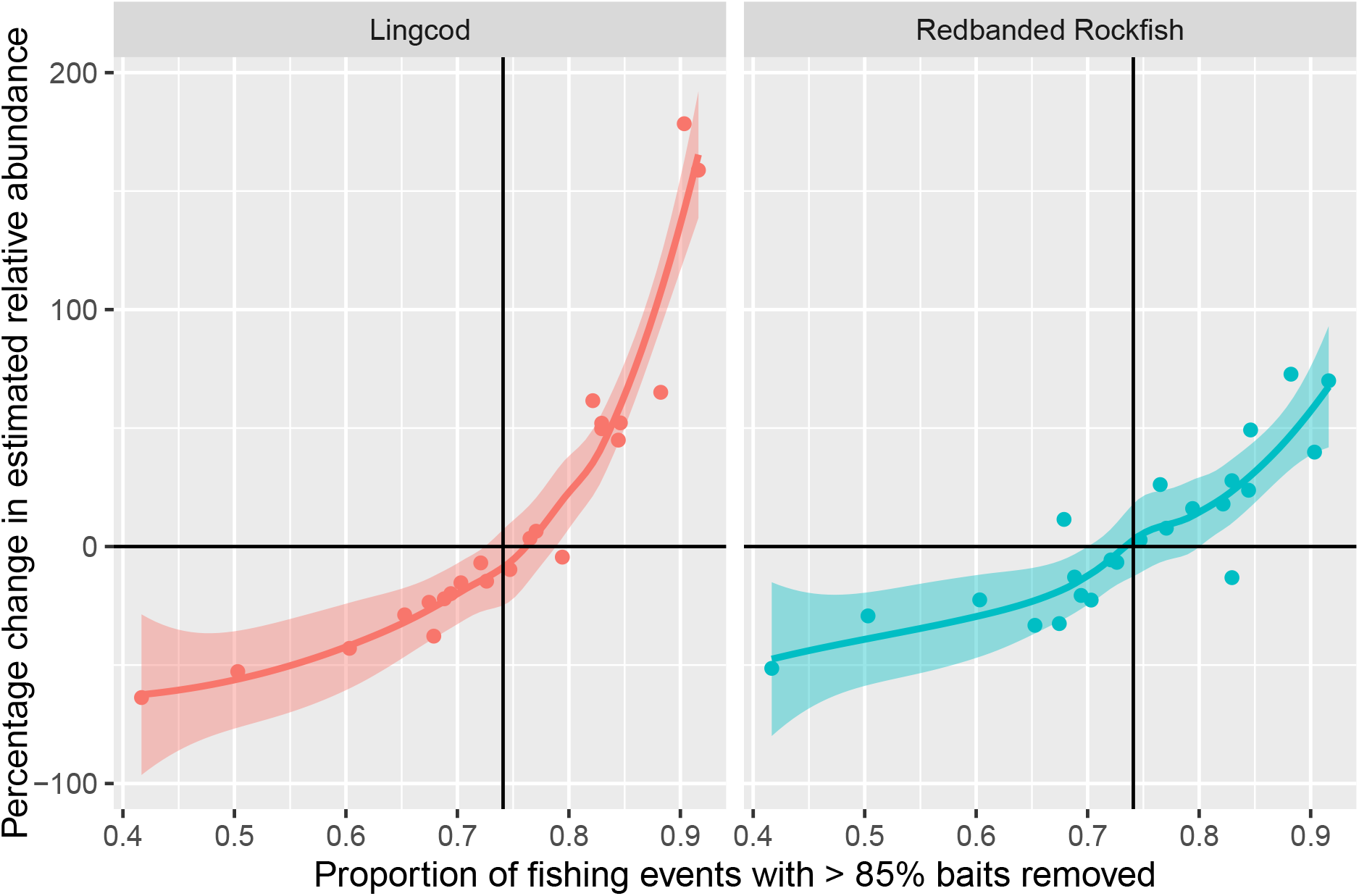
The percentage change in normalized relative abundance estimates from the CPUE method to those from the censored method (one dot for each year) against the annual proportion of censored fishing events, for the two species with 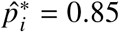. Horizontal lines indicate no change between the methods, and vertical lines show the average annual proportion of fishing events experiencing at least 85% bait removal. Clearly i) relative abundance estimates from the censored method are significantly deflated compared to the CPUE method in years with low levels of censorship, and significantly inflated in years with high levels; ii) the inflation/deflation trend is nonlinearly related to the proportion of censored observations; iii) the magnitude of inflation/deflation varies across species (even with the same value of 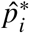) and can be very large. The censored method infers that Lingcod is more severely affected by hook competition than Redbanded Rockfish.

## 4 Discussion

The censored Poisson method introduced here represents a novel approach for handling the biasing effects of hook competition on estimates of relative species abundance collected with longline gear. We showed that our method generally outperformed the methods currently used to account for hook competition. Unlike the CPUE and ICR methods currently used, our method allows researchers to correctly identify when fish populations appear stable through time, and when they show marked changes. Crucially, the new approach moves away from the earlier mathematical approaches that modelled hook competition with fixed-rate ODEs and towards a purely statistical treatment of the problem by viewing hook competition as a data-quality issue.

Treating the catch counts of a species from longline fishing events that experienced the highest levels of bait removal as right-censored allows us to appropriately express our ignorance with the true underlying processes driving hook competition, such as species-specific bait location abilities, mechanical bait loss, and bait removal due to scavengers. Furthermore, the censoring approach also provides a natural way of linking the observed catch counts, which are upper bounded by the number of deployed baited hooks, with the somewhat unbounded density of each species. Conversely, attempting to mathematically describe the numerous interacting processes that govern the hook competition within a system of ODEs is a mammoth challenge and it is unclear how to link the ODE rate parameters to the underlying species densities without requiring additional strong assumptions (e.g. independent bait removal events, non-overlapping hook-level bait odour concentration plumes).

Whilst the simulation study demonstrated the robustness of the censored method to gross mis-specifications of the breakdown point 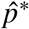, it also clearly showed that improvements in prediction accuracy and uncertainty quantification are both possible if this breakdown point can be better estimated. We introduced an empirical approach to estimating the breakdown point. However, better estimation may be possible by conducting hook-spacing experiments (Sigler, 2000). One can experimentally increase the inter-hook spacing until a significant decrease is observed in the catch rate, and then map this spacing level to an equivalent proportion of baited hooks removed. Sigler (2000) found that Sablefish exhibited an 8% decline in catch rate once the hook spacing was 42 m – equivalent to around 5% of the baits remaining for a standard survey skate with 2-m spacing. Thus, an experimentally-justified estimate for the Sablefish-specific breakdown point would be when 95% of the hooks have been taken, which is the same value found by our empirical approach. Future research on other species could validate this experimental approach and use it to further develop empirical approaches for estimating *p*\*.

Species-specific breakdown points could be linked to the presence of one or more problematic species. A species may be problematic due to many reasons, including it scaring the target species away from the baited hooks (e.g. predators). If only a few problematic individuals present around the longline gear are able to cause large decreases in the observed catch counts of a target species, then even small fluctuations in the abundances of these problematic species could strongly confound the relationship between the catch counts of a target species and its abundance. The approach outlined here for choosing the breakdown point could be extended for such a scenario, by conducting simulation studies to develop methods for identifying the existence of problematic species and choosing a suitable breakdown point.

While the censored method performed well in all settings tested, uncertainty levels were underestimated when applied to species that were immune to hook competition. Knowing which species are insensitive to hook competition would allow researchers to acknowledge that while the censored method likely estimates the relative abundance of these species with low error, the corresponding estimates of uncertainty are likely too low. Identifying immune species should be possible with experimentation, such as identifying the species with the fastest arrival times with hook timers or ROVs (e.g. Obradovich, 2018), and future methodological developments could provide methods to correct for these biases.

Whilst the case study focused on longline data collected by the IPHC for capturing demersal species, the method should apply equally well to midwater/surface longlines for estimating the relative abundances of pelagic species, requiring only a conceptual change towards density quantities and integrals in three-dimensional space with respect to volumes of water instead of surface areas of seafloor. Computational implementation remains the same. For longline surveys that record biomass rather than numbers, the censored likelihood function can be modified to one that handles zero-inflated continuous data.

Mechanical bait loss and bait removal/blocking due to large invertebrates have both been identified as possible sources of bias in abundance indices (High, 1980; Obradovich, 2018). For example, Obradovich (2018) observed sea stars (*Astroidea spp.*) physically blocking baited hooks in underwater ROV experiments, but rarely observed the offenders on deck upon gear retrieval. Similarly, Grimes *et al.* (1982) observed crabs and sea stars remove and/or block 50% of longline bait within 4 hours and Obradovich (2018) also observed Giant Pacific Octopuses (*Enteroctopus dofleini*) and multiple crab species (*Cancridae sp., Lithodoidea spp., Majidae sp., Oregoniidae sp.*) remove large quantities of bait, despite only a single octopus and no crabs ever being observed on deck. Large annual fluctuations in the population sizes of both crabs and Giant Pacific Octopuses have been observed (Hartwick *et al.*, 1984; Zhang and Dunham, 2013). While the censored method can easily account for bait removal, it will fail to account for any biases caused by baits being blocked but not removed. To correct for this: i) the local average proportion of blocked baits could be estimated and used to appropriately decrease the breakdown point; ii) a unique breakdown point could be estimated each year following the empirical approach outlined in this paper – an earlier decline in mean catch count should be visible in years with high levels of blocking; iii) a sensitivity analysis could be conducted, averaging conclusions across a range of plausible breakdown point values.

Spatio-temporal methods are being increasingly proposed and implemented in fisheries science (Thorson *et al.*, 2015, 2016). For example, the IPHC analyzes the halibut catch counts from their survey using a spatio-temporal approach (Webster *et al.*, 2020). The censored Poisson method, derived in equations (1), (3), and (8), was explicitly set up to be extended by allowing for Gaussian Markov random fields to be included within the linear predictor. Taking a spatio-temporal approach to estimating relative abundance can lead to improved precision and it can simplify the construction of joint models using multiple data sources with data sampled at disjoint spatial locations (such as bottom trawl data and longline data; Grüss and Thorson 2019; Webster *et al.* 2020). In fact, since the mean of the censored Poisson model (8) describes a species’ density instead of a rate from differential equations, as is the case with other hook competition approaches, such joint models are more easily and justifiably constructed. An interesting future research question could be to determine if the breakdown point can be sensibly chosen by maximizing a measure of agreement between longline and trawl (or other hook-competition-free fishing method) subcomponents within a joint model.

In addition to the inclusion of spatio-temporal random effects to capture latent spatio-temporal correlations, numerous other improvements to the relative abundance indices computed in the case study could be considered. Other considerations include covariates believed to describe the species’ habitat preferences, including a survey-timing adjustment (as done by the IPHC for halibut, see Webster and Stewart 2013), and allowing for nonlinear effects of effective skate on the mean catch counts of the four species for which nonlinear effects were detected (see Supplementary Material). The purpose of the indices presented in the case study is to provide the reader with an understanding of the magnitudes of changes possible between the competing methods.

Relative abundance indices are frequently produced to monitor populations and to use as inputs to formal stock assessment models. Changes on the order of 50%-150%, as found here, could lead to substantial differences in the resulting outputs from such models and in the consequent advice to fisheries managers. Thus, we recommend our simulation-tested censored method be considered for such applications.

## Acknowledgements

For postdoctoral funding that enabled this work, we thank the Canadian Statistical Sciences Institute (CANSSI) Collaborative Research Team ‘Towards Sustainable Fisheries: State Space Assessment Models for Complex Fisheries and Biological Data’ led by Joanna Mills Flemming, plus Fisheries and Oceans Canada for funding through SPERA (Strategic Program for Ecosystem-based Research and Advice) and FSERP (Fisheries Science and Ecosystem Research Program). MAM also acknowledges funding from the Canada Foundation for Innovation (John R. Evans Leaders program), the British Columbia Knowledge Development Fund Program, the Canadian Research Chairs program, and Natural Sciences and Engineering Research Council of Canada (NSERC). We thank Patrick Thompson and Shannon Obradovich for insightful comments on an earlier draft, and also thank Håvard Rue for developing the censored Poisson INLA family (cenpoisson2) for use in this work.

## Supplementary Material – Extended Methods and Results

Supplementary Material for ‘A statistical censoring approach accounts for hook competition in abundance indices from longline surveys’ by Joe Watson, Andrew M. Edwards, and Marie Auger-Méthé. Submitting to *Canadian Journal of Fisheries and Aquatic Sciences.*

### S.1 Simulation study set-up and additional results

We generate data for the simulation study by determining the underlying numbers of each species that are attracted to the fishing gear (Step 1), determining how quickly the individual fish would take a bait (in the absence of hook competition) depending on the aggressiveness of the species (Step 2), and explicitly simulate hook competition to determine which individual fish actually get caught by the gear (Step 3). We then apply competing methods to the resulting simulated catch counts to infer temporal changes in the abundance of each species (Step 4).

#### S.1.1 Step 1 – generate the numbers of each species attracted to the fishing gear

We sample the number of individuals of each of three species groups that are attracted to the longline gear, which involves first specifying the underlying species density *λ_it_* (**s**). We want to illustrate the strengths and weaknesses of the competing methods in the simplest settings, free from confounding by external factors and spatial heterogeneity. So we do not explicitly simulate the influence of covariates on the population or on the response function that determines whether an individual fish removes bait. This involves setting *x_itkm_* = *w_itkr_* = *Z_it_* (**s**_*k*_) = 0 in (13), and also we can set *β*_*i*0_ = 0 (it can be set to an arbitrary constant that would be absorbed by *α*_*it*0_ anyway).

**Table S.1:** Extra notation used primarily in the Supplementary Material.

|  |  |
| --- | --- |
| $L_i$ | lower bound for the probability that an attracted fish of species $i$ is captured by a baited hook, equal to the probability when only one baited hook remains |
| $S_{tk}$ | soak time of fishing event $k$ in year $t$ |
| $\tilde{c}_{itk}$ | response variable of interest, equal to $c_{itk}$ for the CPUE and censored method, and multiplied by the hook-competition adjustment factor for the ICR method, to enable writing all models under the framework (S.14)–(S.18) |
| $Q$ | fishing events with target species' catch counts lying above the $Q^{th}$ quantile of annual observed catch counts are considered high quality when the 'quantile adjustment' is implemented |
| $c'_{itk}$ | the number of species $i$ caught during the first stage of the soak time in the $k^{th}$ fishing event of year $t$ |
| $c''_{itk}$ | the number of species $i$ caught during the second stage of the soak time in the $k^{th}$ fishing event of year $t$ |
| $\mathbf{g}_{itk}$ | a vector for which each element corresponds to a bite (gobble) time of a fish of species $i$ that was attracted to the gear during fishing event $k$ of year $t$ |
| $p(\tau)$ | the proportion of baits removed $\tau$ units of time into the soak time |
| $u_{itk}$ | upper bound of the catch counts of species $i$ in year $t$ at fishing event $k$ |
| $\mu_{it}$ | expected value of $C_{itk}$ assuming no hook competition and no spatial heterogeneity between stations |

Then (13) reduces to log *μ_itk_* = *α*_*it*0_, and (1) and (10) reduce to log *λ_it_* (**s**) = *α_it_* (**s**) = *α*_*it*0_, such that *μ_itk_* = *λ_it_*(**s**). By spatial homogeneity, *μ_itk_* = *μ_it_* = exp(*α*_*it*0_), with *μ_it_* determining the expected catch count of species *i* in year *t* during any fishing event, assuming no hook competition. These settings help constrain the number of experiments conducted. Extra notation used primarily in the Supplementary Material is summarised in Table S.1.

For species group *i* ∈ {1 = competitor,2 = other, 3 = target}, in year *t* = 1,2,…, 6, and station (replicate) *k* = 1,2,…,*K_t_*, the catch count in the absence of hook competition can be sampled from the following model based on (5) and (13):

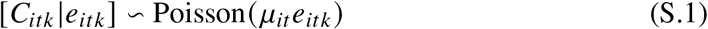

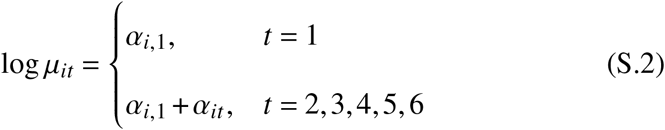

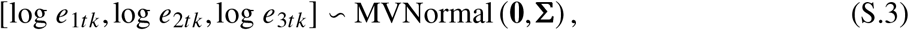

where *e_itk_* represents lognormal random noise to account for overdispersion in catch counts, *α_it_* prescibe the *μ_it_* (though for ease of understanding we give values of *μ_it_*), and **∑** describes: i) the magnitude of overdispersion in the count distributions of the three species groups; ii) the prevalence and severity of schooling behaviours exhibited by each species; iii) the correlations between the count distributions of the three species groups. The covariance matrix **∑** is fixed in time and so exp (*α_it_*) for *t* > 1 defines the relative abundance of species *i* in year *t,* compared with its abundance in year 1 (considered as the baseline level). The exp (*α_it_*) for *t* > 1 are the inferential targets for the simulation study. We chose them since they remain identifiable when *β*_*i*0_ ≠ 0 in (13) (expected in more general settings).

The catch count *C_itk_* of each species in the absence of hook competition is the same as the number of individual fish of that species that are attracted to the fishing gear during the fishing event. We have not yet put in any constraints regarding the number of hooks available to the fish.

#### S.1.2 Step 2 – sample the bite times at which individual fish would take the bait given unlimited hooks

Next, we sample the bite times at which each fish would gobble the bait if there were unlimited hooks, denoted **g**_*itk*_, for which each element of the vector **g**_*itk*_ corresponds to each of the *C_itk_* individuals of species *i* attracted to the gear during the 5-hour soak time of the fishing event at station *k* during year *t*. So there is a vector **g**_*itk*_ for each species and fishing event, and each element of the vector corresponds to the bite time of each of the individual fish that are attracted to the gear. We can vary the aggressiveness of each species by varying the assumptions regarding their bite times.

We consider three distributions to sample from. First, is the truncated exponential distribution with rate 1 (truncated to the 5 hours). The majority of the individuals sampled from this distribution will bite the hooks early on in the soak time. Thus, the exponential distribution reflects the bite times of an ‘aggressive’ species.

Next, is a mixture distribution of a unit rate truncated exponential distribution and a uniform distribution with limits 0 and 5. The mixture weights are fixed at 0.5. This distribution reflects a large species group comprised of fish species with different levels of aggressiveness, olfactory sensitivities, and other biological characteristics. Under this distribution, some of the species are expected to bite early on in the soak time, while others are expected to bite the hooks more consistently throughout the entire soak time.

Finally, a uniform distribution with limits of 0 and 5 is considered. Individuals sampled from this distribution are expected to bite the hooks consistently throughout the entire soak time and are thus relatively ‘slow’ to reach the baited hooks on average (low ‘aggressiveness’).

#### S.1.3 Step 3 – explicitly simulate hook competition to see which fish get caught

Clearly, step 1 may sample more attracted individuals than the *h_tk_* = 800 hooks that are available during every fishing event, yet step 2 has assumed unlimited hooks. So we now sequentially order each unique attracted fish in ascending order of the sampled bite times (so the more aggressive species are more likely to get to the hooks early on, whereas slower species are expected to bite later). Then, for each ordered fish, a Bernoulli random variable with known ‘capture’ probability is sampled from to determine if the fish successfully takes a bait and is captured by an available hook and is subsequently recorded in the data. If no more hooks are available then the gear is saturated and no more fish can be caught. We assume that if a fish takes a bait then it is captured (fish of all species that steal bait without being caught could be explicitly modelled as another species group using counts of empty hooks).

We consider two situations. The first assumes that if any hooks are available then the fish will be captured (i.e. the Bernoulli capture probabilities are all 1). This setting implies the species is adept at locating remaining baited hooks, even when few remain actively fishing, e.g. observed Sablefish behaviour in hook timer experiments (Rodgveller *et al.*, 2011).

The second situation assumes that the species’ capture probabilities can decline as baits are removed. Specifically, a threshold 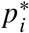 exists for all species. For simplicity in the experiments we set 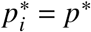. The capture probability of a fish is < 1 if the proportion of baits removed from the longline gear at the fish’s specified bite time is > *p*\*. The probability equals 1 when the proportion is < *p*\*. Explicitly, the species-specific probabilities decrease linearly from 1 as baits are removed beyond *p*\*, reaching a species-specific lower bound, *L_i_*, when one bait remains. For example, if *p*\* = 0.85 and *L_i_* = 0.2, then all attracted individuals of species *i* with bite times before 85% of the baits have been removed are caught, but after that the probability of capture declines linearly from 1 to 0.2 as further baits are removed. Changing *L_i_* changes the severity of the hook competition effects experienced by the species, and setting *L_i_* = 1 means that hook competition only occurs when all the baits are removed (regardless of *p*\*). Thus, *L_i_* = 1 reverts to the first situation. All attracted fish who fail their Bernoulli trial are not considered for future matches.

So, in Step 1 we generate numbers of each species that are attracted to the fishing gear from (S.1), then in Step 2 generate the bite times of each fish of each species depending on the species’ aggressiveness. The bite times for all attracted fish are ordered sequentially and a Bernoulli trial is performed to see if the fish is captured by the gear in Step 3, with a probability that depends on whether the proportion of baits removed has exceeded the threshold *p*\*. This results in simulated catch counts, *c_itk_*, for each species and each fishing event.

#### S.1.4 Step 4 – fit the simulated data using three alternative methods

Having generated the data, we use three methods to fit the catch count data of the target species.

The first method does not account for hook competition, and is the CPUE Poisson-lognormal model from equations (S.1)-(S.3) that was used to generate the catch counts that would occur in the absence of hook competition. The response variables are the observed catch counts of the target species. Thus, in some sense, the model has been correctly specified, albeit with hook competition not considered and with only the lower-right diagonal element of **∑** estimated.

The second method uses a simple scaling to account for hook competition, and is here called the ICR (instantaneous catch rate) method. The method attempts to scale up the catch counts to account for the fish that would have been caught if all hooks had been actively fishing during the full fishing event. For example, if 5 fish of the target species were caught in an event, but only 200 hooks were returned with bait, the second method attempts to infer the number of fish of the target species that would have been caught if all 800 hooks had been actively fishing during the whole fishing event.

To do this, the ICR method multiplies the observed catch count by a competition adjustment factor, and then uses the same Poisson-lognormal model as for the CPUE method just discussed. The adjustment factor is derived and explained in Clark (2008), Webster and Stewart (2013), and Anderson *et al.* (2019), and it assumes that every fishing event with at least one catch is impacted by hook competition. It is applied to each fishing event, and is given by

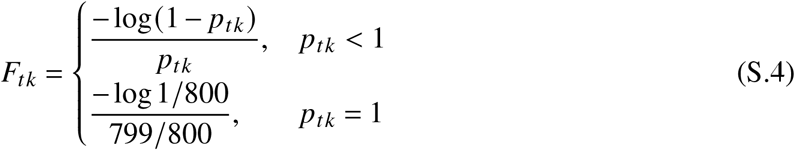

where *p_tk_* = *c._tk_/h_tk_* is the proportion of baits removed during fishing event *k* of year *t*. For each fishing event, the observed catch count of the target species is multiplied by the adjustment factor *F_tk_* and then rounded to the nearest integer, to give a competition-adjusted integer count. The first line of (S.4) is infinite when *p_tk_* = 1 (i.e. when all 800 baits are removed), and so we set its value to the value when 799 of the 800 baits are removed (Anderson *et al.*, 2019), giving the second line of (S.4) which is an adjustment factor of 6.69 for 800 hooks. The adjustment factor derivation involves assumptions on the instantaneous catch rates (ICR) of each species, and so the estimation method is an ICR-adjusted Poisson-lognormal model, hereafter called the ICR method.

The final method is our new censored method described in the main text. In many settings, the assumed censorship proportion for the target species, 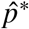, is deliberately mis-specified to not equal the true value *p*\* that was used to generate the data, to test the influence of such mis-specification (since 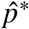 is not estimable within the model). For some settings with 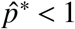, we use a conservative upper bound *u*_3*tk*_ for each fishing event to help improve numerical convergence (see later for derivation). No upper bound is used when 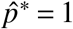. To further improve the convergence of the model, for each year *t* we investigate the effect of considering all observed catch counts *c*_3*tk*_ that exceed or equal the *Q^th^* quantile of the catch counts that year as ‘high-quality’, regardless of the value of *p_tk_*. Thus, the censorship interval is replaced with the original count *c*_3*tk*_ for these counts. We experiment with different values of *Q.*

After all models have been fit, the posterior median, posterior *2.5^th^* percentile, and the posterior 97.5^*th*^ percentile of each exp(*α*_3*t*_) for *t* > 1 are computed. Bias is calculated as the posterior median minus the true value, and the squared error is the bias squared. The coverage indicator variable is assigned value 1 uniquely for each exp(*α*_3*t*_) for *t* > 1, if the 95% credible interval contains the true value, and 0 if not.

Six experiments are performed, with each experiment (except 6) testing a different combination of settings 100 times, to give 100 estimates of the statistics mentioned in the previous paragraph. To summarise the performance of each method, the 100 bias, squared error, and coverage values are aggregated, with confidence intervals produced for each aggregate measure. The confidence intervals allow for levels of statistical significance to be attributed to the comparisons being made between the competing methods. For the bias and squared error values, the median and a robust 95% confidence intervals are computed to reduce the impact of outliers on summary measures of performance. Robust confidence intervals are computed using the median absolute deviation (MAD) measure of dispersion scaled by 1.48 to ensure consistency with standard confidence intervals (Maronna *et al.*, 2019). For the coverage indicators, the proportion of 1’s are computed along with an approximate 95% asymptotic confidence interval for the proportion computed using the normal approximation to the binomial proportion (i.e. ±2SE). For bias and median-squared error (MSE), a value close to zero is desired. For coverage, the desired value is 95%.

The different settings for each experiment are now given.

#### S.1.5 Experiment 1

For experiment 1 we use the following settings for a single-species scenario, with all combinations of the two choices of each of *μ*_3,*t*_ and *p*\* giving the four panels of results in Figure S.1:

- 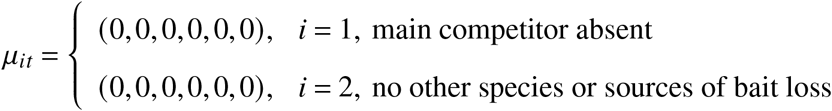
- 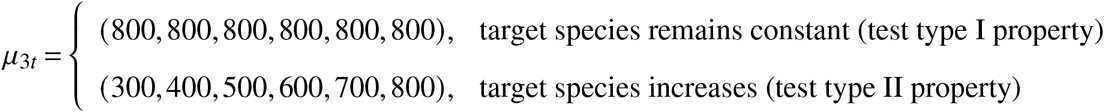
- *L*_3_ = 0.2, severity of hook competition (i.e. gear saturation effects) when 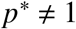
- **g**_3*rk*_ ~ U(0.5), uniform distribution of bite times
- 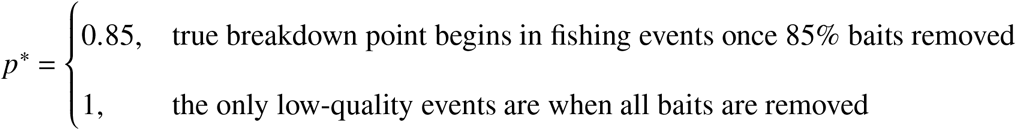
- 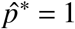, assumed breakdown point for fitting is not always the true value
- 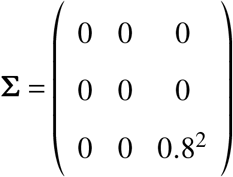, covariance matrix (only a single schooling 1007 species is present)
- *K_t_* = 30, number of stations sampled at per year
- 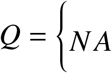, the largest catch counts can be censored

Thus, in experiment 1, there is only one species present, the target species. Very high levels of overdispersion and right-skewness are present in the target species’ count distribution (indicative of frequent schooling behaviours) due to the value of 0.8^2^ in **∑**.

When *μ*_3*t*_ = 800 for all years, the target species’ relative abundance does not change. This setting explicitly evaluates type I properties. This results in around 50% of the fishing events having all 800 baits removed. The purpose is to confirm that the censored method can correctly identify the lack of trend, regardless of whether attracted individuals are always captured if any baited hooks remain available (*p*\* = 1), or the probability of capture declines as less hooks are available (*p*\* = 0.85), and even when the assumed value of 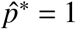 does not equal the true value of *p*\*. The target species’ count distribution is designed to reflect a species that often appears in group sizes that are large enough to saturate the fishing gear.

When *μ*_3*t*_ = (300,400,500,600,700,800) we are explicitly evaluating type II properties. The high levels of overdispersion results in all 800 baits being removed during many of the fishing events, including in the early years. The purposes of this experiment are: i) to verify the result of the theoretical example given in the main text, namely, that the CPUE index will underestimate the increase in the relative abundance due to the frequent gear saturation; ii) to verify that the censored method can still recover the true trend of a schooling species, despite the saturated fishing gear. Note that no upper bound on the species catch count is assumed (i.e. *u_itk_* = ∞ is assumed).

#### S.1.6 Experiment 2

In experiment 2 many species are caught, and characterized as the main competitor (*i* = 1), other (*i* = 2) and the target species (*i* = 3). The target species is non-schooling due its distribution not being overdispersed. The lack of overdispersion implies that counts of the target species have the highest possible signal-to-noise ratio which should result in the methods inferring the relative abundance trends with the greatest precision. Again, the relative abundance of the target species is fixed through time to assess the type I property and increased to assess the type II property.

However, in this experiment the target species faces fierce competition from the main competitor for the hooks. The main competitor is a schooling species with a strongly overdispersed distribution. Thus, the main competitor frequently appears in very large group sizes and can rapidly deplete the baits. Furthermore, the main competitor increases in abundance through time (either slowly or quickly) and is able to, on average, reach the baits ahead of the target species on average (when bite times have a mixed distribution). Lastly, the other species group consists of multiple species with differing behaviours and levels of overdispersion, but the mean abundance of the other species group is stationary. Specific settings are:

- 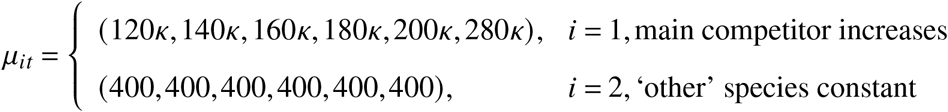
- 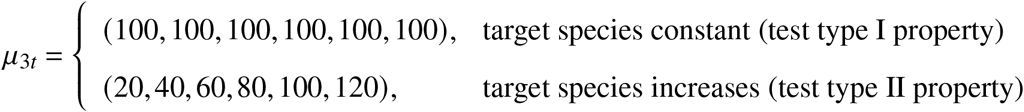
- 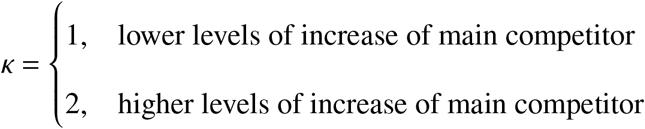
- (*L*_1_, *L*_2_, *L*_3_) = (1,0.8,0.2), gear saturation effects when 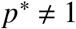
- 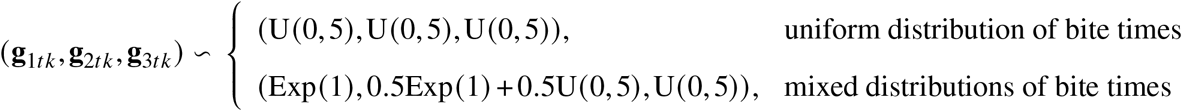
- 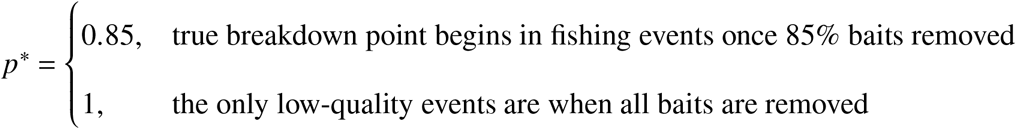
- 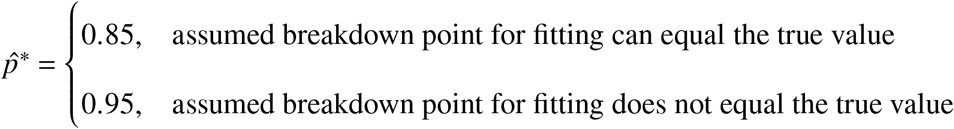
- 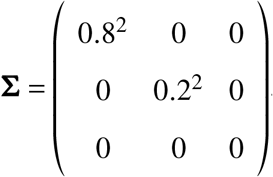, target species is non-schooling with two schooling competitors
- *K_t_* = 30, number of stations sampled at per year
- *Q* = *NA*, no restriction on which catch counts can be censored.

The four combinations of the two choices of each of *μ*_3,*t*_ and *κ* give the four simulated abundances in the leftmost column of Figure 1. The four combinations of the two choices of each of the breakdown points (*p*\*) and the bite rate distributions (the **g**’s) give the four columns of results of Figure 1. The CPUE and ICR methods, plus the Censored method with the two values of 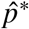 (which is only correctly specified when 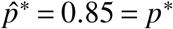) are tested, giving the four methods in the legend in Figure 1. Hence we have four different methods tested for each of the sixteen different simulation settings, giving 64 sets of results.

To summarize, interspecies hook competition (sometimes severe) is present between three species groups with independent distributions (i.e. no correlation is present due to the zero off-diagonal entries of **∑**). The target species in this experiment never appears in large schools, and (especially in the ‘mixed’ bite time distribution setting, also called ‘Differing Aggressiveness’ in the Figures) is sensitive to the competition for hooks from the other two species groups. The level of competition increases with time due to the increasing abundance of the main competitor. Note that a conservative upper bound, *u_itk_*, is placed on the species catch counts but is not believed to impact the results (derived and discussed later).

#### S.1.7 Experiment 3

Experiment 3 uses the same settings as experiment 2 except for:

- 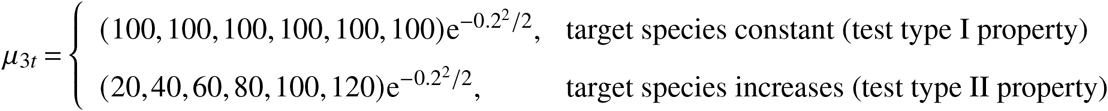
- (**g**_1*tk*_, **g**_2*tk*_, **g**_3*tk*_) ~ (Exp(1), 0.5Exp (1)+ 0.5U(0,5), U(0,5)), mixed distribution of bite rates
- *p*\* = 0.85, true breakdown point in simulations
- 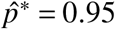, the breakdown point is always mis-specified
- 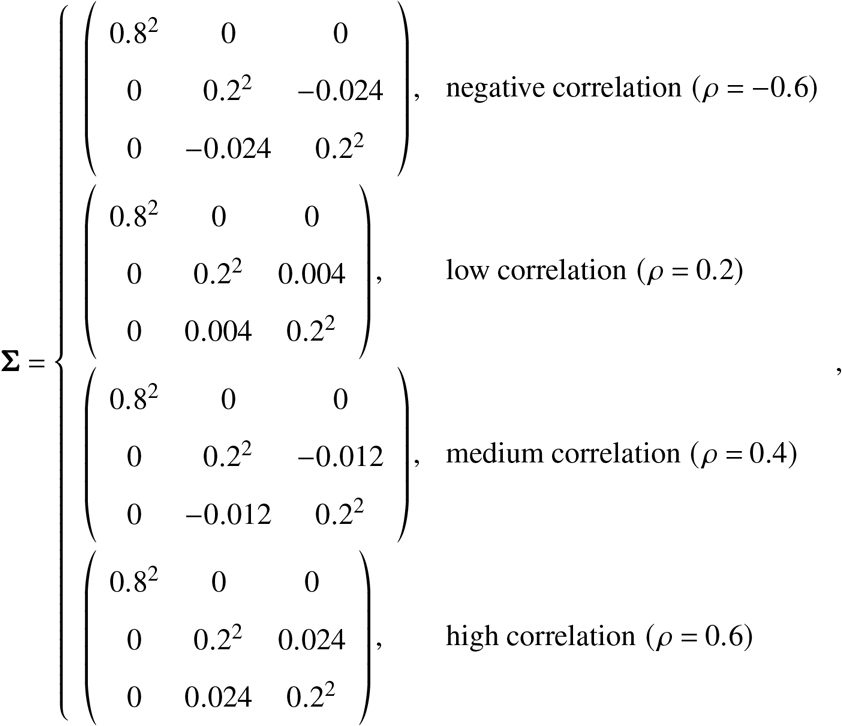
- *K_t_* = 100, an increased number of stations sampled at per year
- 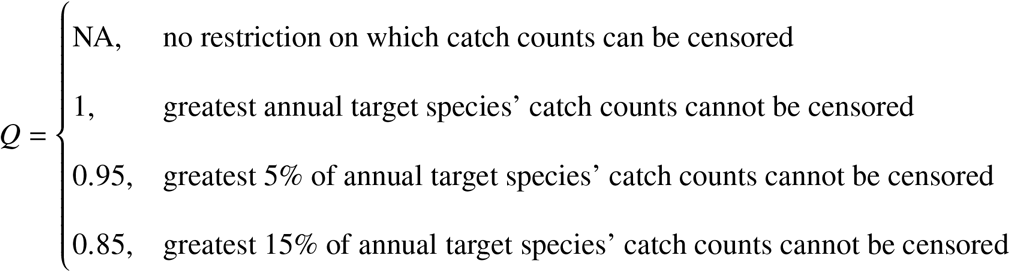

Thus, experiment 3 allows the count distribution of the target species to be overdispersed and correlated with the count distribution of the ‘other’ species group. Note that we scale the mean catch count of the target species *μ*_3,*t*_ by e^-0.2^2^/2^ to account for the bias due to overdispersion and to ensure the expected catch counts match experiment 2. The target species is highly sensitive to the competition for hooks from the other species, typically reaching baits later in the soak times of fishing events. More stations are sampled because: i) the extra overdispersion and hence the schooling behaviours now exhibited by the target species reduces the signal-to-noise ratio in the target species distribution; ii) the extra correlations between the target and the other species group’s distributions induces an additional stochastic dependency between the probability that a fishing event will be censored and the expected catch count of the target species. Only the results of *Q* = NA and *Q* = 0.85 are shown since the performance for the other two values across all metrics lies in-between. The combination of settings gives the results in Figure 3. Note that a conservative upper bound, *u_itk_*, is placed on the species catch counts but is not believed to impact the results (derived and discussed later).

#### S.1.8 Experiment 4

For experiment 4 we repeat experiment 2, but with:

- *p*\* = 0.5, much lower true breakdown point in simulations
- 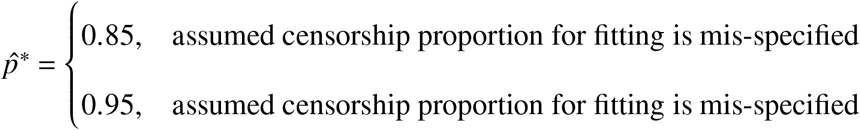

Thus we still specify 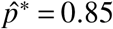 and 0.95 as in experiment 2, but neither are close to the true value of *p*\* = 0.5.

#### S.1.9 Experiment 5

For experiment 5 we repeat experiment 3 but with:

- (**g**_1*tk*_, **g**_2*tk*_, **g**_3*tk*_) ~ (U(0,5),0.5Exp(1)+ 0.5U(0, 5),Exp(1)), the target species is now fastest
- 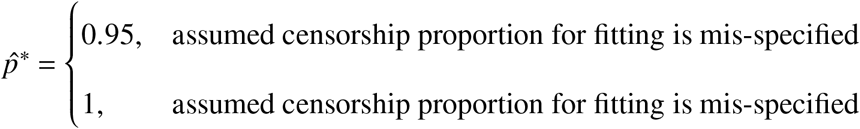

The target species typically reaches the baits ahead of the competing species. The purposes of this experiment are to: i) empirically confirm the results of the theoretical example presented in the main text, namely, that the ICR method will not satisfy the type I setting due to the insensitivity of the target species to inter-species hook competition; ii) investigate the robustness of the censored method when the target species is not strongly affected by hook competition, nor appears in large enough numbers to saturate the fishing gear. Put differently, experiment 5 considers a setting where a target species is largely ‘immune’ to hook competition in the data collected. Note that a conservative upper bound, *μ_itk_*, is placed on the species catch counts when 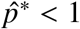 but is not believed to impact the results (discussed and derived later).

#### S.1.10 Experiment 6

Exploratory analysis of the simulated observed catch counts *c_itk_* (after the bite times have been matched to the next available hook) versus *p_tk_* for the target species of experiments 2-4 appears to show that the mean of *c_itk_* declines rapidly after *p_tk_* exceeds 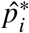. This trend appears to be consistent across different choices of 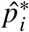. Conversely, a rapidly increasing and negligible trend is seen between *c_itk_* and *p_tk_* for the main competitor of experiments 2-4 and the immune species of experiment 5, respectively.

Experiment 6 formally investigates if these trends between *c_tk_* and *p_tk_* persist across very large numbers of replications and changing settings. If so, this would imply that exploratory analysis of *c_tk_* vesus *p_tk_* in real longline data can help the scientist: i) select a suitable value of 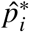 as the value where the trend begins; ii) understand the statistical properties of the estimates, most notably helping the scientist to understand the levels of trust they can assign the reported levels of uncertainty based on the findings from experiments 1-5.

We simulate data (scenarios described below) with *p*\* = 0.85 and fit the following model to each simulated dataset (values of *c*_3*tk*_ and *p_tk_*):

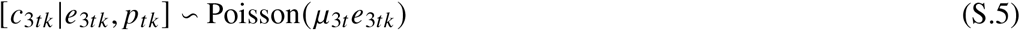

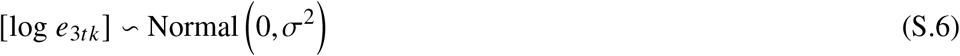

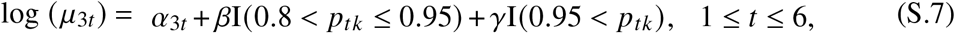

where *σ* is an estimated standard deviation, I(·) is the indicator function (1 if the condition is satisfied, 0 if not), *α*_3*t*_ + *β* represents the log of the mean catch counts observed at ‘medium’ values of *p_tk_*, and *α*_3*t*_ + *γ* the log of the mean catch counts observed at ‘high’ values. We use these discrete values (rather than continuous) here for simplicity, but allow continuous in the application to real data described in the case study.

We define these intervals (0.8 < *p_tk_* ≤ 0.95 and 0.95 < *p_tk_*) to see if we can detect a change in the mean observed catch count when *p_tk_* ≈ 0.85 (the known breakpoint is *p*\* = 0.85) to when *p_tk_* ≈ 1. These values were also chosen based on the exploratory plots. Then, the target for inference is exp(*γ* – *β*), the relative value of the mean catch counts observed at high and medium levels of *p_tk_*. By plotting estimates of exp (*γ* – *β*) across a wide range of simulation settings, we aim to develop a method for choosing 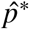 for real data. We repeat all combinations of the following settings 20 times:

- 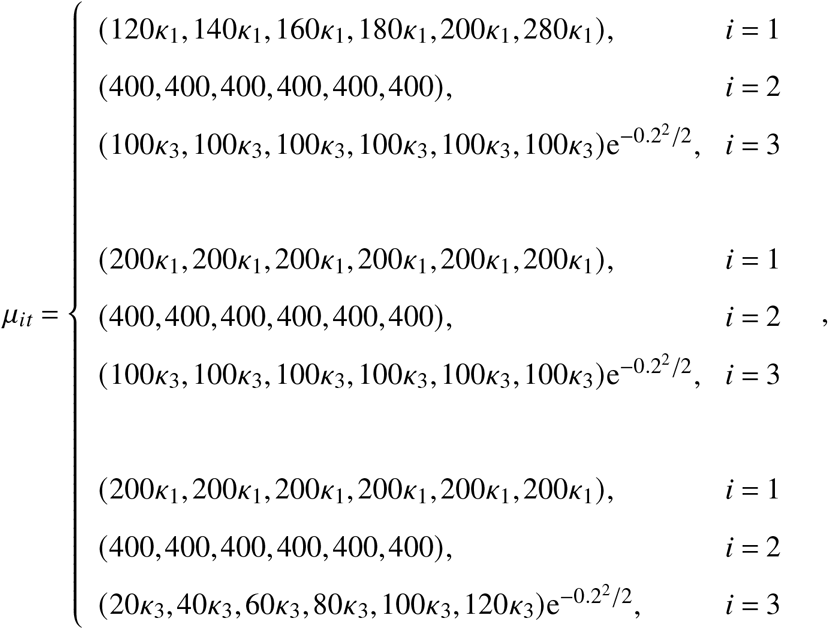
- 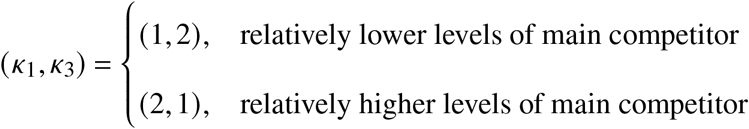
- (*L*_1_, *L*_2_, *L*_3_) = (1,0.8,0.2), saturation effects when 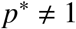
- 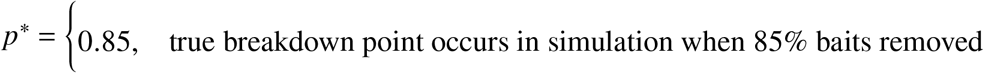
- 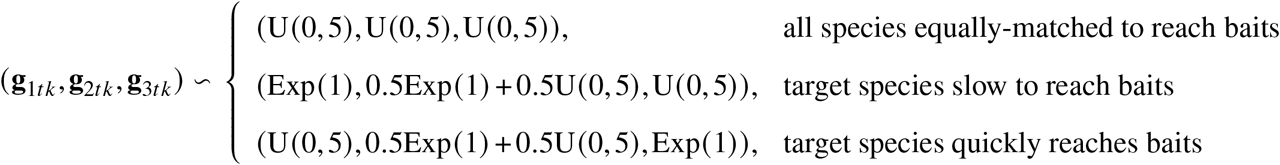
- 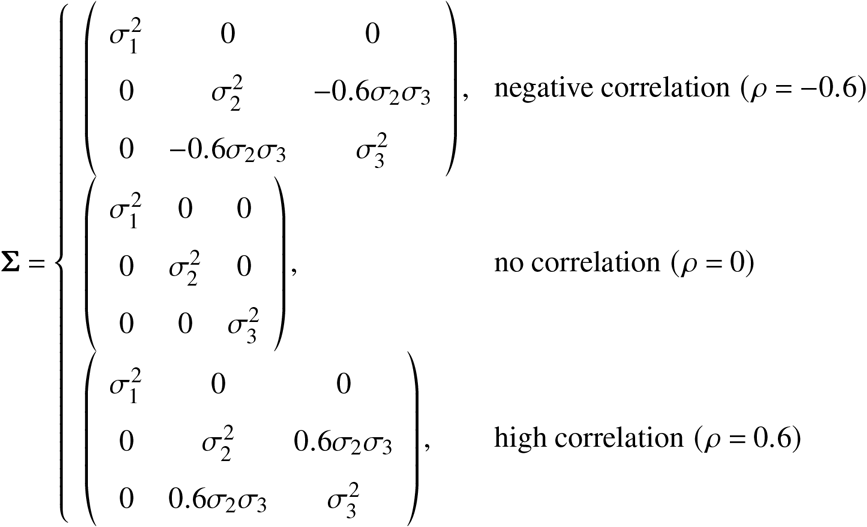
- 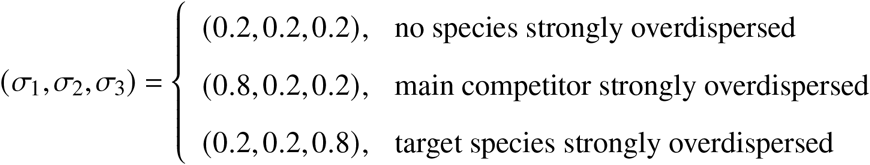
- *K_t_* = 100, high number of stations sampled at per year

Finally, we produce boxplots of estimates of exp(*γ* – *β*) across all the settings, but grouped by the relative speeds of each species group to the baits (i.e. the 3 **g**_*itk*_ settings) and the relative degree of schooling exhibited by the three species groups (i.e. the 3 *σ_i_* settings). See Figure 5 in the main text for results.

#### S.1.11 Fitting details for all experiments and additional results

For all experiments, we use the R-INLA package for model fitting (Rue *et al.*, 2009; Lindgren *et al.*, 2011; Lindgren and Rue, 2015). We place a flat prior on *α*_3,1_ and a weakly-informative normal prior with mean 0 and variance 1000 on the remaining *α*_3,*t*_ for *t* > 1. Lastly, we use R-INLA’s default Gamma prior with shape 1 and scale 5 × 10^-5^ for the precision of the lognormal distribution describing the *e*_3*t,k*_ terms. To derive the posterior median and posterior quantiles used for the model comparison, we generate 5000 approximately independent Monte Carlo samples of *α*_3,*t*_ from the (integrated nested Laplace approximated) posterior distribution, exponentiate each sampled value, and then compute the necessary functions.

For experiment 1, Figures S.1, S.2, and S.3 present the bias, MSE, and coverage respectively. Only the censored method satisfies both the type I and type II properties. In particular, the CPUE and ICR methods perform poorly across all three metrics when the abundance of the single species increases over time. In the type I settings, the coverage levels of the censored method fall to 75%. The reason for this is currently unknown, however we do not see this is a cause for concern given the very low MSE values.

For experiment 2, Figures 1, S.4, and 2 present the bias, MSE, and coverage respectively. Results demonstrate the improved performance of the censored method, with: i) up to 100-fold reductions in MSE observed in both type I and type II settings; ii) very low bias across all settings; iii) coverage levels close to 95% in all settings. Note that the other two methods perform poorly, coverage often drops to 0% in both type I and type II settings and the bias in estimates can be extreme.

For experiment 3, conclusions are similar to experiment 2 and the censored method performs best across all metrics. Figures 3, S.5, and 4 present the bias, MSE, and coverage respectively. For the censored method, coverage levels close to 95% are attained in all type I settings and they always exceed 50% in the type II settings. Coverage levels from the other two methods are close to 0% in many settings. A moderate deterioration in the performance of the censored method occurs in settings where the target species’ abundance distribution is positively correlated with the abundance distributions of competing species. Figure S.5 shows an increase in MSE as the correlation increases. The quantile-modification has only a small impact on the statistical properties of the censored method but ensures convergence (Figure S.6).

For experiment 4, conclusions are similar to experiment 2, despite the badly mis-specified 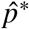 value. Figures S.7, S.8, and S.9 present the bias, MSE, and coverage respectively. They show the bias and MSE from the grossly mis-specified censored method to be at least as good as the competing methods across all settings tested, and far better in most settings. In summary, results show the censored method retains improved statistical properties when 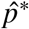 is poorly chosen.

For experiment 5, Figures S.10, S.11, and S.12 present the bias, MSE, and coverage respectively. They confirm that the CPUE method yields values that closely follow the true abundance trends of a target species when it is unlikely to be subject to hook competition. When absolute fish numbers are low, the CPUE method method performs best with respect to MSE and coverage. However, this performance improvement is lost as fish numbers increase (increasing the proportion of baits removed) and the censored method outperforms across all metrics once again. In fact, coverage levels of both the ICR and CPUE methods fall close to 0% in both type I and type II settings when the total fish density is high. Unfortunately, however, the coverage of censored method can fall as low as 40% and MSE values can remain high. Note that the ICR method performs poorly in all settings across all metrics, in agreement with the earlier theoretical example – the ICR method is not well suited for species that can typically reach baited hooks early on in the soak times since it assumes that hook competition impacts every fishing event.

For experiment 6, cross-correlating the results shown in Figure 5 with those from experiments 1-5 suggest a simple empirical test for assessing if the censored method is suitable for modelling a given species. The censored method performed very strongly for the target species of experiments 2-4, for which Figure 5 presents a decreasing abundance trend with *p_tk_*. The censored method also performed well for experiment 1, for which Figure 5 presents a strong increasing abundance trend with *p_tk_* for cases where the target species are responsible for frequently saturating the fishing gear. Finally, the censored method performed adequately for the ‘immune’ species of experiment 5, for which Figure 5 presents a weak abundance trend with *p_tk_* for the target species in this experiment.

Thus, the censored method should perform very well for species exhibiting strong declines in their observed catch counts *c_tk_* at ‘high’ values of *p_tk_*. This negative trend should remain after controlling for the effects of exogenous factors (e.g. year). A statistically significant decline in mean catch counts exceeding 10% appears to be a conservative decision rule; whilst the censored method was found to perform very well in many settings that fail this decision rule, the rule appears to filter out problematic ‘immune’ species seen in experiment 5. By comparing the ‘best-performing’ 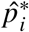 values in experiments 2-4 with the exploratory analysis between *c_itk_* and *p_tk_*, we recommend that for species exhibiting a 10% decline, a value of 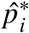 be chosen close to the value of *ptk* where the negative trend first begins.

Similarly, the censored method should perform well for species exhibiting large increases in their observed catch counts *c_itk_* at ‘high’ values of *p_tk_*. A statistically significant increase in mean catch counts exceeding 20% appears to be a conservative decision rule for confidently classifying a schooling species responsible for frequently saturating the longline fishing gear. Results from experiment 1 suggest setting 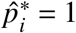, however uncertainty levels may be somewhat underestimated.

For species exhibiting neither a strong increase or decrease in mean catch count at ‘high’ values of *p_tk_*, results from experiment 5 suggest the censored method should be cautiously used with 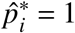. The user should be aware that the species could be largely immune to hook competition in the dataset. Hence, levels of uncertainty could be underestimated (due to the lower coverage levels seen in immune species). Furthermore, setting 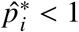 for an immune species can hurt the performance of the censored method.

For experiments 2 and 4 there were a few simulation iterations where the censored method failed to converge. When this happened, we simply repeated the simulation iteration (i.e. generated new data and re-fit all models) and ignored the convergence failure. In experiment 3, the convergence issues were much more prevalent due to the lower signal-to-noise ratio and were a target of research. Thus, for experiment 3, we recorded in the output instances where a model was reported by R-INLA as failing to converge.

Across all experiments, between 30-60% of fishing events returned <5% of the baits, and between 20-50% of events returned 0% (see Figure S.6). Interestingly, during experiment 3, convergence issues were more common when the all-species density was lowest. Over 40% of the iterations failed to converge in the most challenging settings if the censored method was used without adjustment. Implementing the quantile adjustment improved things. This adjustment considers all catch counts *c_itk_* exceeding the *Q^th^* annual quantile as high quality, regardless of the value of *p_tk_*. Using *Q* = 0.95 led to almost 100% convergence across all simulation settings, with *Q* = 0.85 ensuring 100% convergence throughout. We hypothesize the improved convergence is due to an improved ability to identify the variance of the overdispersion terms, *σ*^2^, since the right-tail of the catch count distribution is (artificially) observed.

Note that after conducting the simulation experiments, we discovered that convergence could be greatly improved by starting the numerical optimization with both initial values and Laplace approximations started at the final converged values found in the CPUE model. We have not repeated the simulation study with this change implemented due to the simulation experiments taking a very long time to fit, but note that the 40% estimate should be viewed as a highly conservative upper bound - 10 out of the 11 models in the case study fit without convergence issues and without requiring the use of any upper bound *u*_3,*t,k*_ (including the one described later), or requiring the quantile adjustment (i.e. *Q* ≤ 1 for 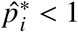).

**Figure S.1:**
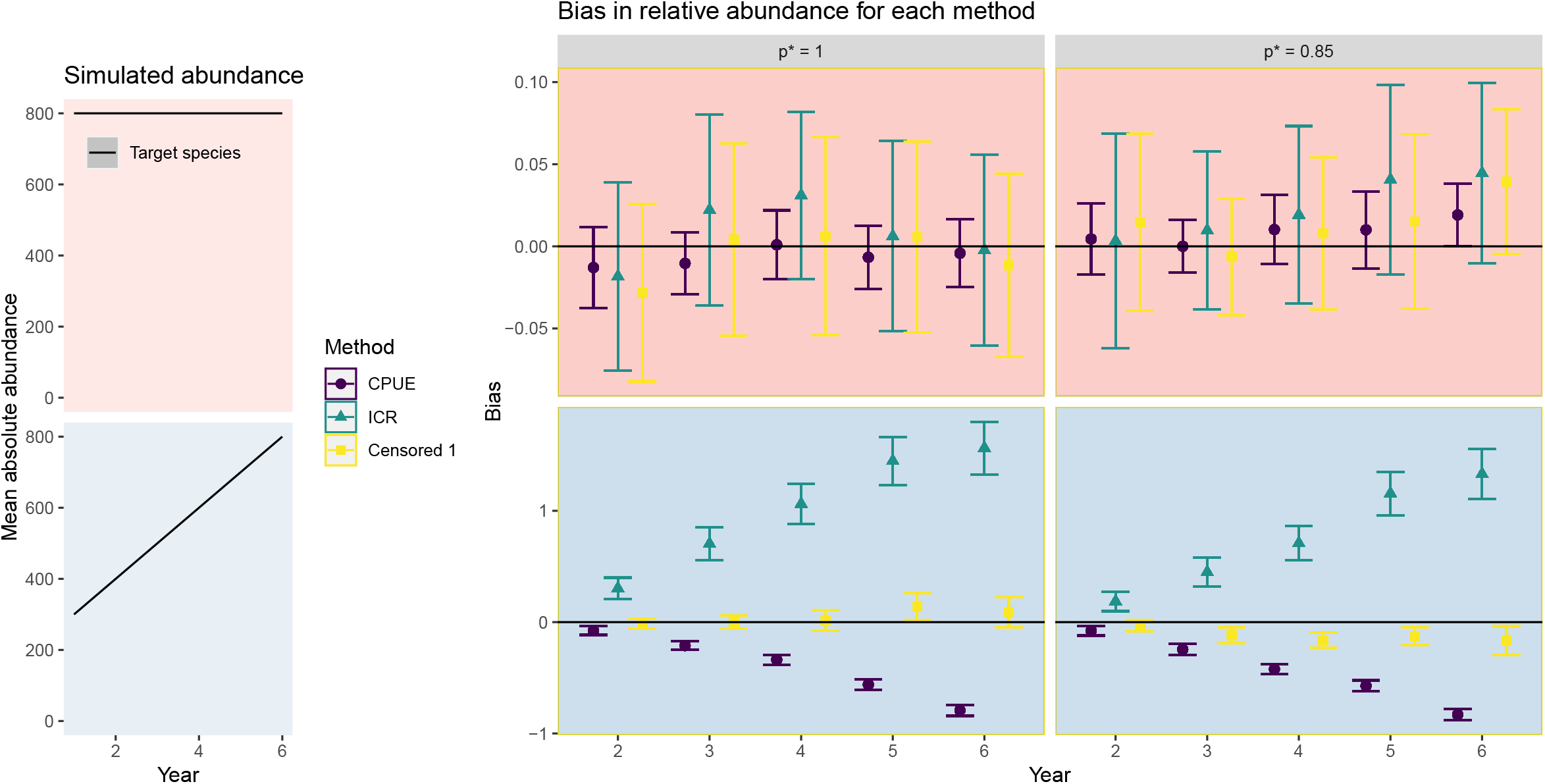
Bias in the relative indices across three methods for experiment 1. The leftmost column shows the mean abundance in the target species across time, such that the first row is testing the type I properties and the second row the type II. Columns specify if hook competition occurs only when all baits are removed (*p*\* = 1) or begins when 85% are removed (*p*\* = 0.85). Plotted are the median and robust 95% confidence intervals calculated from the 100 repetitions. The ‘Censored 1’ results correspond to the censored method with 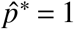. The bias from the censored method remains low in all settings, unlike the other methods which become severely biased in the type II setting.

**Figure S.2:**
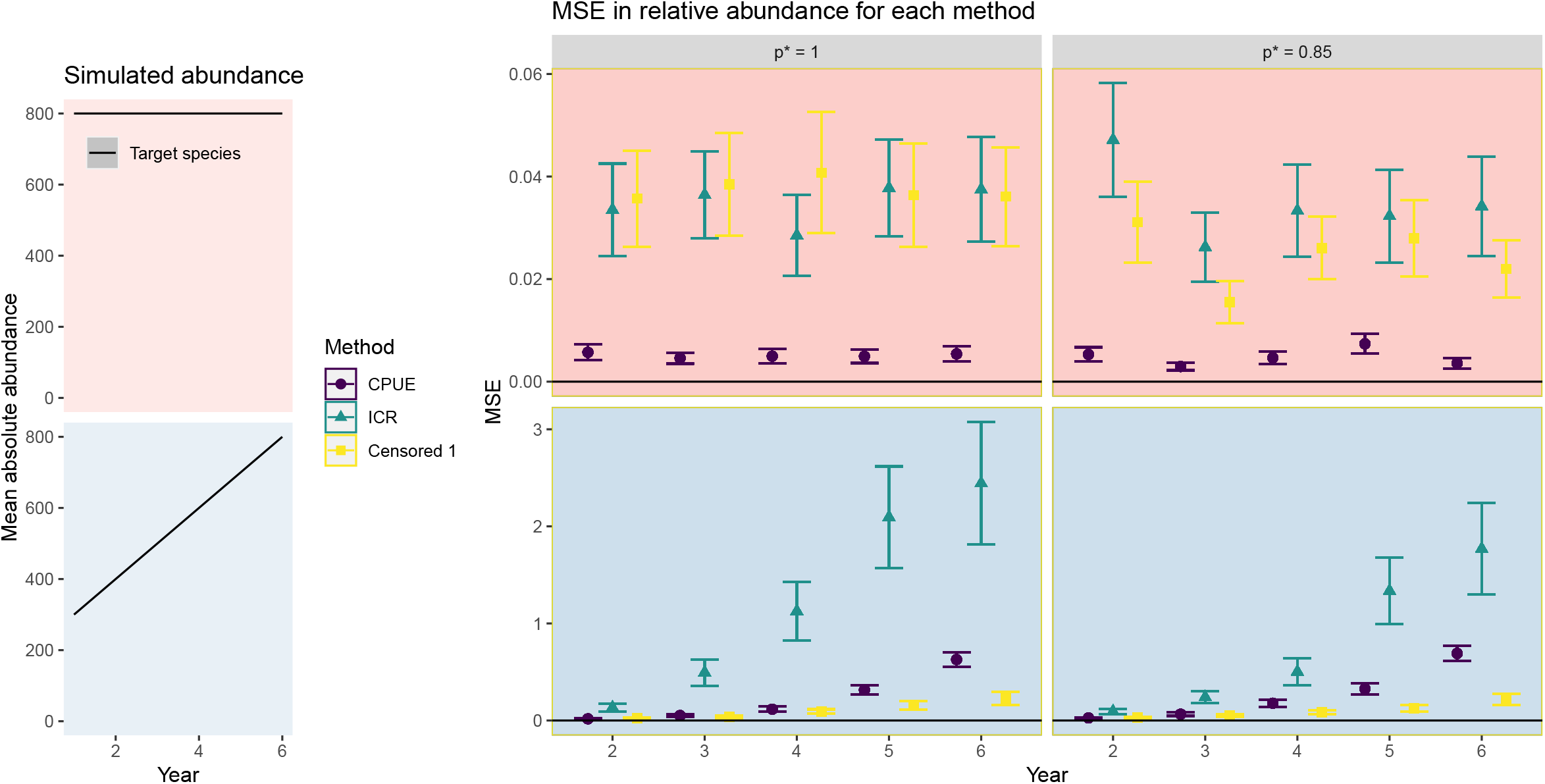
Median-squared error results for experiment 1. Rows and columns as for Figure S.1. Plotted are the median and robust 95% confidence intervals calculated from the 100 repetitions. The errors from the censored method remain low in type I settings (though not as low as the CPUE method) and are far smaller than the CPUE and ICR methods in the type II settings.

**Figure S.3:**
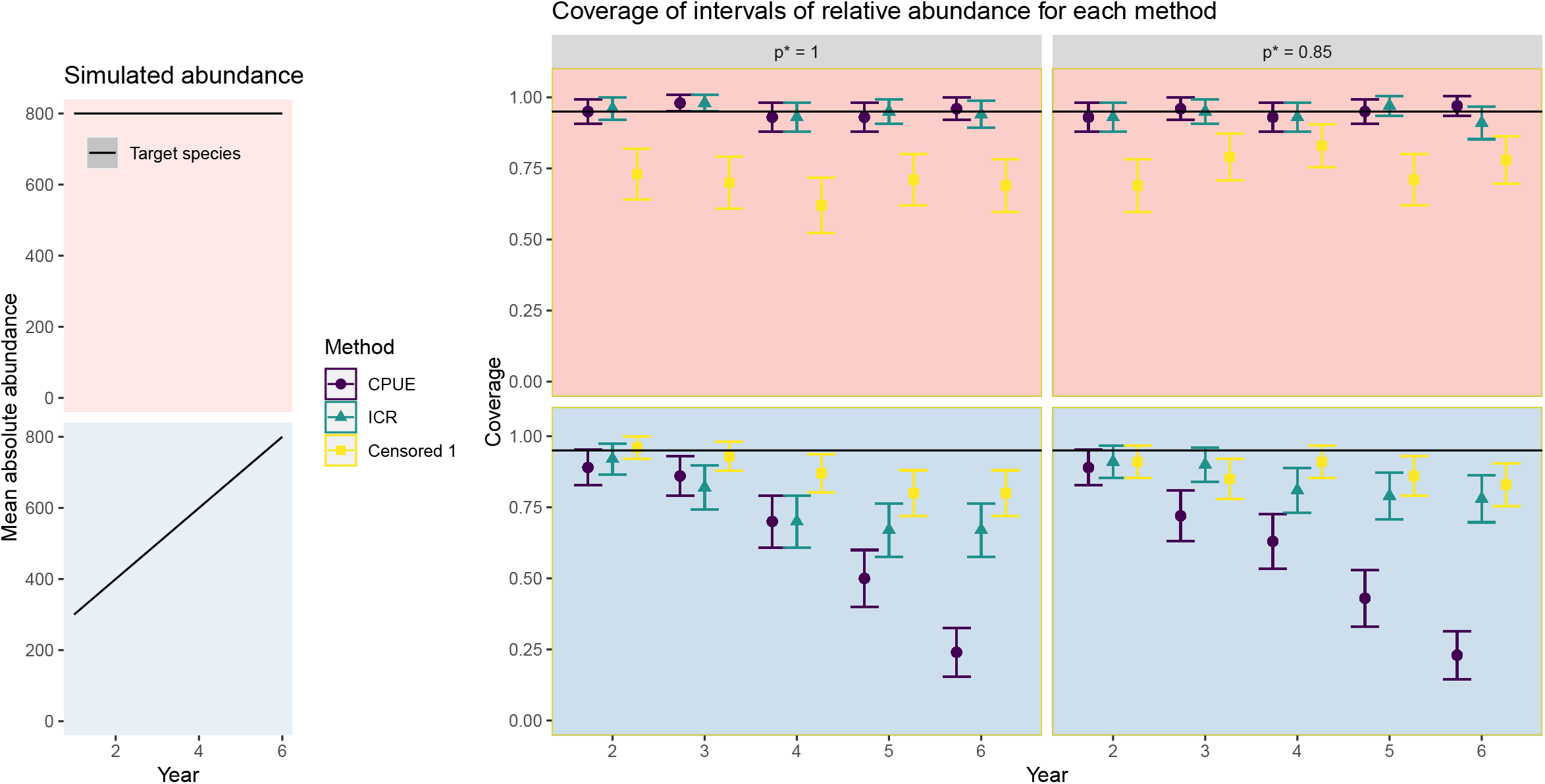
Coverage results for experiment 1. Rows and columns as for Figure S.1. Plotted are the mean and approximate 95% confidence intervals across the 100 repetitions. The coverage levels of the censored method remain high in all settings, however, can fall to ~70% in the type I setting.

**Figure S.4:**
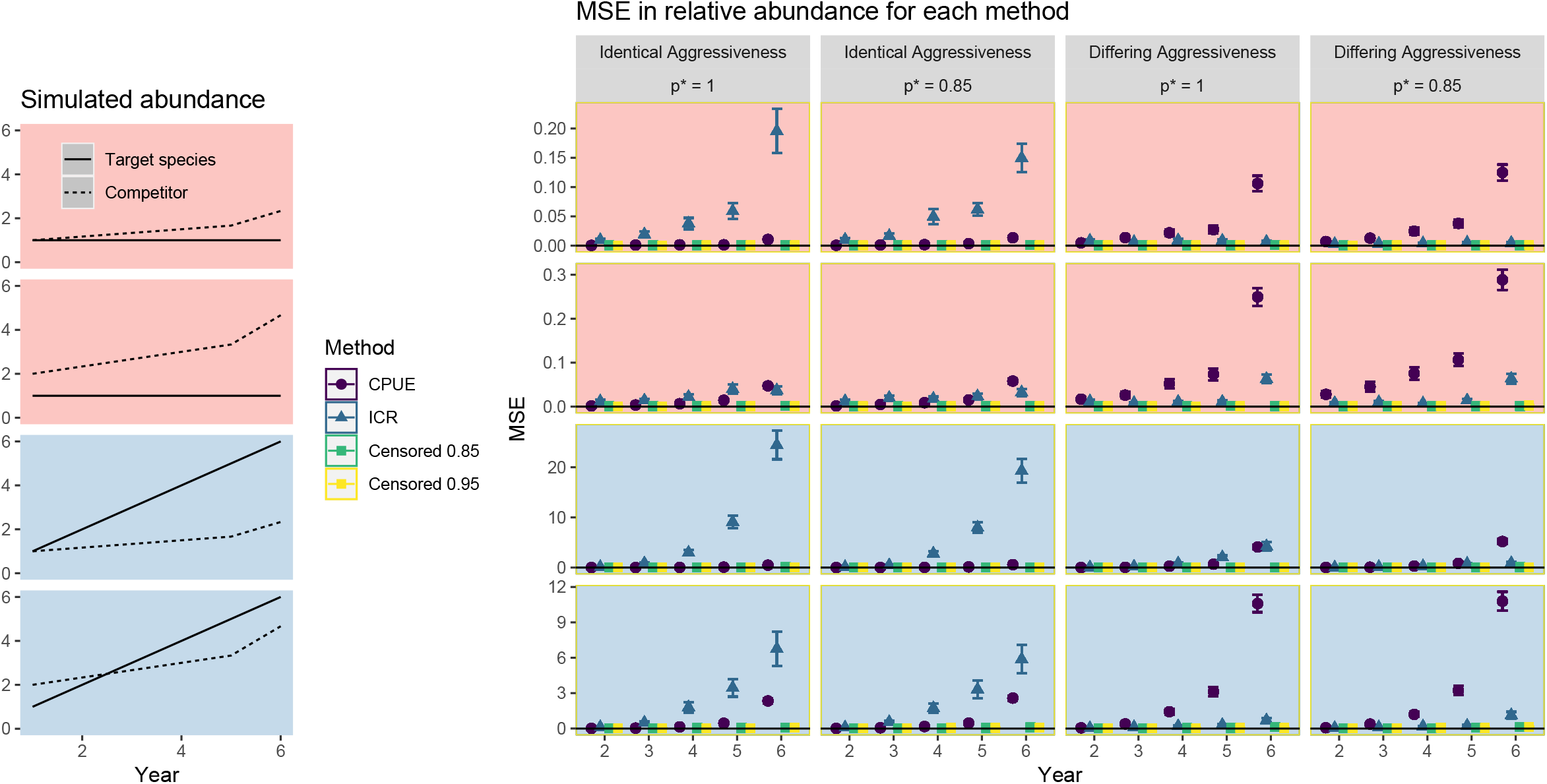
The median-squared error of the relative abundance indices for experiment 2. Columns and rows as in Figure 1. Plotted are the median and robust 95% confidence intervals computed from the 100 repetitions. The ‘Censored 0.85’ and ‘Censored 0.95’ methods assume 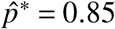 and 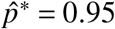, respectively. The censored method performs better than the others across all setting.

**Figure S.5:**
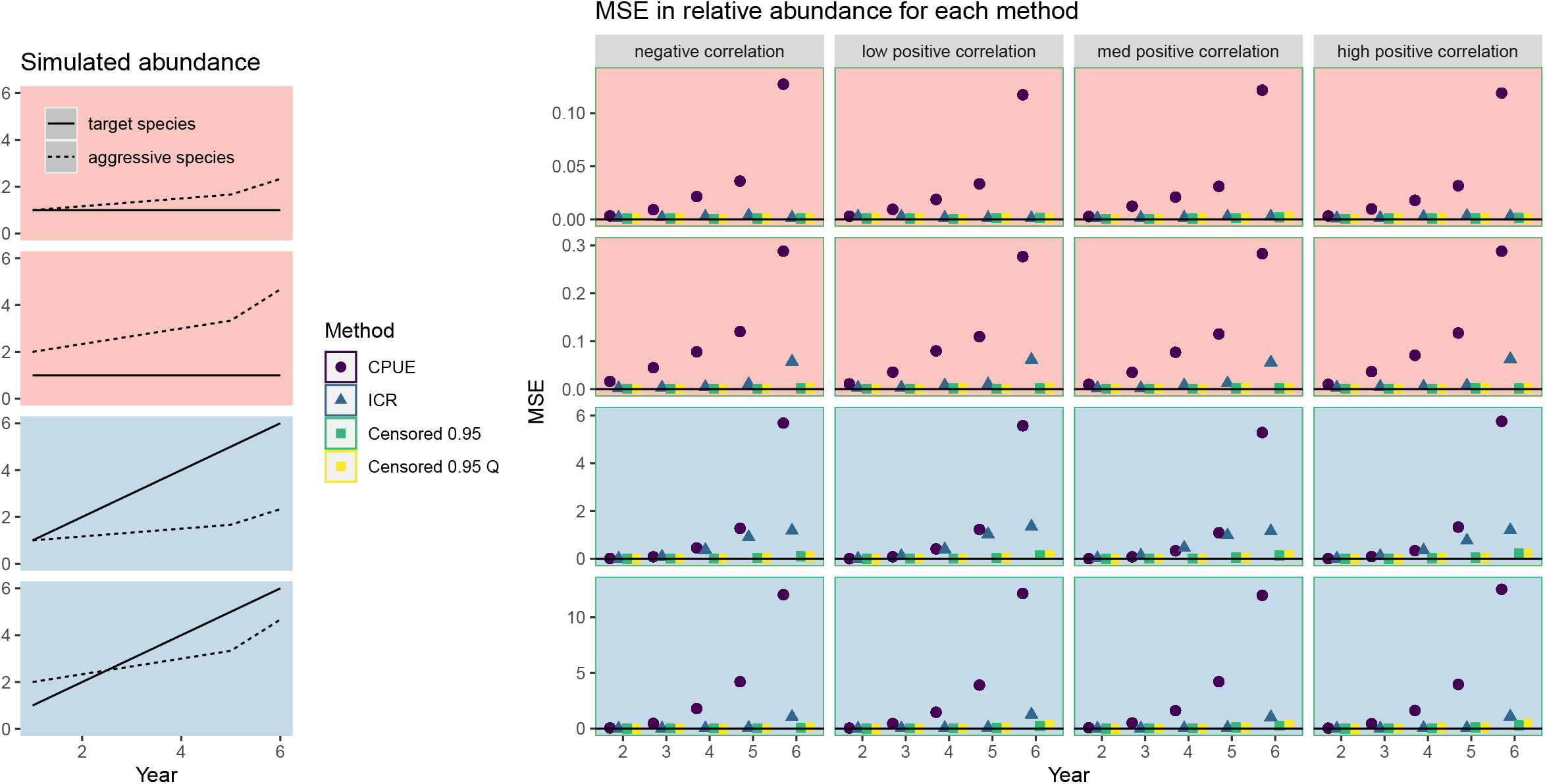
The median-squared error of the relative abundance indices for experiment 3. Rows, columns, and methods as in Figure 3. Also included are results from the ‘Censored 0.95 Q’ estimator, denoting the censored method where all observed counts of the target species ≥ the 85^*th*^ observed annual quantile are considered ‘high-quality’, regardless of *p_tk_*. Plotted are the median values. The approximate 95% intervals are omitted since they are typically smaller than the plotting character. The censored methods better estimates the relative abundance across all settings compared to the CPUE and ICR-base methods. Little difference between the two censored methods can be seen. Errors for the CPUE method are really high in all settings and errors for the ICR method are highest in the settings with the greatest levels of bait removal by all sources.

**Figure S.6:**
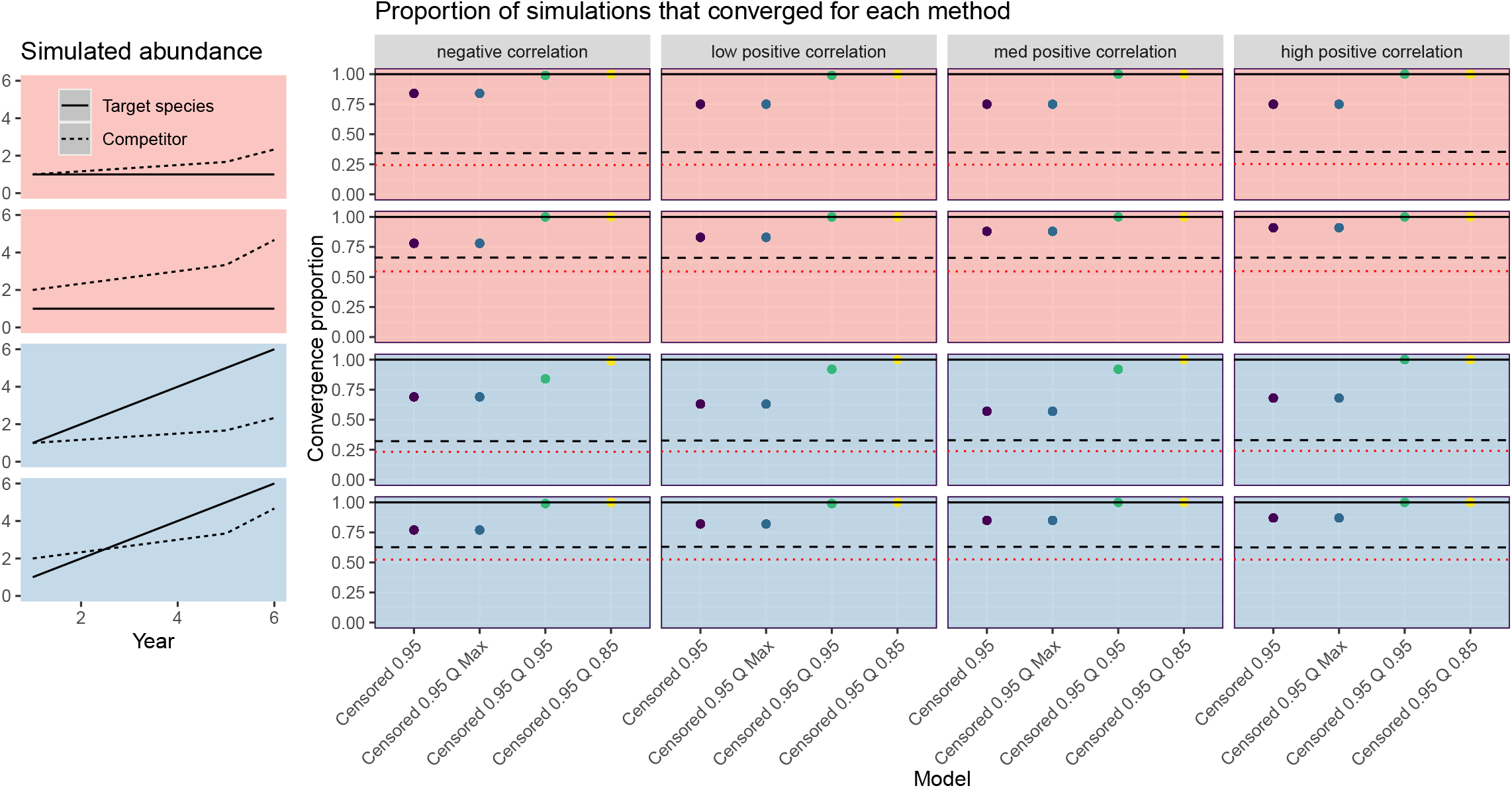
The proportion of the 100 simulations that successfully converged for the four variants of the censored method tested in experiment 3 with *p*\* = 0.85 throughout. We omit the CPUE and ICR methods as they always converge. Columns and rows as in Figure S.5. Plotted as points are the convergence proportions. Also plotted as a dashed black line and a dotted red line are the proportions of fishing events across the 6 years that experienced at least 95% and 100% bait removal. ‘Censored Q X’ denotes the censored method where all observed counts of the target species ≥ the *X^th^* observed annual quantile are considered ‘high-quality’, regardless of *p_tk_*. The ‘Censored 0.95 Q 0.85’ method converged in all settings but the ‘Censored 0.95’ method without the quantile adjustment failed to converge up to 45% of the time in certain settings. Over 60% of fishing events in rows 2 and 4 return 5% or fewer of the baits, and over 50% of events return no baits. In rows 1 and 3 these numbers are 20% and 30%. Note that convergence is now much higher in light of the improved optimization approach discovered after conducting this simulation study.

**Figure S.7:**
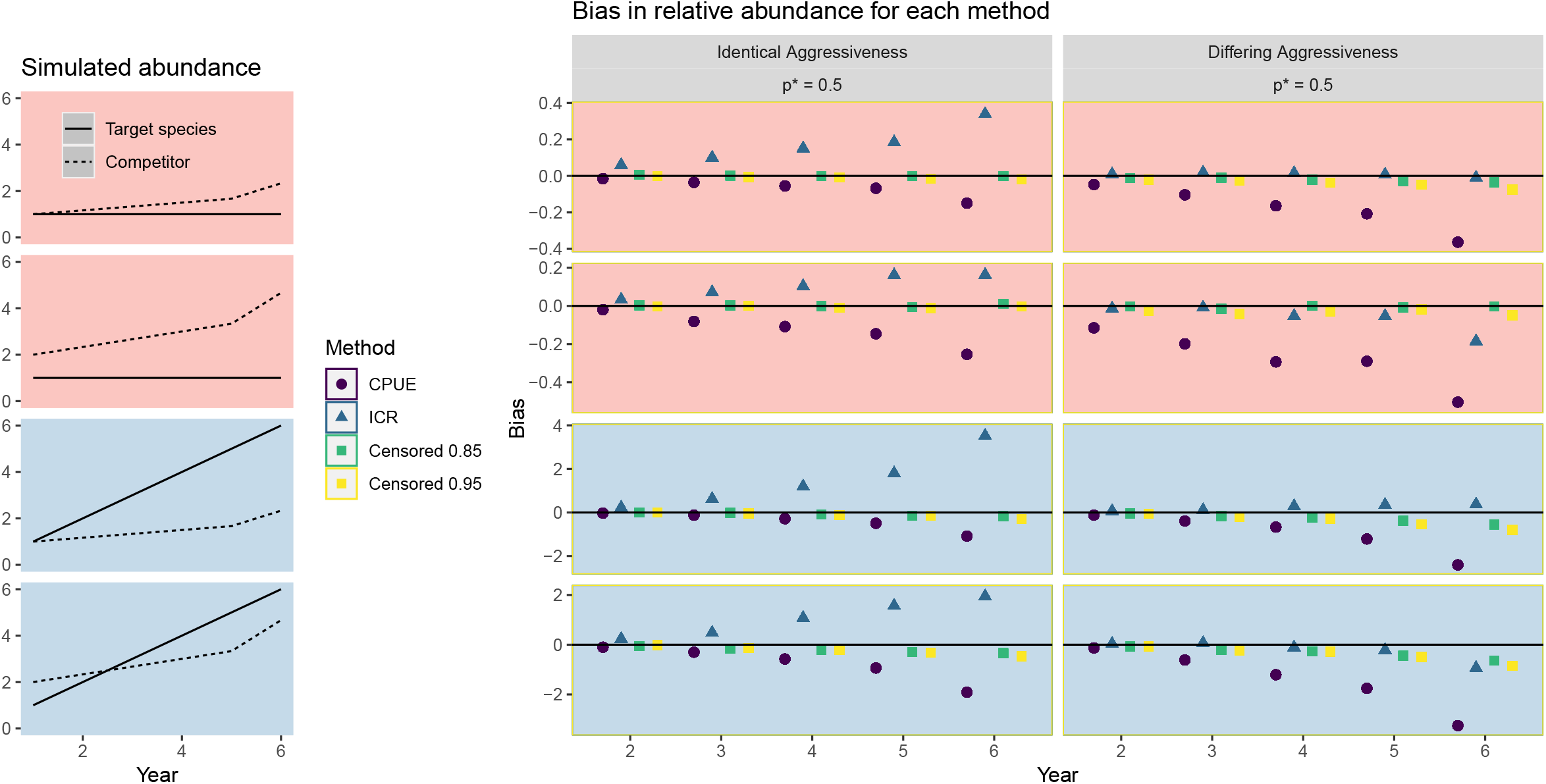
Bias results for experiment 4. Rows are defined as in Figure 1, with the two columns testing the alternative bite-time scenarios. Plotted are the median values. The approximate 95% intervals are omitted since they are typically smaller than the plotting character. The censored method typically outperforms the other two.

**Figure S.8:**
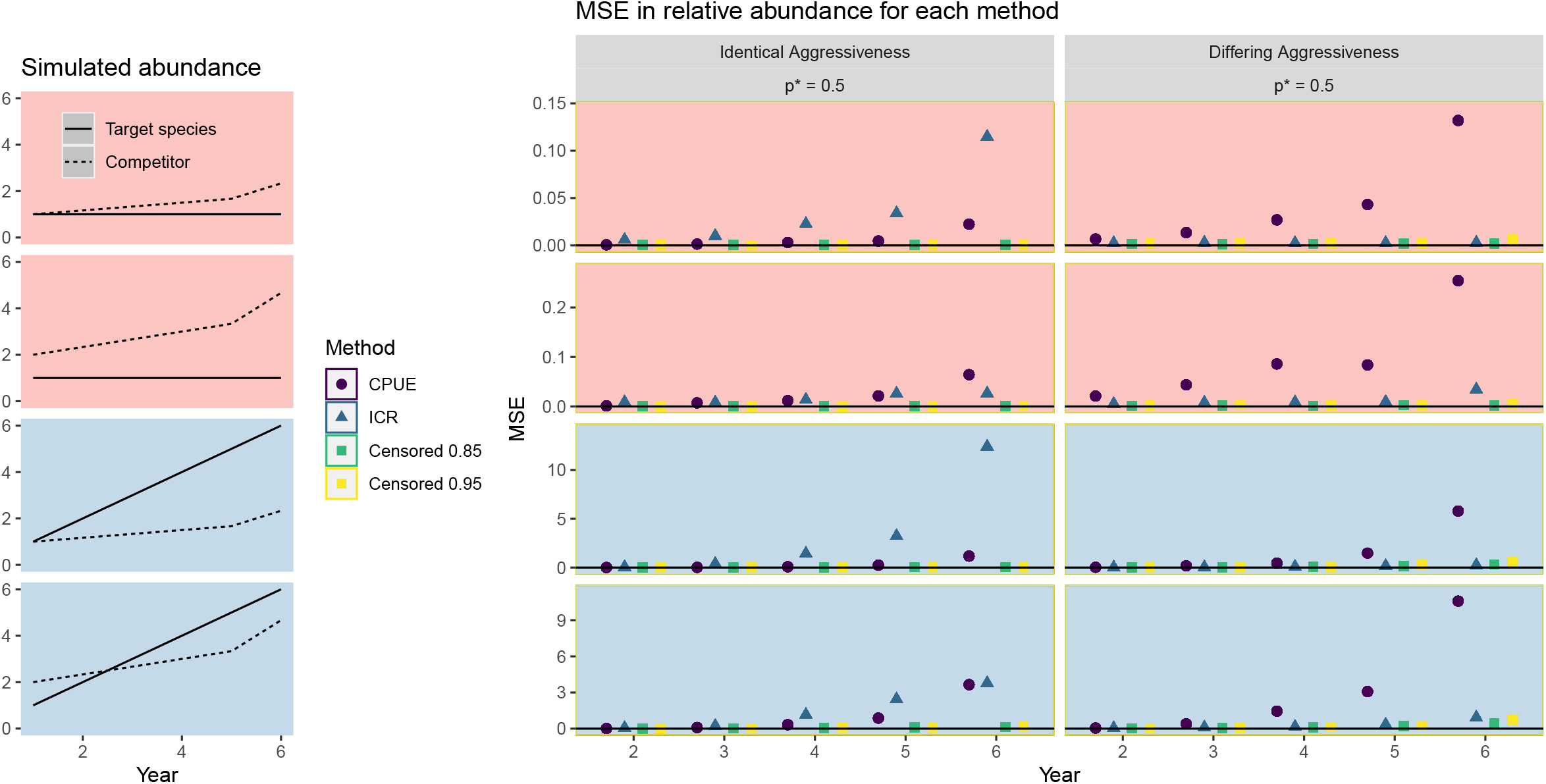
Median-squared error results for experiment 4. Details as for Figure S.7. The MSE is always lowest for the censored method by year 6.

**Figure S.9:**
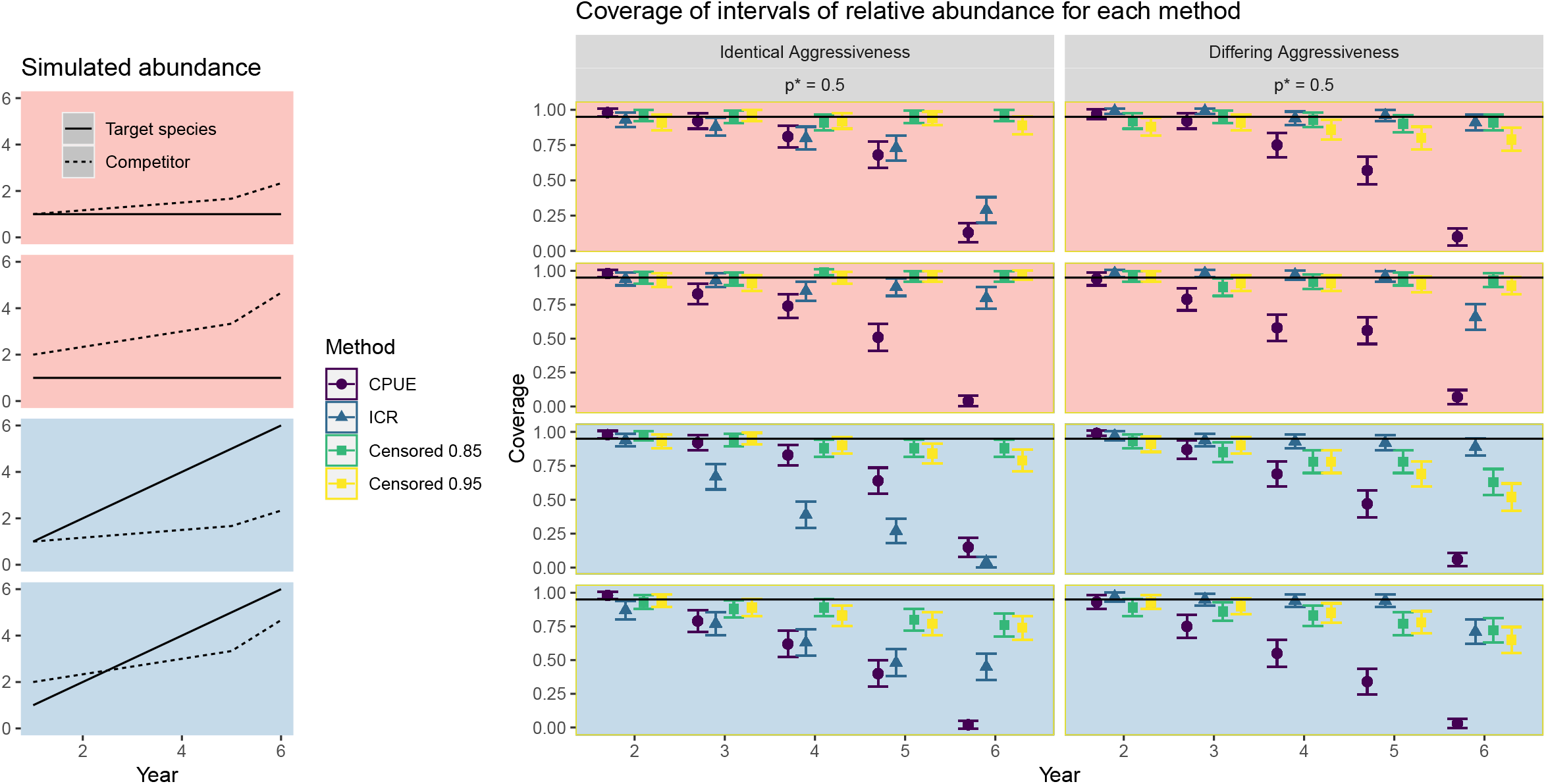
Nominal coverage of 95% credible intervals for experiment 4, for which *p*\* = 0.5 is fixed (way below the assumed 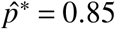 and 0.95 values used for the ‘Censored 0.85’ and ‘Censored 0.95’ methods). Details as for Figure S.7. The censored method attains the best coverage levels in all settings and close to 95% in the first two rows, satisfying the type I property but not always the type II property.

**Figure S.10:**
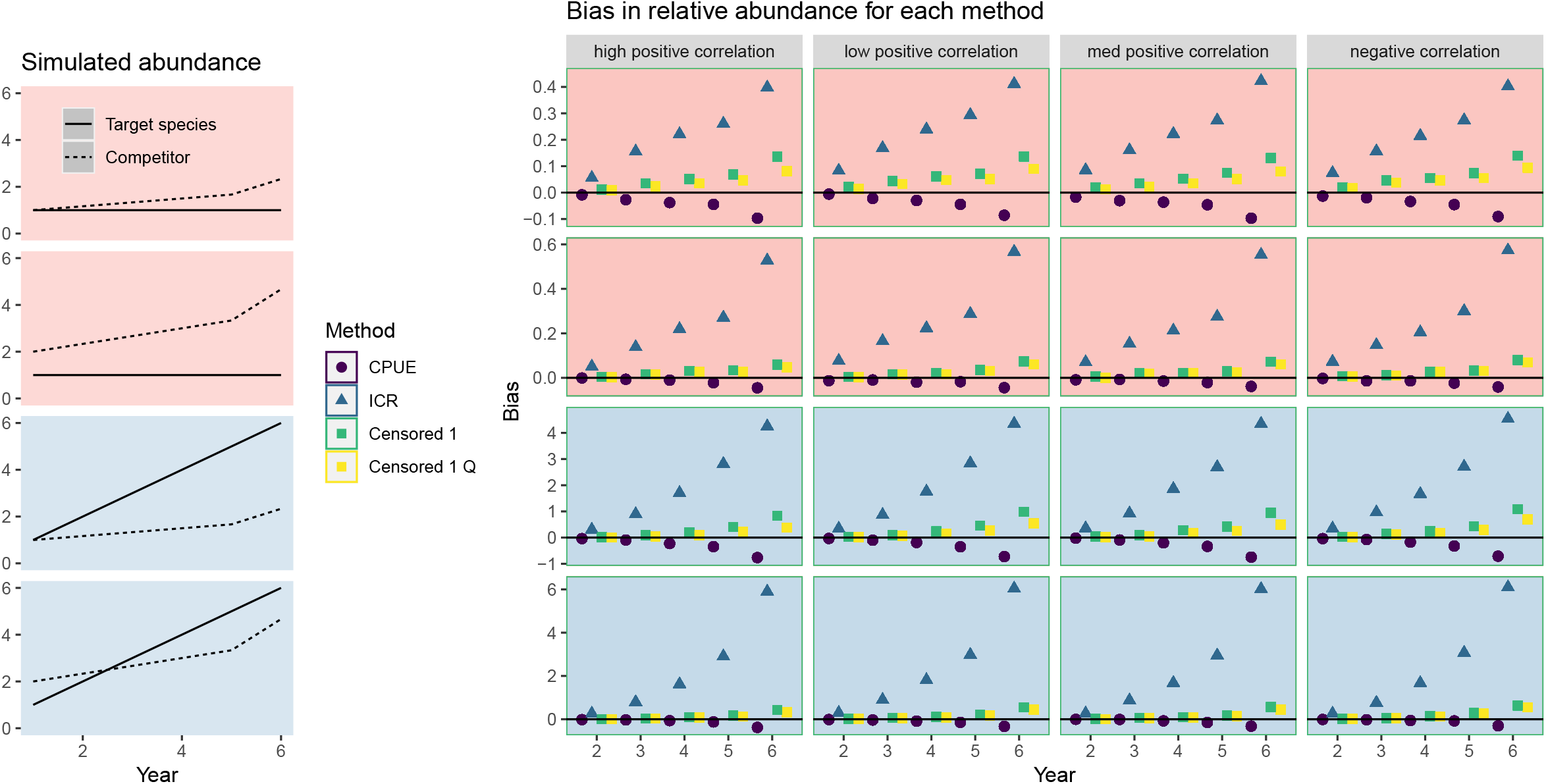
Bias results for experiment 5. Rows, columns, and methods as in Figure 3, with the extra methods ‘Censored 1’ and ‘Censored 1 Q’ which assume 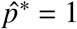. ‘Censored 1 Q’ considers all observed counts of the target species ≥ the 85^*th*^ observed annual quantile as ‘high-quality’, regardless of *p_tk_*. The bias from the censored method remains low in all settings and is at least as good as the CPUE method method, which is expected to perform well here due to the insensitivity of the target species to hook competition. The approximate 95% intervals are omitted since they are typically smaller than the plotting character.

**Figure S.11:**
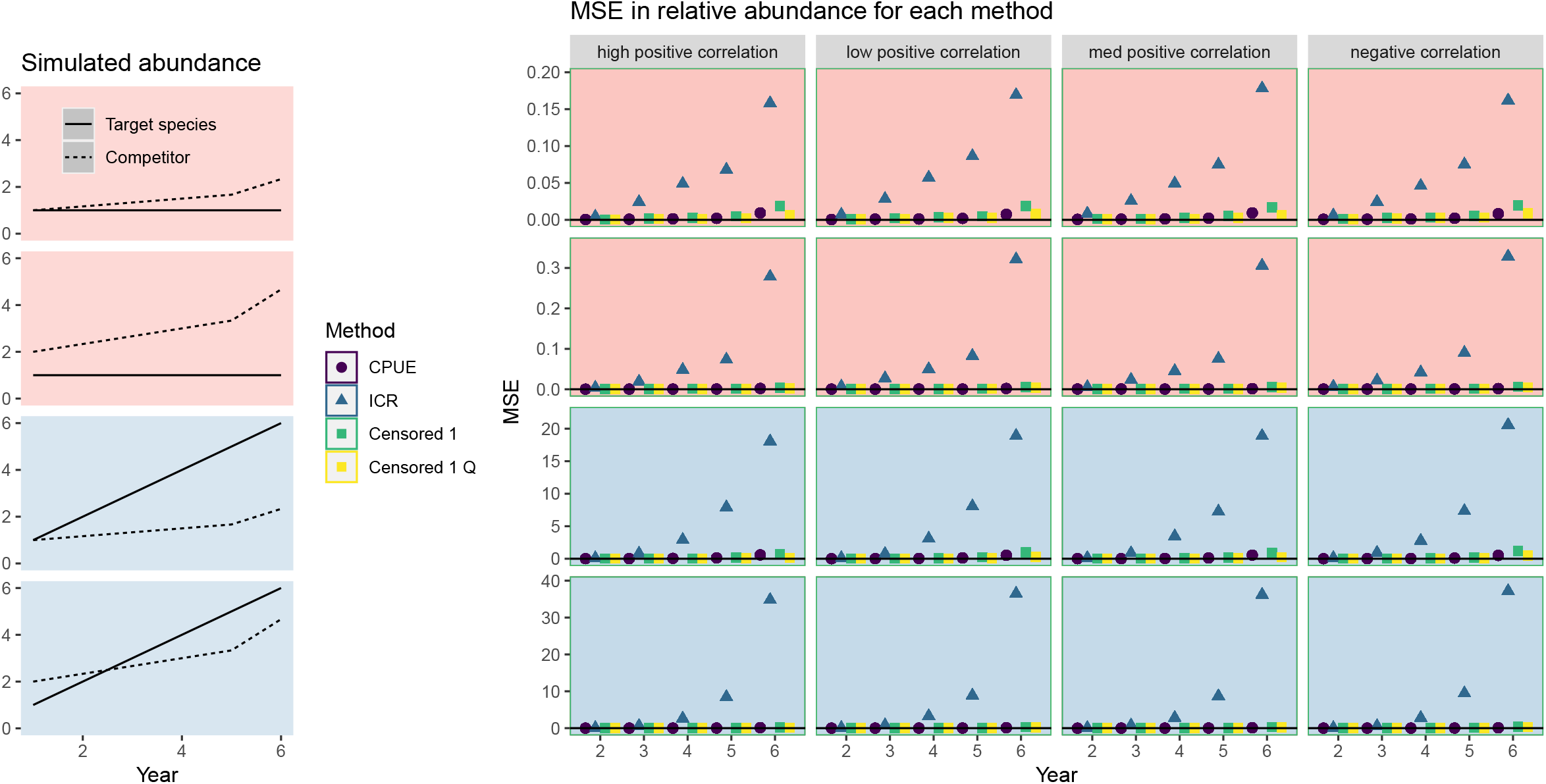
Median-squared error results for experiment 5. Details as for Figure S.10. Values for the censored method remain competitively low in all settings (compared with the CPUE method), but are highest in the negative correlation setting. The errors from the ICR method are extremely high. The approximate 95% intervals are omitted since they are typically smaller than the plotting character.

**Figure S.12:**
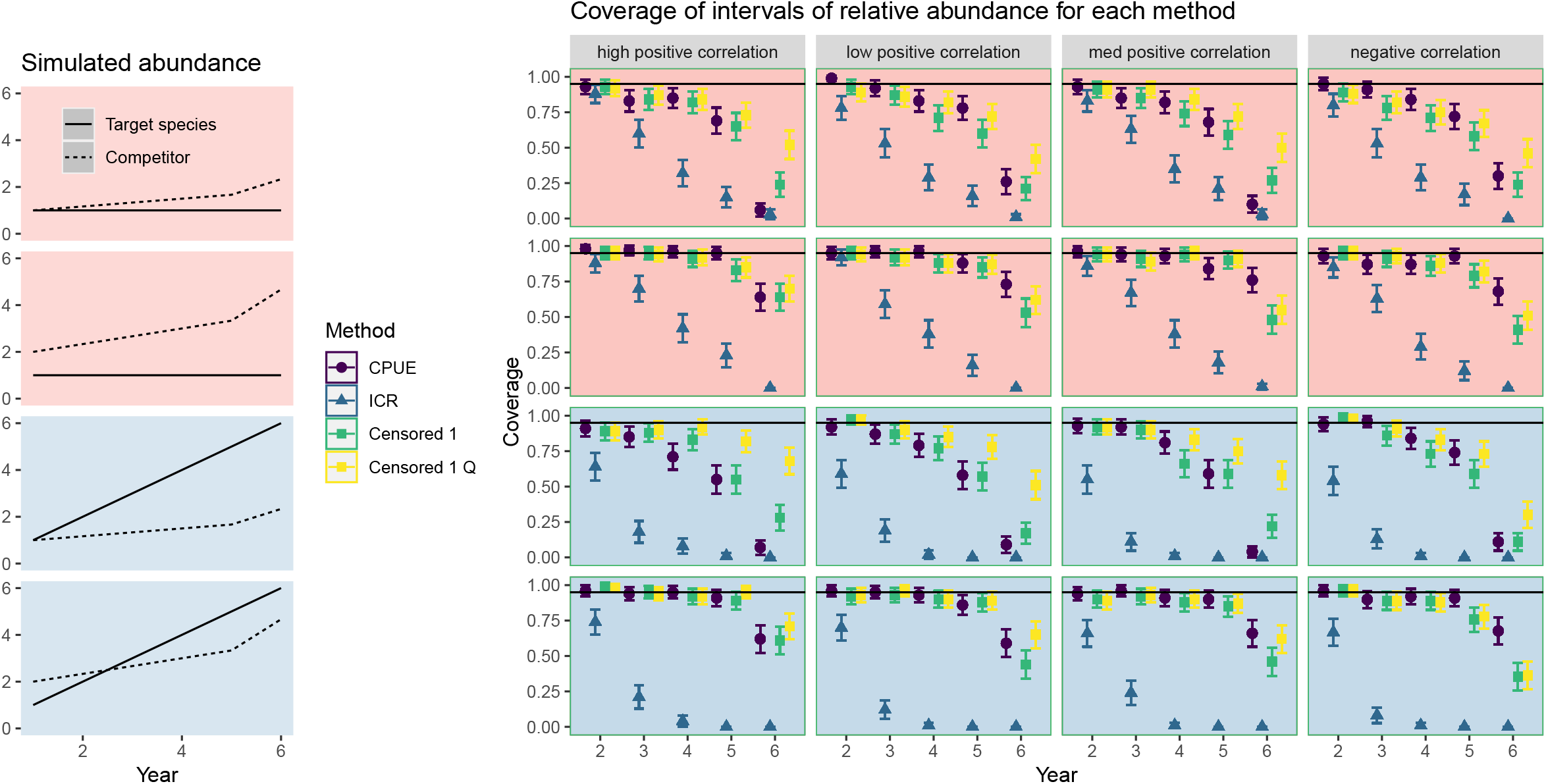
Coverage results for experiment 5. Details as for Figure S.10. The coverage levels for the censored method are higher than the competing methods in most settings, but can fall as low as ~20%. In general, the censored method’s coverage is lowest in the negative correlation setting, and highest when *Q* = 0.85 is specified.

### S.2 Proof of third theoretical example

We prove the result of the third theoretical example presented in the introduction of the main text. This considered two species over two years. The abundance of species A (denoted Λ_*At*_) was assumed stationary (Λ_*A*,1_ = Λ_*A*,2_), but species B increased in abundance in year 2 (Λ_*B*,1_ < Λ_*B*,2_). Species A was assumed to always reach the baits ahead of species B. There were 1000 baited hooks deployed in a single fishing event each year and 990 fish of species A were caught in both years. For species B, 0 and 9 fish were caught in years 1 and 2 respectively. We drop the *k* subscript due to the single fishing event each year.

Let *c_At_* denote the observed catch of species *A* in year *t* and let *F_t_* be the multiplicative competition adjustment factor from (S.4) but without depending on *k*, such that

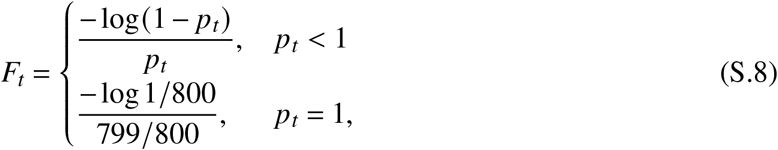

where *p_t_* is the proportion of the baits removed during the single fishing event of year *t.*

To estimate the relative abundance of species *i* in year *t*, the ICR method scales the catch per unit effort *c_At_*/1000 by *F_t_* as:

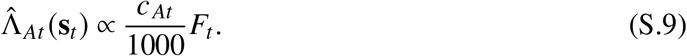

Knowing that species A can always reach baits ahead of species B suggests that estimates of relative abundance for species A should not change between the years. However, the multiplicative competition adjustment factor depends on the observed catch count of species B (through the dependency between *c_Bt_* and *F_t_*).

Substituting in the values 990/1000 and 999/1000 for *p_t_*, we see that the multiplicative scale factor *F_t_* approximately equals 4.65 and 6.91 in years 1 and 2. Since *c_At_* is fixed across the years, an increase of 6.91/4.65 ≈ 49% in the relative abundance of species A is estimated, yet its true abundance did not change.

Thus, the ICR method’s estimator of relative abundance is highly sensitive to changes in the abundances of competing species, which can easily occur in real-world settings.

### S.3 Deriving an upper bound to improve censored method’s convergence in data-poor settings

For some of the simulation experiments with 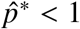, we placed an upper bound on the catch counts of the target species, denoted *u_itk_*, to improve the convergence of the censored method. We now derive the upper bound, argue why it is likely conservative when used in general settings, and then empirically confirm this claim for the simulated data of experiments 2-5. Note that subsequent improvements in the optimization routine have largely eliminated the need to implement this upper bound (the case study did not implement an upper bound for any species).

Let a collection of longline fishing events in year *t* be indexed by *k*, each with *h_tk_* baited hooks deployed and with assumed censorship threshold values 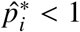 for each species. First, assume that attracted individuals of species *i* do not have any difficulties locating a baited hook when a proportion less than 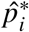 of them have had their baits removed. We refer to this period of the soak time as stage 1. We do not need to make any further assumptions about the arrival rates of the fish to the baits during this stage. We denote the total catch counts of species *i* after soak time stage 1 as 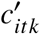.

Next, at a given moment in time, 0 < *τ* < *S_tk_*, during the soak time *S_tk_*, denote the current proportion of baits removed as *p*(*τ*). We refer to the period of the soak time where 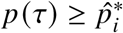, when it occurs, as stage 2. During stage 2, we assume that each species arrives at the fishing gear at times which are independent exponentially distributed with constant instantaneous rates *ρ_itk_*. Then, the probability of capture is assumed to decrease linearly with time as

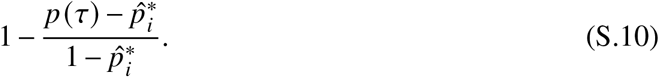

The assumed exponentially-distributed bait removal times during the stage 2 are chosen so that the fishing process in stage 2 satisfies the assumptions required of the ICR scale factor adjustment *F_tk_*.

Let 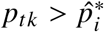. If we knew the number of fish of the target species that were caught during stage 2, denoted 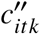, then an ICR-based estimate for the number of fish of species *i* that could have been caught if more hooks been available during the stage 2 soak time would be:

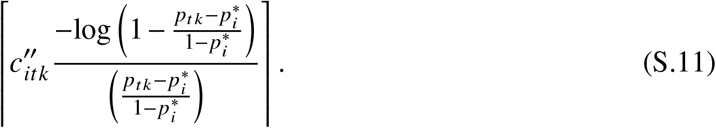

Thus, under the above assumptions, the ICR-based estimate for the total number of fish caught during the entire soak time would equal:

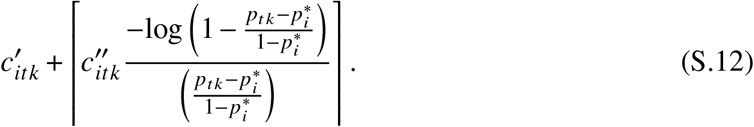

Clearly, without hook timers, we will be unable to determine 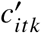 or 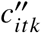. However, since we are only requiring *u_itk_* to be an upper bound, we replace these numbers with known quantities: i) *c_itk_*, the observed catch count of species *i* after the soak time *S_tk_*; ii) the number of hooks observed with their bait removed by *all sources* in the second stage, 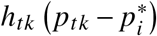. Thus, we define a conservative upper bound for each species *i* during fishing events where *c_.tk_* (the observed number of bait removals by all causes during fishing event *k* of year *t*) satisfies 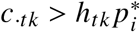:

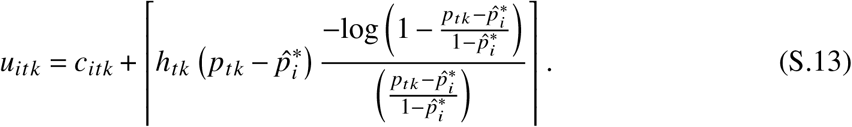

In words, our upper bound for the censored observations is the observed species-specific catch count *c_itk_* plus the observed number of baited hooks removed in stage 2 by all sources, 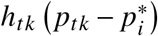, scaled by the competition adjustment factor *F_tk_* evaluated only for the stage 2 hooks. This value is then rounded up to the nearest integer.

Note that we are not assuming that the above model describes the true bait removal process in stage 2. Instead, we are trying to utilize a set of assumptions that justify a reasonable upper bound to improve the convergence of the censored method. Given that we are assuming the instantaneous catch rates of each species is non-decreasing throughout the soak time of stage 2 and that all bait removals in stage 2 were due to species *i,* this should ensure that *u_itk_* is a conservative upper bound in many settings where multiple species are competing for large numbers of hooks.

We empirically evaluate this claim. The upper bound is always used in simulation experiments 2-4. For these settings, the upper bound lies above the true number of attracted individuals *C_itk_* in 99.9% of cases and exceeds it by a mean number of 350 fish when 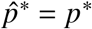. Thus, *u_itk_* is a a successful (and highly conservative) upper bound which should have only a minor impact on the results of the simulation study, compared with if the simulation had been run without any upper-bound at all. Thus, we are confident that the results from experiments 2-4 with the upper bound used generalize to the setting where no upper bound is used, without us needing to re-run the computationally-costly simulation experiments without the upper-bound and using the new optimization routine.

An upper bound was considered in experiment 5 when 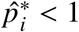. However, in this experiment, results showed that performance was optimized with 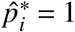 when no upper bound was used and so we do not repeat the empirical evaluation for experiment 5. The upper bound was not used in experiment 1.

### S.4 Exploratory analysis for the Case Study

#### S.4.1 Assessing the linear relationship between effective skate and catch counts

The geographic extent of the IPHC data used in the Case Study is shown in Figure S.13, along with the locations of the stations on the 10 × 10 nautical mile square grid.

We investigate the hypothesis that the measure of effective skate (Yamanaka *et al.*, 2008) is a suitable proxy for the integral of the response surface (equation 11) for all 11 species analyzed in the case study. Despite the IPHC data being a survey with fixed protocols deployed, in some years, only the first 20 hooks of each skate were enumerated. The effective skate values have been scaled to account for this, but we remain unsure about the validity of this correction. For effective skate to be a valid proxy implies that the expected catch count of a species scales linearly with effective skate, holding all other factors constant. We test this now.

**Figure S.13:**
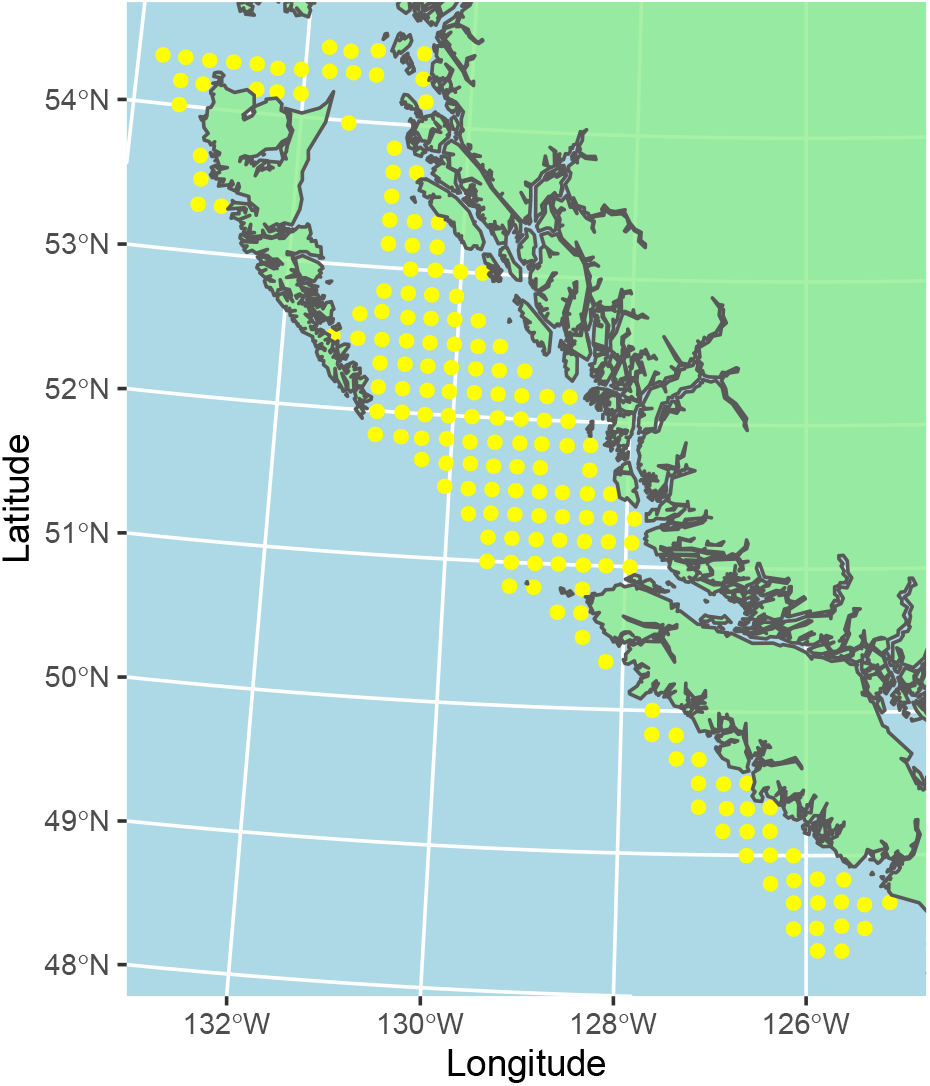
The locations of the 171 International Pacific Halibut Commission stations that were fished in Canadian waters off British Columbia between 1998-2020.

Let *i* denote the species and let *E_tk_* denote the effective skate value corresponding to the fishing gear deployed in year *t* at station *k*. We fit the following negative binomial GAMM to the observed species catch counts (*c_itk_*) and perform exploratory analysis to empirically assess this assumption:

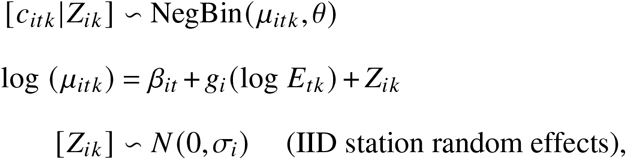

where *g_i_* (log *E_tk_*) denotes (possibly nonlinear) effects of effective skate, to be estimated with cubic spline smooths, with the degree of ‘wiggliness’ chosen automatically. We use the mgcv package to perform the model fitting (Wood, 2004, 2011; Wood *et al.*, 2016).

Thus, the above model attempts to control for the changes in abundance through time and the station-station variability with random effects. Then, lastly, any nonlinear effects of effective skate are estimated in the *g_i_*(·) terms. Note that no adjustments are made for hook competition and hence the model is only exploratory in nature. The negative binomial model is chosen to account for overdispersion in the catch counts, whilst being available in the computationally-fast mgcv R package, unlike the Poisson lognormal used elsewhere in this work.

We plot the mean and 95% confidence intervals for the *g_i_*(·) terms for all 11 species. Figure S.14 fails to show any strong departures from a linear trend between the mean catch counts and the value of effective skate for 6 of the species, with the 95% confidence intervals for these species covering a line with unit slope. However, significant departures are detected for 5 species, most severely for Arrowtooth Flounder, Pacific Cod, and Sablefish. Overall, we conclude that the linearity assumption between effective skate and the mean catch counts appears to be a reasonable approach to account for heterogeneous fishing effort across the fishing events for 6 of the 11 species. Future work should investigate how to better control for heterogeneous observer effort in the other 5 species. We continue to use effective skate as an offset value to account for heterogeneous fishing effort when estimating the relative abundances of all 11 species, however, the estimated indices should be treated with caution.

**Figure S.14:**
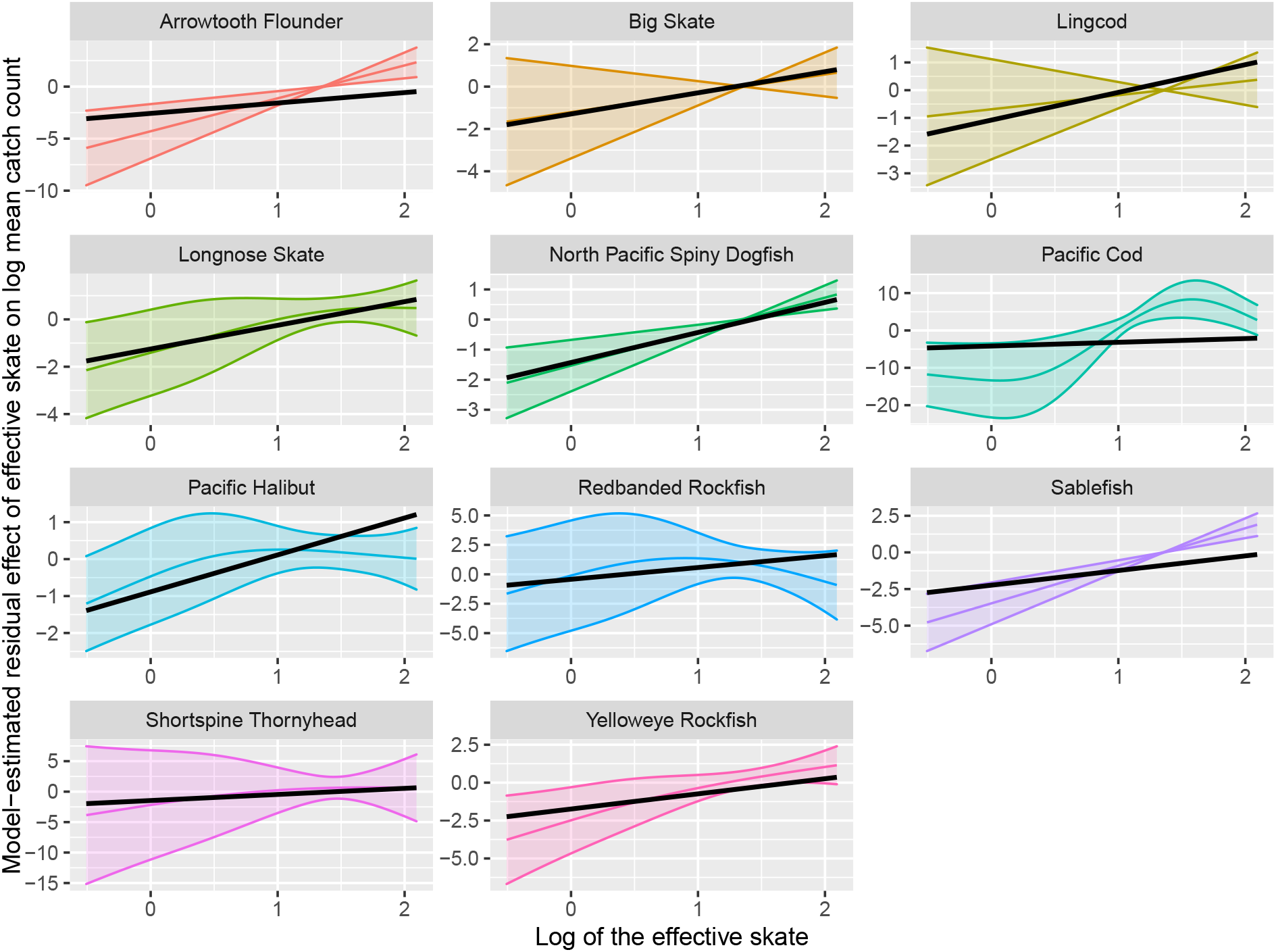
A plot showing the estimated nonlinear effects of log effective skate on the log mean catch counts of all 11 species, after controlling for temporal effects. 6 out of 11 of the 95% confidence intervals cover the unit slope line across all values of log effective skate. This indicates that the effective skate is a useful proxy for the heterogeneous fishing effort experienced across the fishing events for these species. Future work should investigate how to control for heterogeneous observer effort in the other 5 species.

#### S.4.2 Empirically assessing values of 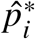

Next, we investigate suitable choices for the censorship proportion, 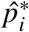, for all 11 species. To do this, we once again attempt to control for the temporal effects on the mean abundances of both species. Based on the results of the previous subsection, we remove the cubic spline smoother of effective skate and assume a linear relationship.

To detect this hook saturation effect, we include a cubic spline of the proportion of baits removed during each fishing event (*p_tk_*) inside the model as follows:

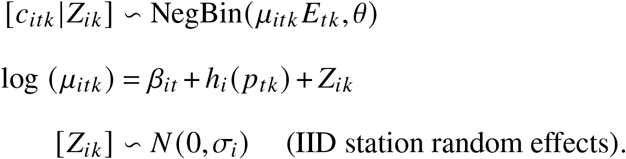

Thus, the above model once again attempts to control for the changes in abundance through time and the station-station variability with random effects. Then, lastly, any residual nonlinear effects of hook competition should appear in the *h_i_*(·) terms.

We plot the mean and 95% confidence intervals for exponentially transformed *h_i_*(·) terms for all 11 species. Figure 6 shows that all species experience a statistically-significant decline in the mean catch count exceeding 10%, with the exceptions of: i) dogfish which experiences an statistically significant increase in mean catch count exceeding 65%; ii) Shortspine Thornyhead which experiences a non-significant decline of only 1%. The plots show that statistically significant declines in mean catch counts are first seen around *p_tk_* = 0.85 for Lingcod and Redbanded Rockfish, and around *p_tk_* = 0.95 for the remaining 7 species.

Based on advice given in the main text from simulation experiment 6, values of 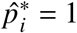 for dogfish and Shortspine Rockfish, 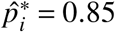 for Lingcod and Redbanded Rockfish, and 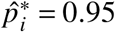 for all other species should be used. Furthermore, based on the decision rule given in the main text, we should treat the censored method’s indices with caution for Shortspine Thornyhead until we can determine that neither fish is ‘immune’ to hook competition from other species.

### S.5 Additional details on the case study models

We assume that doubling the value of effective skate of a fishing event doubles the expected integral of the response function attributable to the fishing event. By denoting 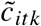 to be the observed raw catch count for the CPUE and censored method, and the hook-competition adjusted value for the ICR-adjusted method (i.e. equal to *c_itk_ F_tk_*), we can formally write all three methods under the same framework as:

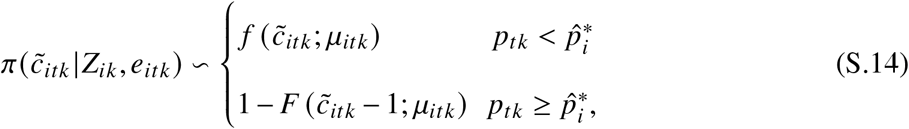

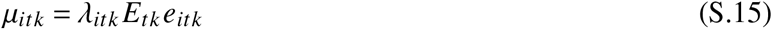

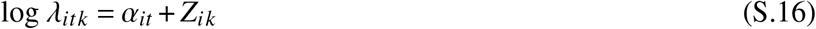

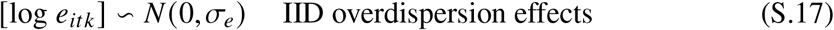

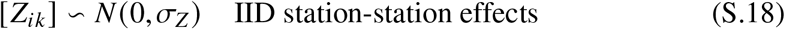

where *E_tk_* is the effective skate number of fishing event *k* in year *t* and *σ_e_* and *σ_z_* are standard deviations to be estimated. By setting 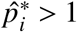 for the CPUE and ICR-adjusted methods (which do not require values of 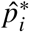), (S.14) reduces to just its first row, correctly specifying those models. The temporal trends are captured by the unique intercept each year, *α_it_*. These are unstructured fixed effects to allow for flexible changes in abundance to be inferred and avoid smoothing over the temporal variation. Normalized estimates (each series divided by its geometric mean) of exp (*α_it_*) are used to represent the relative abundance trends in Figure 7 for three species, and in Figures S.17-S.19 for all species, characterised by their 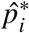 values.

All models were fit using the sdmTMB and TMB software packages (Anderson *et al.*, 2022; Kristensen *et al.*, 2016) within R (R Core Team, 2022). Models converged for 10 out of the 11 species without requiring any upper bounds *u_itk_* be placed on the catch counts or any other modification. To attain convergence for Redbanded Rockfish, the upper bound derived earlier in the Supplementary Material was used and the upper 5% of catch counts each year were considered high quality, regardless of the values of *p_tk_*.

### S.6 Additional case study results

Figures 8, S.15, and S.16 help us to understand the magnitudes of the differences between Figures S.17, S.18, and S.19. They plot the estimated percentage change in relative abundance estimate from the censored method compared to the CPUE method for each year, plotted with the proportion of fishing events with 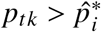 on the x-axis. In addition to assessing the absolute magnitudes of change, the differing sensitivities of each species to hook competition can also be inferred by assessing the relative magnitude of change witnessed for each species. Of note are Pacific Halibut and dogfish which are both estimated to be only slightly impacted by hook competition. Conversely, Lingcod, Sablefish, and Pacific Cod are all estimated to be acutely sensitive to hook competition by the censored method.

Figure S.20 shows that the rank correlations between the CPUE and ICR methods’ indices are consistently high (≈ 0.93) when averaged across all 11 species, whereas the rank correlations between the censored method’s indices and the CPUE and ICR methods’ indices are much lower (≈ 0.70 and ≈ 0.83 respectively). Similar results are seen with respect to the percentage overlap of 95% confidence intervals for abundance (Figure S.20). All of the intervals overlap for the CPUE and ICR methods, unlike the censored method’s intervals which overlap with around 80-85% of the CPUE and ICR methods’ intervals.

Figure S.20 also shows that the estimates of uncertainty are smallest for the CPUE method and largest for the ICR-based method, with the censored method in-between. Furthermore, the MAD and ‘Percentage Unchanged’ results of Figure S.20 show that relative abundance estimates from the ICR method are most variable and those from the CPUE method are least variable, while the ‘Range’ results indicate that the ICR method infers the greatest fluctuations in abundance.

**Figure S.15:**
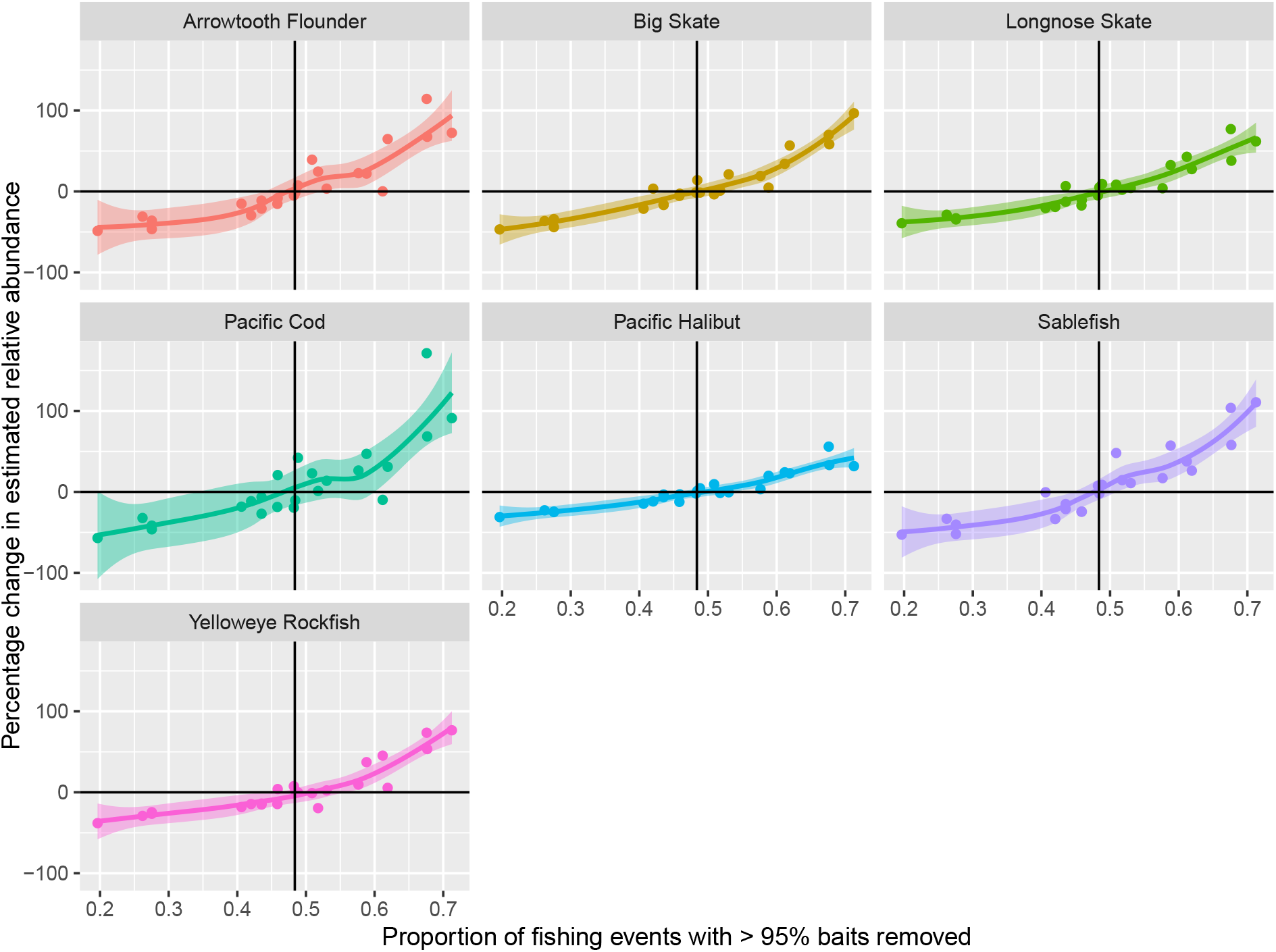
As for Figure 8 but for the seven species with 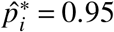, with vertical lines showing the average annual proportion of fishing events experiencing at least 95% bait removal. The general conclusions are the same as for Figure 8. Here, the censored method infers that Pacific Halibut is the least affected by hook competition, with Pacific Cod the most affected.

**Figure S.16:**
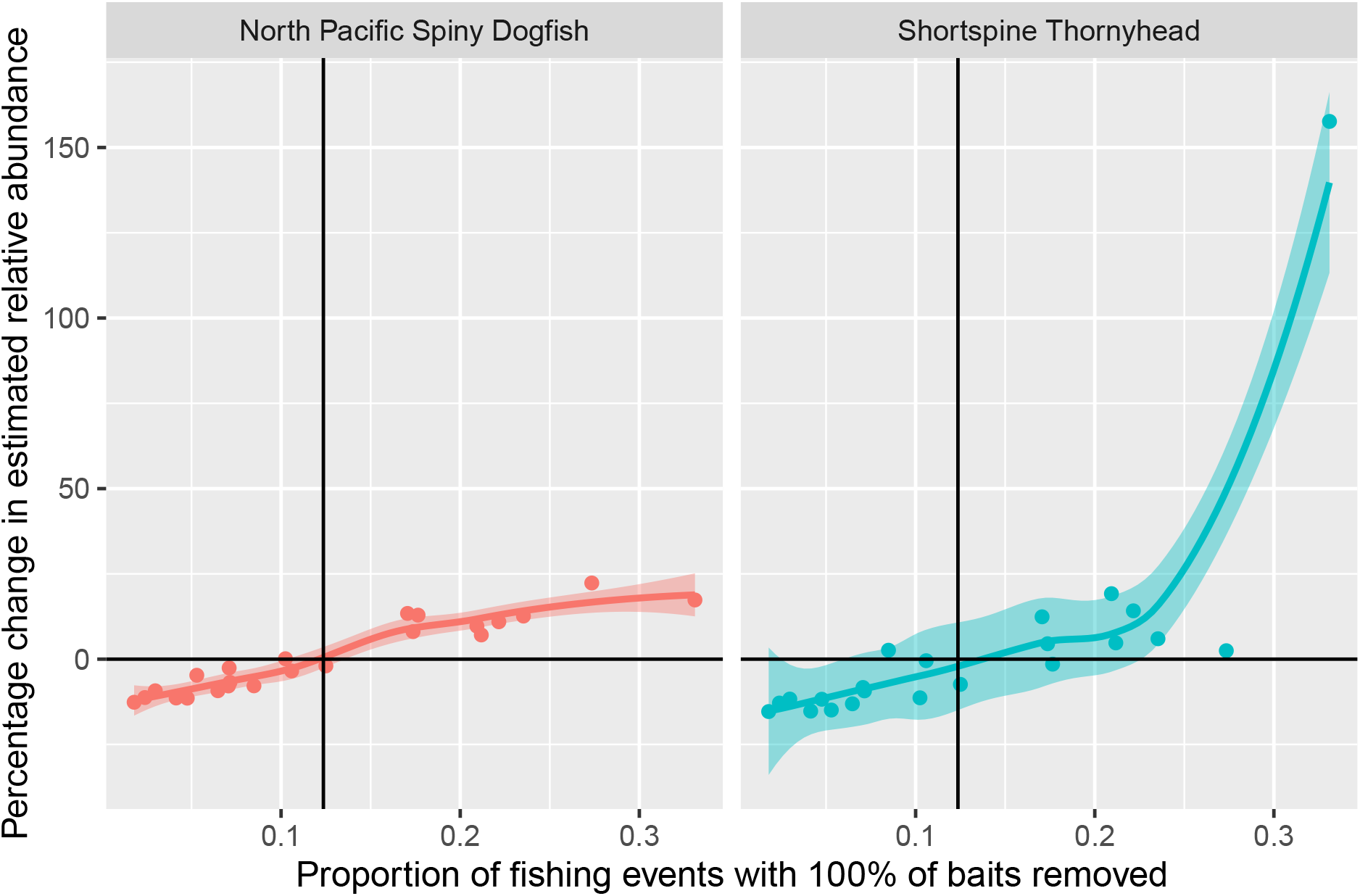
As for Figures 8 and S.15 but for the two species with 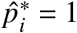, with vertical lines showing the average annual proportion of fishing events experiencing 100% bait removal. Again, the general conclusions are the same as for Figure 8. Here, the censored method infers Shortspine Thornyhead to be more severely affected by hook competition than dogfish, although this conclusion may be unduly influenced by a single outlier seen in the top right.

**Figure S.17:**
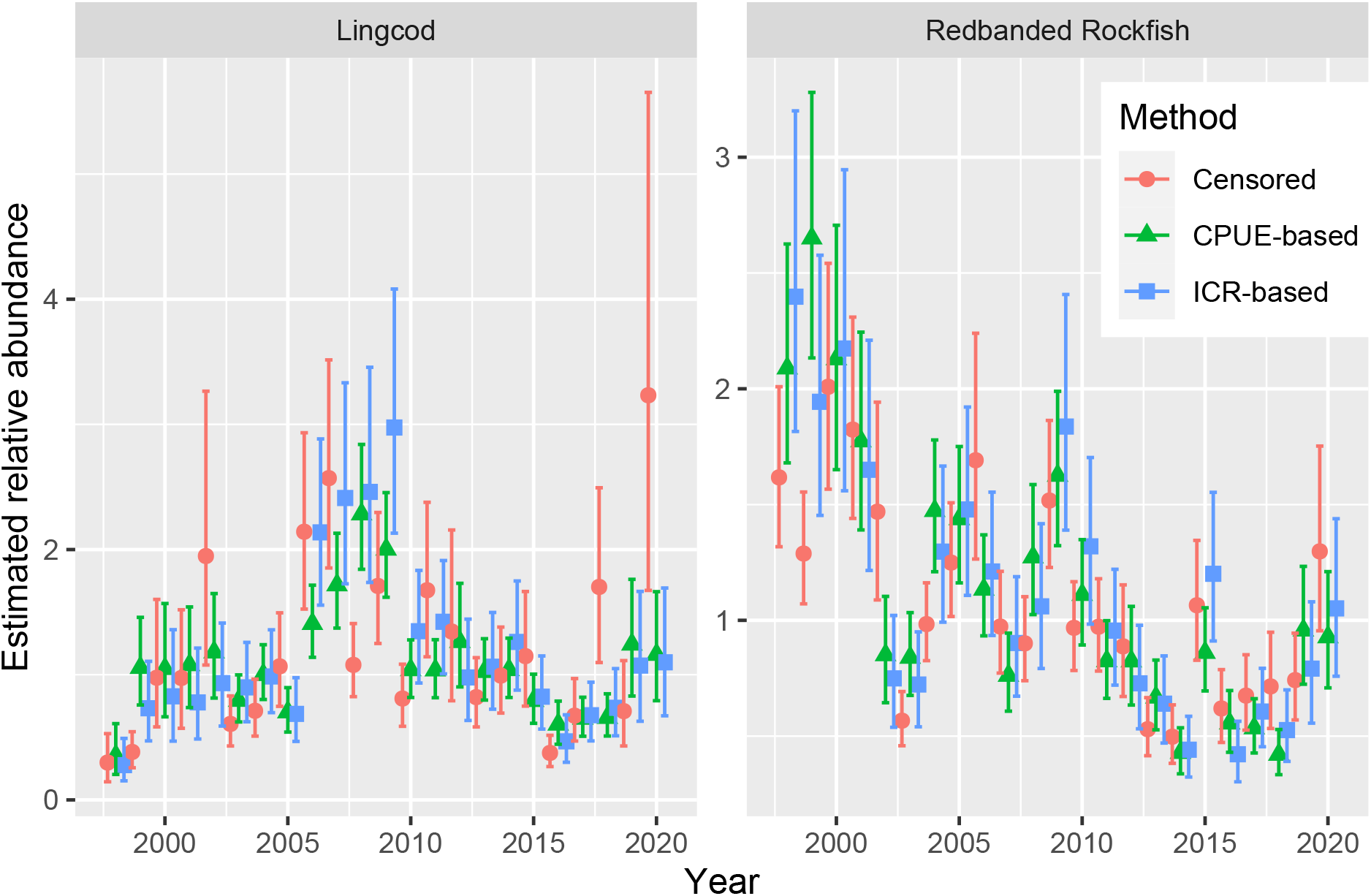
Plots showing the estimated relative abundances of the two species for which 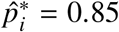 was judged appropriate. Shown are the mean and 95% confidence intervals from three model-based methods. The points and intervals have been ‘jittered’ along the x-axis for readability. The ICR and CPUE methods closely agree for both species. Conversely the censored method’s estimates differ greatly for Lingcod.

**Figure S.18:**
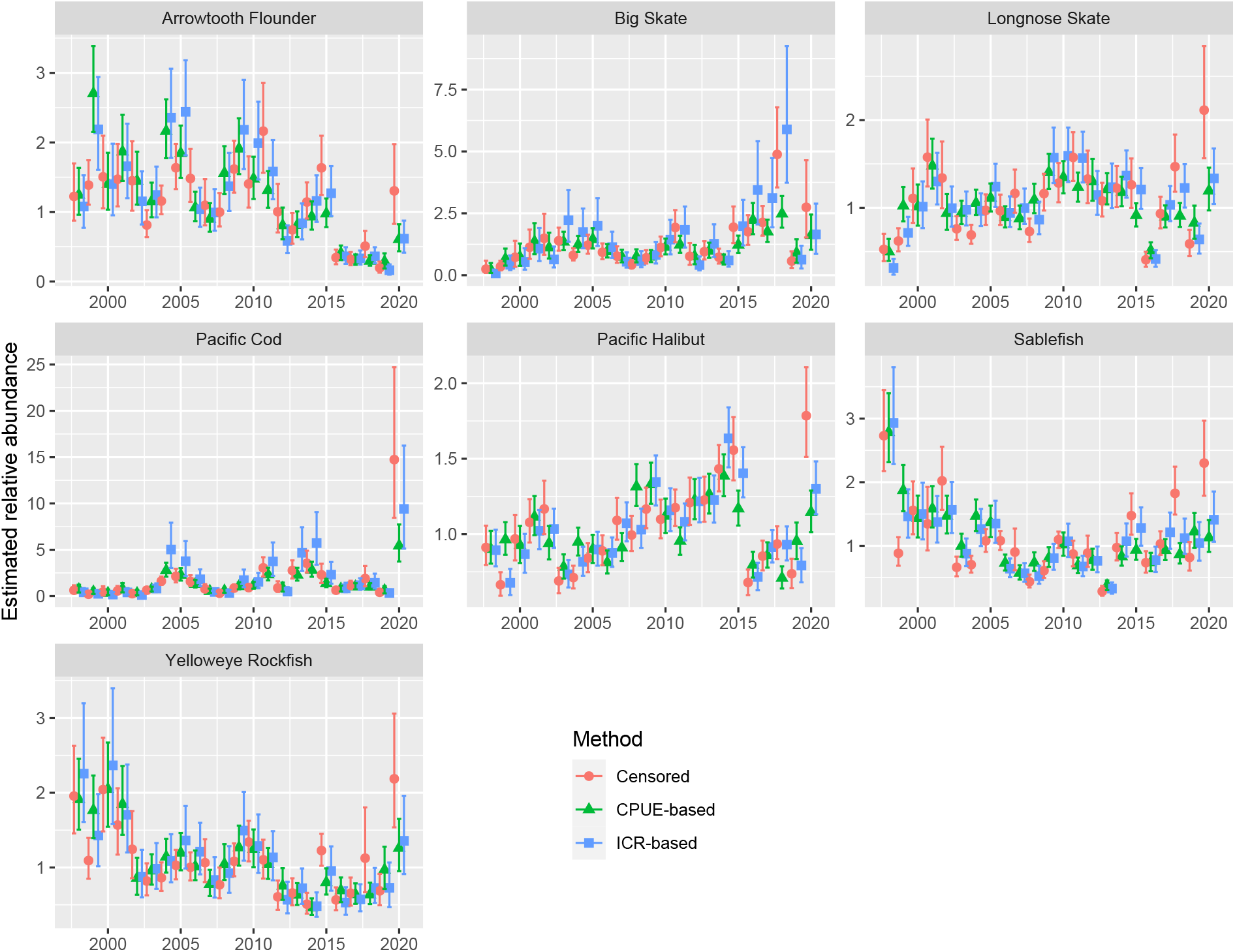
Plots showing the estimated relative abundances of the seven species for which 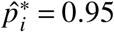 was judged appropriate. Shown are the mean and 95% confidence intervals from three model-based methods. The points and intervals have been ‘jittered’ along the x-axis for readability. The ICR and CPUE methods closely agree for all species. Conversely the censored method’s estimates differ greatly, most notably for Sablefish and Arrowtooth Flounder.

**Figure S.19:**
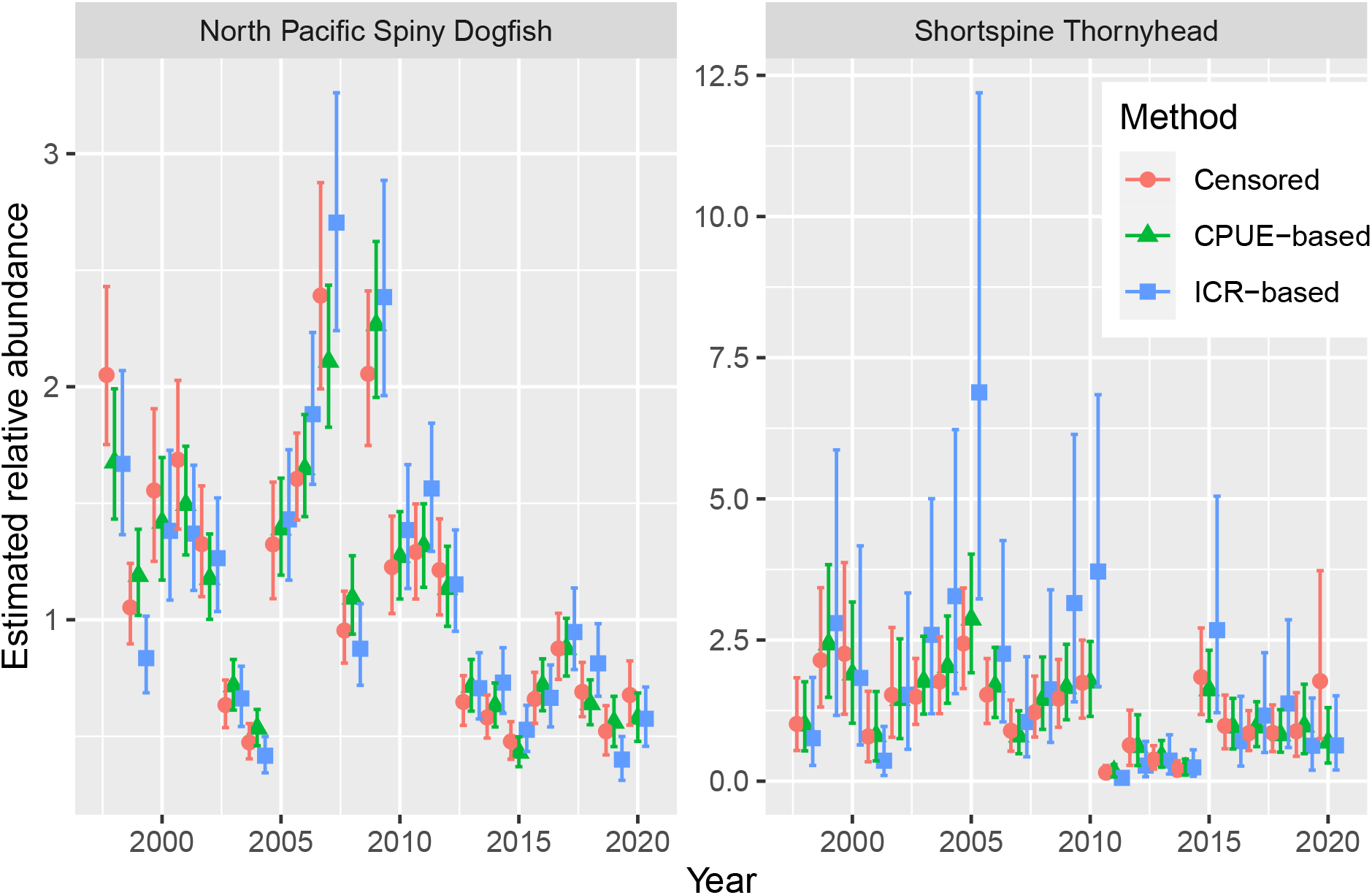
Plots showing the estimated relative abundances of the two species for which 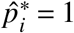 was judged appropriate. Shown are the mean and 95% confidence intervals from three modelbased methods. The points and intervals have been ‘jittered’ along the x-axis for readability. The CPUE and censored methods’ estimates closely agree for both species. Conversely the ICR method’s estimates differ greatly for Shortspine Thornyhead.

**Figure S.20:**
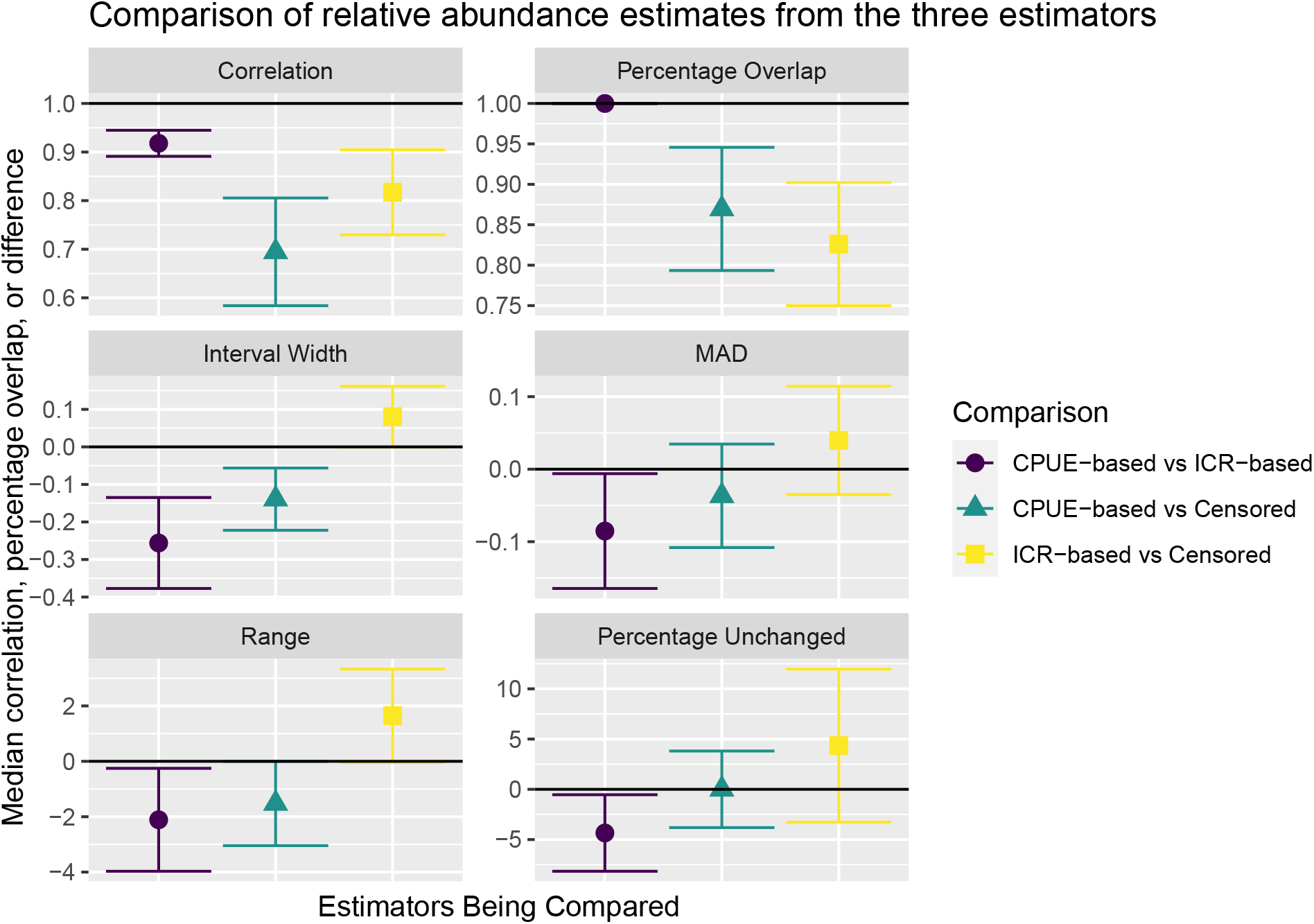
Plots showing empirical comparisons of the relative abundance estimates from the three model-based indices averaged across all 11 species. Confidence intervals are computed assuming the 11 species are a representative sample of all species. Shown are: Spearman’s rank correlations between the intervals, percentage overlap between the competing 95% confidence intervals, differences in the widths of the 95% confidence intervals, differences in the median absolute deviations about the median (MAD) values, differences in the range of abundance estimates, and differences in the percentage of 95% confidence intervals covering 1 (i.e. abundance is not significantly different from the mean abundance). The censored method’s estimates are least correlated with and its intervals overlap least with those from the other two methods. The CPUE intervals are narrowest and the ICR intervals are widest. The CPUE estimates of relative abundance vary the least (smallest range and smallest MAD).

### S.7 R code for fitting the censored method

We provide R code to simulate censored longline count data of a single species *Y_i_* from the following Poisson regression model:

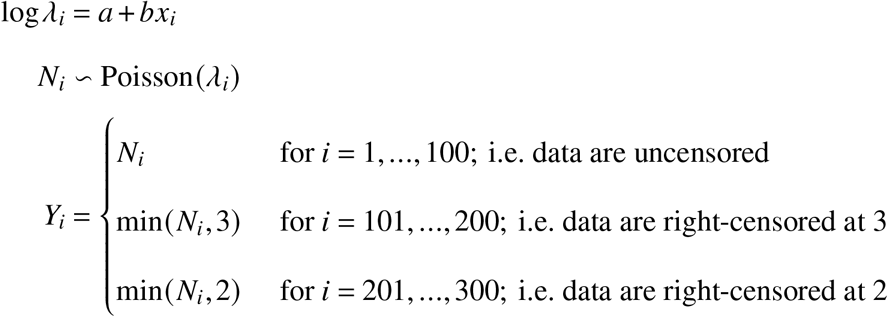

where *x_i_* are values of an observed covariate (for *i* = 1,…, 300), *a* and *b* are parameters relating log*Λ_i_* to *x_i_*, *Λ_i_* is the mean number of attracted fish, and the *Y_i_* are the observations. The first 100 observations are uncensored and collected by trawling (i.e. *Y_i_* = *N_i_* for *i* = 1,…, 100), the second 100 observations are right-censored at 3 (i.e. collected by longline fishing gear with only 3 hooks deployed), and the final 100 observations are right-censored at 2 (i.e. collected with longline fishing gear with only 2 hooks deployed). Note that *p*\* = 1 and identical integrated response functions across the three fishing gears are assumed for this simulated example.

The goal is then to account for the different fishing gear and hence levels of censorship to infer the parameters *a* and *b.* In R, we simulate the data as follows:

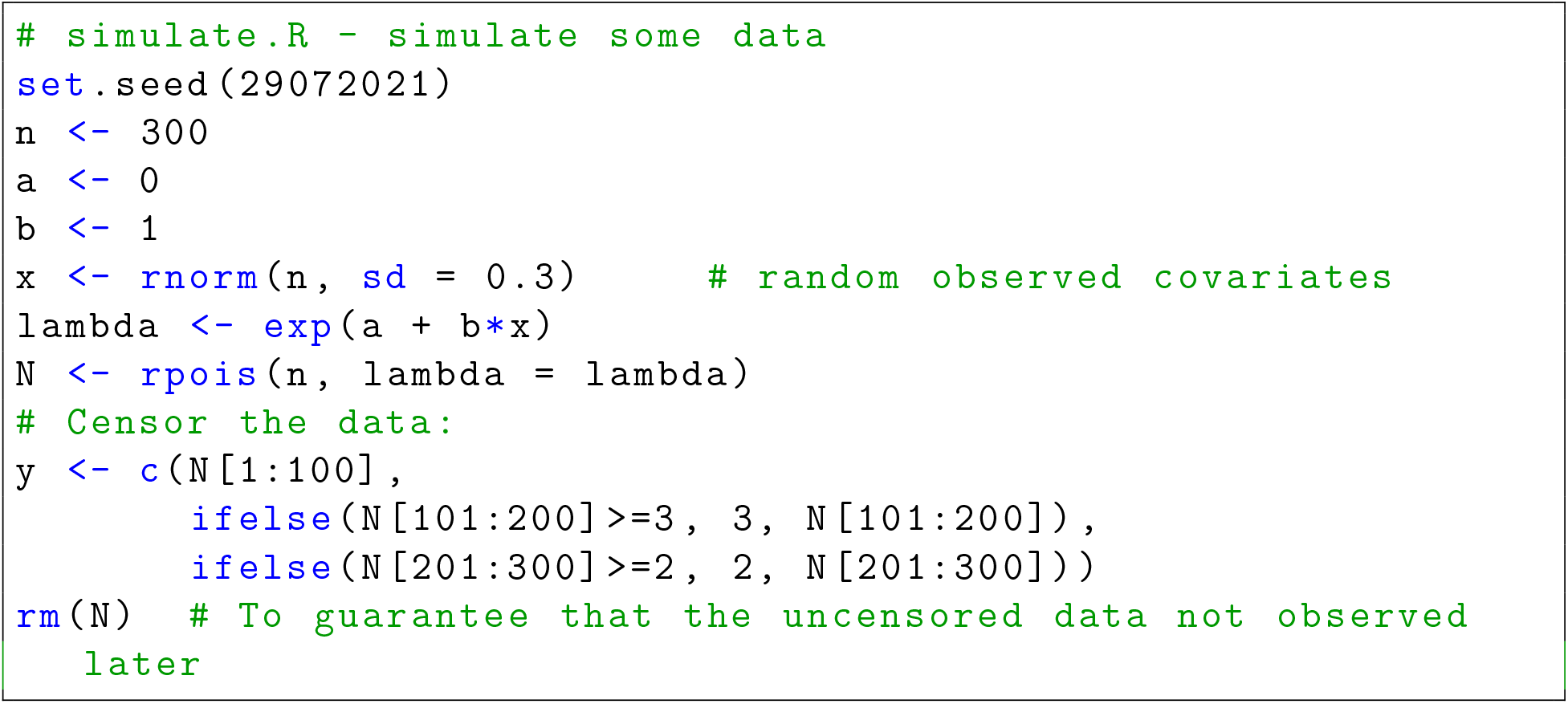

Next, we show how R-INLA, sdmTMB/TMB, and brms/stan can be used to correctly model the censored fishing data.

#### S.7.1 R-INLA code

To fit the censored model in R-INLA, we run the following R code:

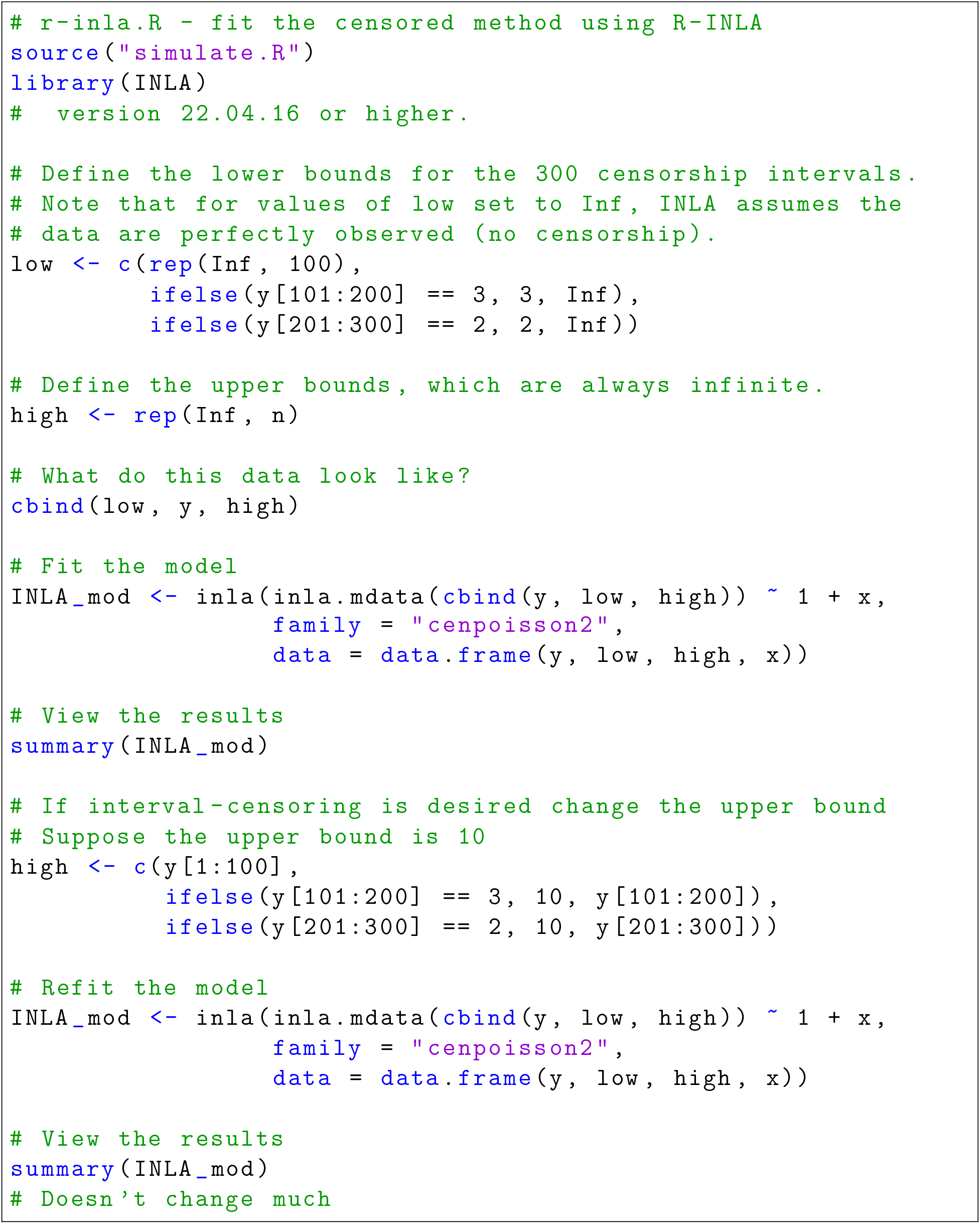

#### S.7.2 TMB code

To fit the censored model in sdmTMB and hence TMB, we run the following R code:

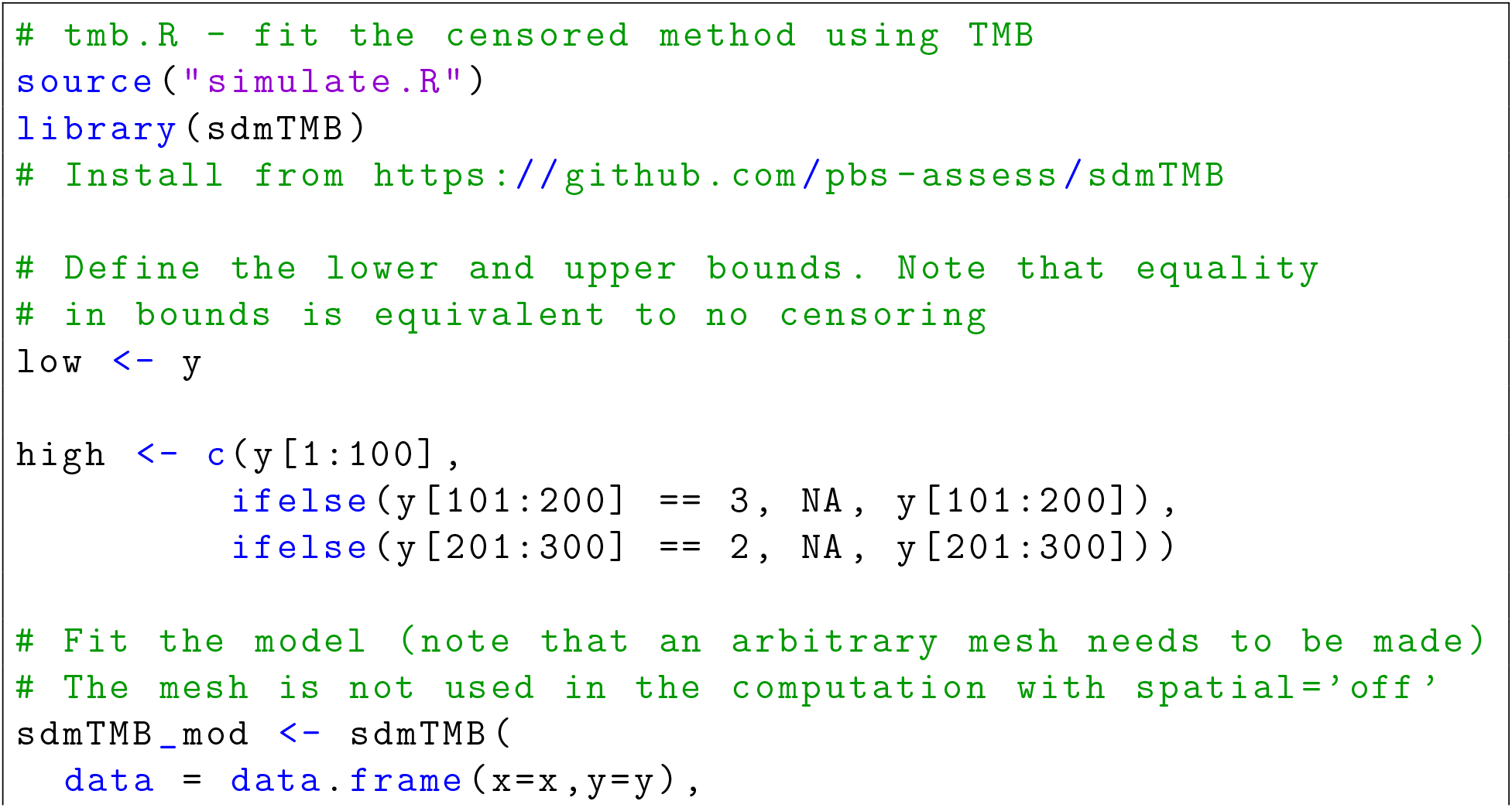
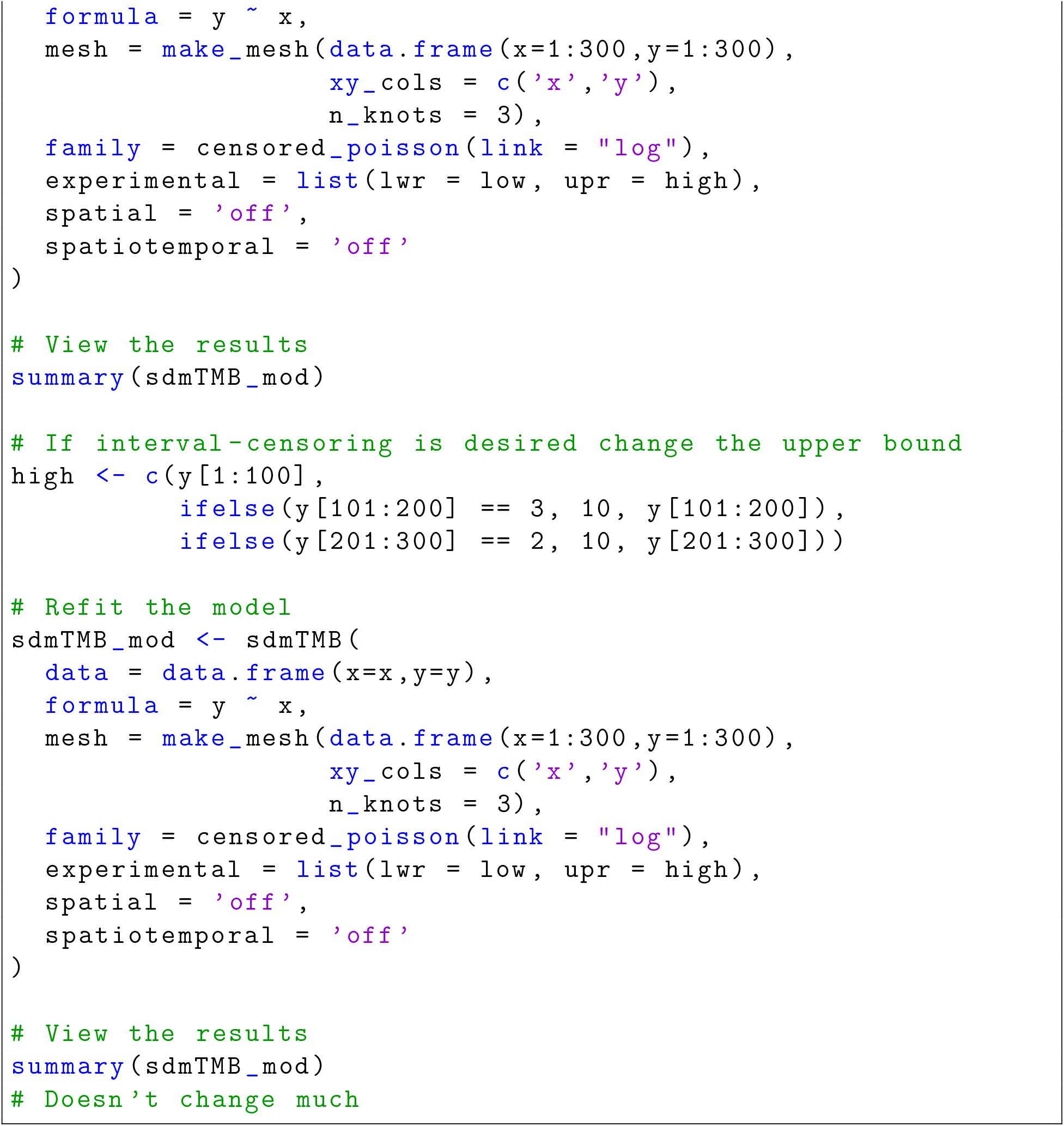

#### S.7.3 Stan code via the BRMS package

To fit the censored model in brms and hence stan, we run the following R code:

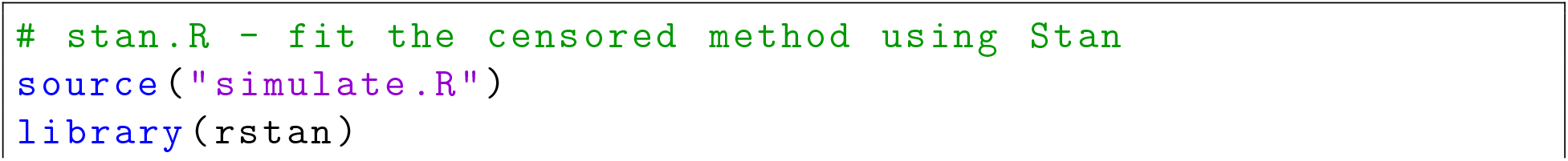
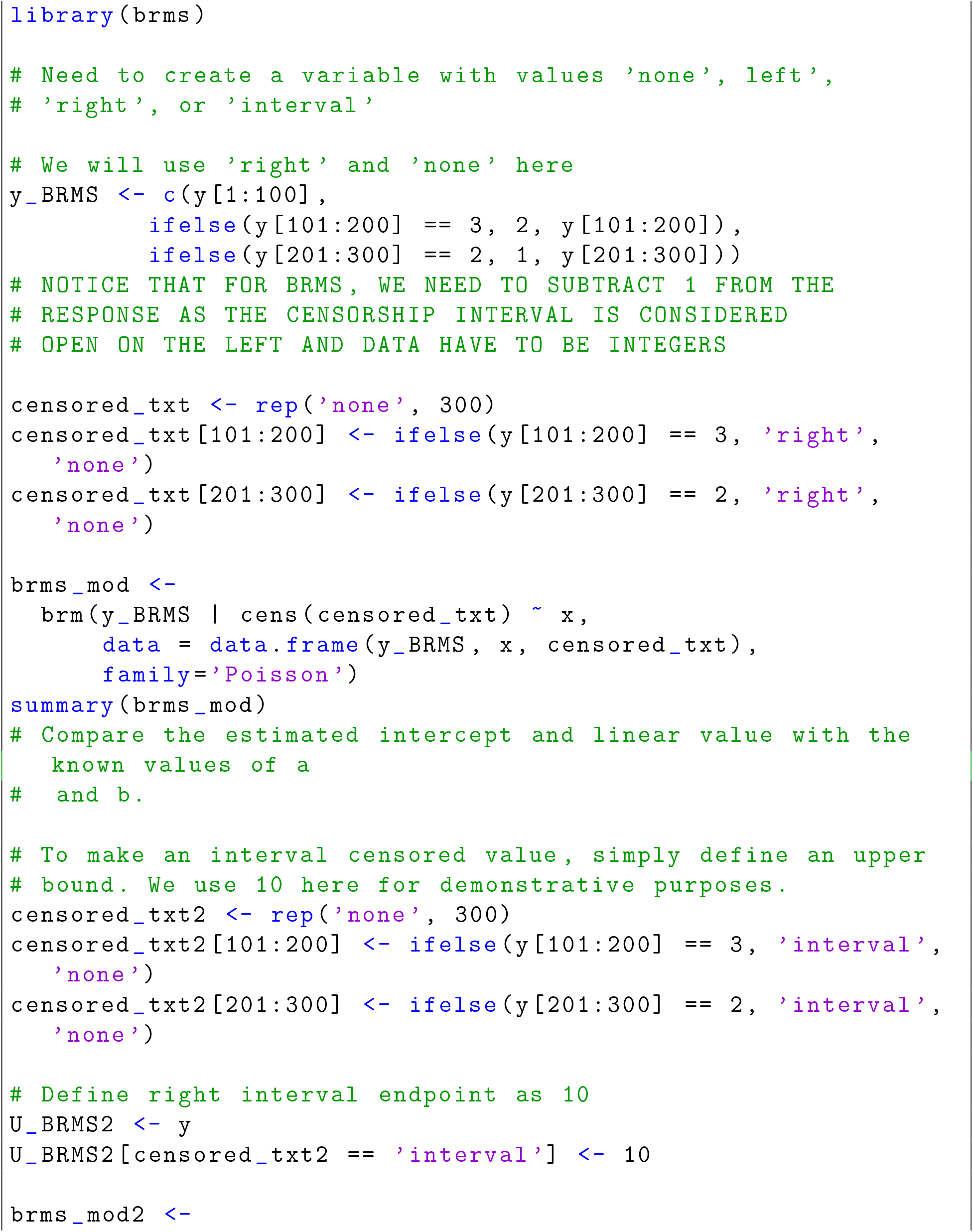
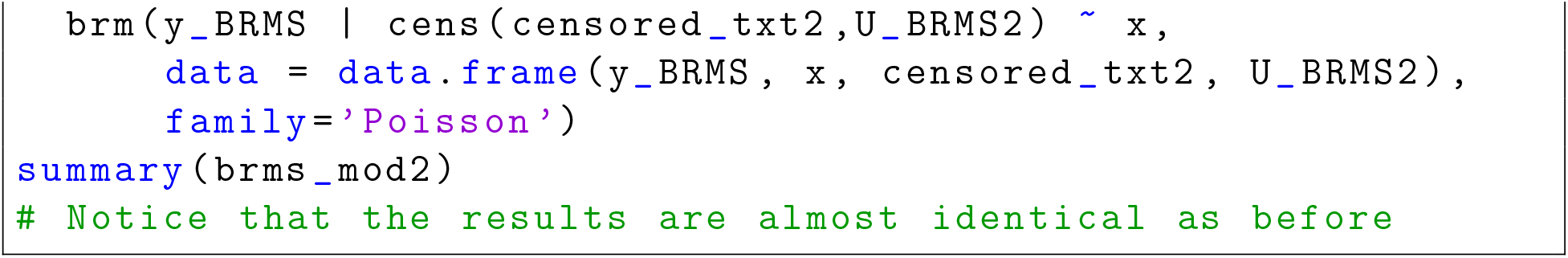

## References

Anderson, S. C., Keppel, E. A., and Edwards, A. M. (2019). A reproducible data synopsis for over 100 species of British Columbia groundfish. DFO Can. Sci. Advis. Sec. Res. Doc., 2019/041, vii + 321 p.

Anderson, S. C., Ward, E. J., English, P. A., and Barnett, L. A. K. (2022). sdmTMB: an R package for fast, flexible, and user-friendly generalized linear mixed effects models with spatial and spatiotemporal random fields. Available at https://github.com/pbs-assess/sdmTMB.

Bulmer, M. G. (1974). On fitting the Poisson lognormal distribution to species-abundance data. Biometrics, 30, 101–110.

Bürkner, P.-C. (2017). brms: An R package for Bayesian multilevel models using Stan. J. Stat. Software, 80(1), 1–28.

Carpenter, B., Gelman, A., Hoffman, M. D., Lee, D., Goodrich, B., Betancourt, M., Brubaker, M., Guo, J., Li, P., and Riddell, A. (2017). Stan: a probabilistic programming language. J. Stat. Software, 76(1), 1–32.

Clark, W. G. (2008). Effect of hook competition on survey CPUE. In Int. Pac. Halibut Comm. Report of Assessment and Research Activities 2007, pages 211–215.

Edwards, A. M., Haigh, R., and Starr, P. J. (2017). Redbanded Rockfish *(Sebastes babcocki)* stock assessment for the Pacific coast of Canada in 2014. DFO Can. Sci. Advis. Sec. Res. Doc., 2017/058, v + 182 p.

Edwards, A. M., Anderson, S. C., Keppel, E. A., and Grandin, C. (2022). gfiphc: Data Extraction and Analysis for Groundfish Data from the IPHC Longline Survey in BC. R package available at https://github.com/pbs-assess/gfiphc.

Efron, B. (1987). Better bootstrap confidence intervals. J. Am. Stat. Assoc., 82(397), 171–185.

Engen, S., Lande, R., Walla, T., and DeVries, P. J. (2002). Analyzing spatial structure of communities using the two-dimensional Poisson lognormal species abundance model. Am. Nat., 160(1), 60–73.

Fernö, A., Solemdal, P., and Tilseth, S. (1986). Field studies on the behaviour of whiting (Gadus merlangus L.) towards baited hooks. FiskDir. Skr. Ser. HavUnders., 18, 83–95.

Grimes, C. B., Able, K. W., and Turner, S. C. (1982). Direct observation from a submersible vessel of commercial longlines for tilefish. Trans. Am. Fish. Soc., 111(1), 94–98.

Grüss, A. and Thorson, J. T. (2019). Developing spatio-temporal models using multiple data types for evaluating population trends and habitat usage. ICES J. Mar. Sci., 76(6), 1748–1761.

Gulland, J. A. (1955). Estimation of growth and mortality in commercial fish populations. Fishery Invest., Lond. II, 18, 1–46.

Gunderson, D. R. (1993). Surveys of Fisheries Resources. John Wiley & Sons.

Hartwick, E. B., Ambrose, R. F., and Robinson, S. M. (1984). Den utilization and the movements of tagged Octopus dofleini. Mar. Freshwater Behav. Physiol., 11(2), 95–110.

High, W. L. (1980). Bait loss from halibut longline gear observed from a submersible. Mar. Fish. Rev., 42, 26–29.

Hilborn, R. and Walters, C. J. (2013). Quantitative Fisheries Stock Assessment: Choice, Dynamics and Uncertainty. Springer Science & Business Media.

IPHC (2022). International Pacific Halibut Commission Annual Report 2021, IPHC-2022-AR2021-R. International Pacific Halibut Commission.

Kristensen, K., Nielsen, A., Berg, C. W., Skaug, H., and Bell, B. M. (2016). TMB: automatic differentiation and Laplace approximation. J. Stat. Software, 70, 1–21.

Kuriyama, P. T., Branch, T. A., Hicks, A. C., Harms, J. H., and Hamel, O. S. (2019). Investigating three sources of bias in hook-and-line surveys: survey design, gear saturation, and multispecies interactions. Can. J. Fish. Aquat. Sci., 76(2), 192–207.

Lindgren, F. and Rue, H. (2015). Bayesian spatial modelling with R-INLA. J. Stat. Software, 63(19), 1–25.

Lindgren, F., Rue, H., and Lindström, J. (2011). An explicit link between Gaussian fields and Gaussian Markov random fields: the stochastic partial differential equation approach. J. R. Stat. Soc., Ser. B, 73(4), 423–498.

Løkkeborg, S. and Bjordal, Å. (1995). Size-selective effects of increasing bait size by using an inedible body on longline hooks. Fish. Res., 24(4), 273–279.

Løkkeborg, S., Olla, B., Pearson, W., and Davis, M. (1995). Behavioural responses of sablefish, Anoplopoma fimbria, to bait odour. J. Fish Biol., 46(1), 142–155.

Maronna, R. A., Martin, R. D., Yohai, V. J., and Salibián-Barrera, M. (2019). Robust Statistics: Theory and Methods (with R). John Wiley & Sons.

Obradovich, S. G. (2018). Evaluating key assumptions of a hook-based relative abundance index derived from the catch of bottom longlines. Ph.D. thesis, University of British Columbia, Canada.

R Core Team (2022). R: A Language and Environment for Statistical Computing. R Foundation for Statistical Computing, Vienna, Austria.

Ranganathan, P. and Pramesh, C. S. (2012). Censoring in survival analysis: potential for bias. Perspect. Clin. Res., 3(1), 40.

Ricker, W. E. (1975). Computation and interpretation of biological statistics of fish populations. Bull. Fish. Res. Bd. Can., 191, 1–382.

Rodgveller, C. J., Lunsford, C. R., and Fujioka, J. T. (2008). Evidence of hook competition in longline surveys. Fishery Bull., 106(4), 364–374.

Rodgveller, C. J., Sigler, M. F., Hanselman, D. H., and Ito, D. H. (2011). Sampling efficiency of longlines for Shortraker and Rougheye Rockfish using observations from a manned sub-mersible. Mar. Coast. Fish., 3(1), 1–9.

Rothschild, B. J. (1967). Competition for gear in a multiple-species fishery. ICES J. Mar. Sci., 31(1), 102–110.

Rue, H., Martino, S., and Chopin, N. (2009). Approximate Bayesian inference for latent Gaussian models by using integrated nested Laplace approximations. J. R. Stat. Soc., Ser. B, 71(2), 319–392.

Sigler, M. F. (2000). Abundance estimation and capture of sablefish (Anoplopoma fimbria) by longline gear. Can. J. Fish. Aquat. Sci., 57(6), 1270–1283.

Skud, B. E. and Hamley, J. M. (1978). Factors affecting longline catch and effort. International Pacific Halibut Commission.

Somerton, D. A. and Kikkawa, B. S. (1995). A stock survey technique using the time to capture individual fish on longlines. Can. J. Fish. Aquat. Sci., 52(2), 260–267.

Stoner, A. W. (2003). Hunger and light level alter response to bait by Pacific halibut: laboratory analysis of detection, location and attack. J. Fish Biol., 62(5), 1176–1193.

Stoner, A. W. (2004). Effects of environmental variables on fish feeding ecology: implications for the performance of baited fishing gear and stock assessment. J. Fish Biol., 65(6), 1445–1471.

Stoner, A. W., Ottmar, M. L., and Hurst, T. P. (2006). Temperature affects activity and feeding motivation in Pacific halibut: implications for bait-dependent fishing. Fish. Res., 81(2-3), 202–209.

Thorson, J. T., Shelton, A. O., Ward, E. J., and Skaug, H. J. (2015). Geostatistical delta-generalized linear mixed models improve precision for estimated abundance indices for west coast groundfishes. ICES J. Mar. Sci., 72(5), 1297–1310.

Thorson, J. T., Ianelli, J. N., Larsen, E. A., Ries, L., Scheuerell, M. D., Szuwalski, C., and Zipkin, E. F. (2016). Joint dynamic species distribution models: a tool for community ordination and spatio-temporal monitoring. Global Ecol. Biogeogr., 25(9), 1144–1158.

Ward, P., Myers, R. A., and Blanchard, W. (2004). Fish lost at sea: the effect of soak time on pelagic longline catches. Fishery Bull., 102(1), 179–195.

Watson, J., Joy, R., Tollit, D., Thornton, S. J., and Auger-Méthé, M. (2021). Estimating animal utilization distributions from multiple data types: A joint spatiotemporal point process framework. Annal. Appl. Stat., 15(4), 1872–1896.

Webster, R. A. and Stewart, I. J. (2013). Apportionment and regulatory area harvest calculations. In Int. Pac. Halibut Comm. Report of Assessment and Research Activities 2013, pages 187–205.

Webster, R. A., Hare, S. R., Valero, J. L., and Leaman, B. M. (2011). Notes on the IPHC set-line survey design, alternatives for estimating biomass distribution, and the hook competition adjustment. In Int. Pac. Halibut Comm. Report of Assessment and Research Activities 2010, pages 229–240.

Webster, R. A., Soderlund, E., Dykstra, C. L., and Stewart, I. J. (2020). Monitoring change in a dynamic environment: spatiotemporal modelling of calibrated data from different types of fisheries surveys of Pacific halibut. Canadian Journal of Fisheries and Aquatic Sciences, 77(8), 1421–1432.

Wood, S. N. (2004). Stable and efficient multiple smoothing parameter estimation for generalized additive models. J. Am. Stat. Assoc., 99(467), 673–686.

Wood, S. N. (2011). Fast stable restricted maximum likelihood and marginal likelihood estimation of semiparametric generalized linear models. J. R. Stat. Soc., Ser. B, 73(1), 3–36.

Wood, S. N., Pya, N., and Säfken, B. (2016). Smoothing parameter and model selection for general smooth models (with discussion). J. Am. Stat. Assoc., 111, 1548–1575.

Yamanaka, K. L., Obradovich, S. G., Cooke, K., Lacko, L. C., and Dykstra, C. (2008). Summary of non-halibut catch from the Standardized Stock Assessment Survey conducted by the International Pacific Halibut Commission in British Columbia from May 19 to July 22, 2006. Can. Tech. Rep. Fish. Aquat. Sci., 2796, vii + 58 p.

Yamanaka, K. L., McAllister, M. M., Etienne, M.-P., Edwards, A. M., and Haigh, R. (2018). Stock assessment for the outside population of Yelloweye Rockfish (*Sebastes ruberrimus*) for British Columbia, Canada in 2014. DFO Can. Sci. Advis. Sec. Res. Doc., 2018/001, ix + 150 p.

Zhang, Z. and Dunham, J. S. (2013). Construction of biological reference points for management of the Dungeness crab, Cancer magister, fishery in the Fraser River Delta, British Columbia, Canada. Fish. Res., 139, 18–27.

